# Tunable Cytosolic Chloride Indicators for Real-Time Chloride Imaging in Live Cells

**DOI:** 10.1101/2024.08.08.606814

**Authors:** Jared Morse, Maheshwara Reddy Nadiveedhi, Matthias Schmidt, Fung-Kit Tang, Colby Hladun, Prasanna Ganesh, Zhaozhu Qiu, Kaho Leung

## Abstract

Chloride plays a crucial role in various cellular functions, and its level is regulated by a variety of chloride transporters and channels. However, to date, we still lack the capability to image instantaneous ion flux through chloride channels at single-cell level. Here, we developed a series of cell-permeable, pH-independent, chloride-sensitive fluorophores for real-time cytosolic chloride imaging, which we call CytoCl dyes. We demonstrated the ability of CytoCl dyes to monitor cytosolic chloride and used it to uncover the rapid changes and transient events of halide flux, which cannot be captured by steady-state imaging. Finally, we successfully imaged the proton-activated chloride channel-mediated ion flux at single-cell level, which is, to our knowledge, the first real-time imaging of ion flux through a chloride channel in unmodified cells. By enabling the imaging of single-cell level ion influx through chloride channels and transporters, CytoCl dyes can expand our understanding of ion flux dynamics, which is critical for characterization and modulator screening of these membrane proteins. A conjugable version of CytoCl dyes was also developed for its customization across different applications.

## Introduction

As the most abundant anions in cells, chloride plays a crucial role in various cellular functions,^1–3^ including the regulation of cytoplasmic composition,^4^ cell volume,^5,6^ membrane potential,^7^ excitability,^8^ lysosome function,^9–14^ and innate immune response.^15,16^ The intracellular chloride concentration varies from organelle to organelle, spanning a range from 10 mM to 120 mM.^17–19^ It has been proposed that chloride homeostasis correlates with cell/organelle integrity. Dysregulation of chloride homeostasis is observed in neurodegenerative diseases^20,21^ such as lysosomal storage disorders.^12,13,22,23^ On the other hand, mutation of chloride channels can cause severe diseases such as cystic fibrosis,^24^ proteinuria,^25,26^ kidney stones,^27^ and osteoporosis.^28,29^ Additionally, chloride channels such as the proton-activated chloride (PAC) channel and volume-regulated anion channel^30^ have been recently reported to be involved in brain damage after ischemic stroke.^31,32^ Together, these studies highlight the importance of measuring intracellular chloride, which is also pivotal for discovering and characterizing new chloride channels, investigating the role of chloride and chloride channels in cellular processes, and aiding in diagnosis.

Fluorescence imaging allows us to directly visualize intracellular ion change, which is one of the most crucial methods for studying ion homeostasis and characterizing ion channels.^33^ However, the lack of tools for live-cell chloride imaging, particularly for real-time applications, is a significant impediment for chloride study. Due to the lack of effective imaging tools, recent studies still estimate the steady state cytosolic [Cl⁻] by measuring [Cl⁻] in cell lysates.^15,34–36^

Premo^TM^ YFP halide sensor is the only commercially available tool for chloride imaging. Other genetically encoded protein sensors have also been developed to monitor intracellular chloride changes.^17,37–44^ However, a notable limitation associated with these protein-based sensors is their susceptibility to variations in pH.^33^ Observed changes in fluorescence signal may result from fluctuations in pH levels rather than concentration changes of the targeted analyte. Even though there are pH-correctable, protein-based chloride sensors,^45,46^ their inherent pH sensitivity complicates experimental design and interpretation, making it challenging to ensure the fidelity and accuracy of the acquired data. Their pH-correctable design also limits their application in real-time imaging.

On the other hand, a few cell-impermeable, quinolinium-based and acridinium-based chloride-sensitive fluorophores have been developed^47–51^ over the past few decades.^33,52,19,53^ However, the quinolinium-based fluorophores need to be reduced for cell membrane penetration and re-oxidized in cytosol for chloride imaging.^19,52^ This indirect approach requires prior chemical steps before use, which severely hinders ease and practicality of use. The acridinium-based fluorophores such as 10,10′-bis(3-carboxypropyl)-9,9′-biacridinium (BAC)^54,55^ is also cell-impermeable and its application is restricted in lysosomes, as lysosome-labeling via endocytosis does not require the dye to be cell-permeable.

Due to the limitation of existing methods, we herein developed a series of cell-permeable, pH-independent, chloride-sensitive fluorophores for cytosolic chloride imaging (**1**‒**6**, Figure 1a), which we call CytoCl dyes. CytoCl dyes exhibit distinct emission spectra ranging from green to yellow and can detect chloride in a pH-independent manner. The ability of CytoCl dyes to monitor intracellular chloride was validated by in-cell chloride clamping and chloride channel modulator treatment. The high photostability of CytoCl dyes make it particularly suitable for real-time imaging in live cells. It could capture the instantaneous halide flux, which occurs within one second. With our new tools, we can capture rapid changes and transient events of halide flux, which cannot be captured by steady-state imaging. As a proof of concept, we demonstrated that our tools could image the real-time ion flux through the acid-sensitive PAC channel at single-cell level. To our knowledge, this is the first real-time cytosolic chloride imaging dyes, which is unique in enabling real-time imaging of instantaneous halide influx at single-cell level. CytoCl dye series are practical and versatile tools for cytosolic chloride imaging, valuable in chloride channel characterization and modulator screening, particularly in studying ion flux dynamics. A conjugable version of CytoCl dyes was also developed for its customization across different applications.

**Figure 1.**
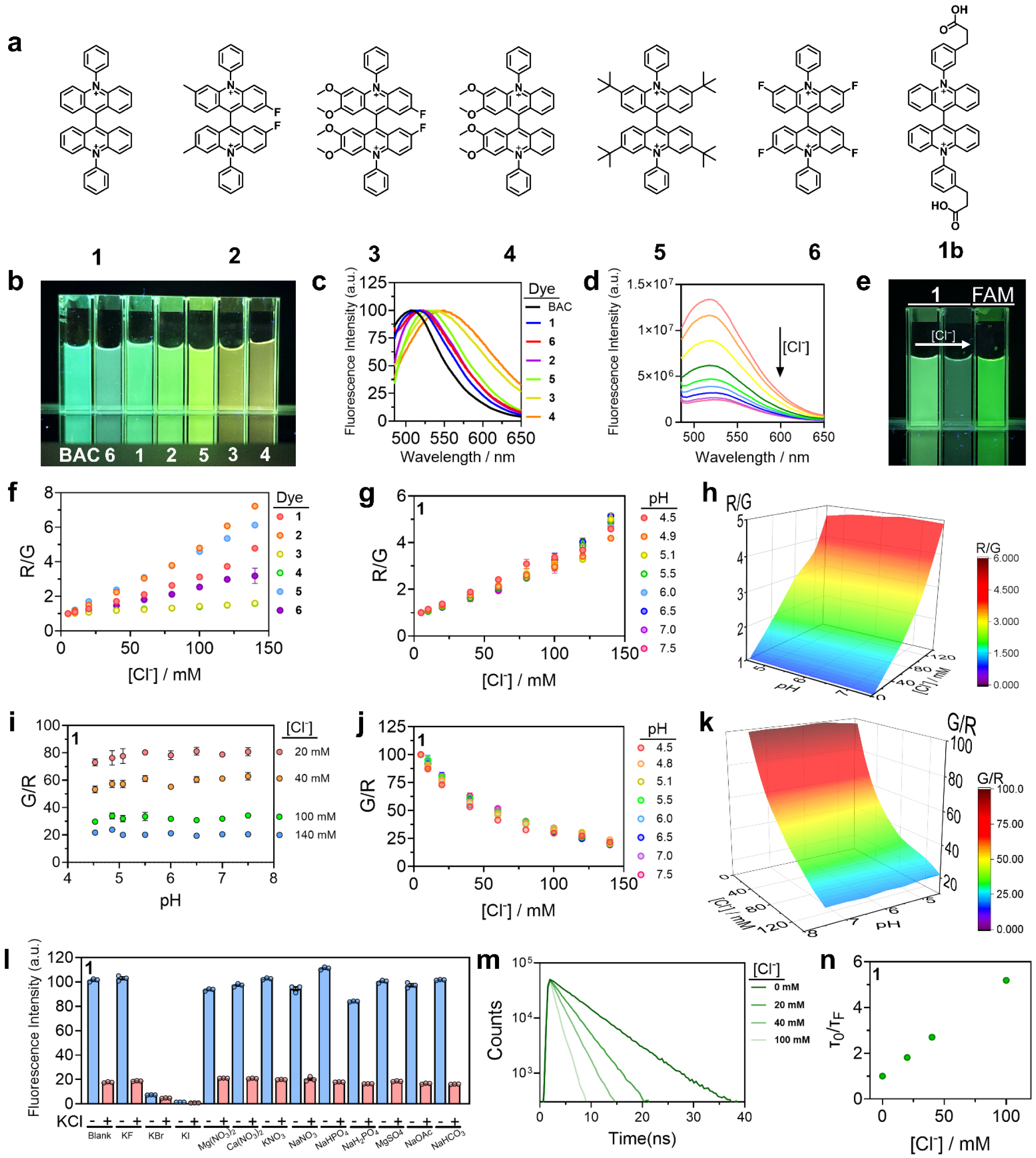
Design and *in vitro* characterization of CytoCl Dyes. **a**, Chemical structures of CytoCl dyes (**1**‒**6**, **1b**). **b**, **1**‒**6** in 5 mM UB4 (5 mM HEPES, MES and sodium acetate, 150 mM KNO_3_, 5 mM NaNO_3_, 1 mM Ca(NO_3_)_2_ and Mg(NO_3_)_2_) buffer at pH 7.0 under UV illumination. **c**, Normalized fluorescence emission spectra of **1**‒**6** in 5 mM UB4 at pH 7.0. **d**, Fluorescence emission spectra of **1** in 5 mM UB4 at pH 7.0 with increasing concentration of Cl^−^. **e**, **1** in 5 mM UB4 at pH 7.0 in the presence and absence of 100 mM potassium chloride under UV illumination. **f**, Normalized fluorescence intensity ratio (R/G) of Cyanine5 (R) and **1**‒**6** (G) as a function of Cl^−^ concentration. R/G values at different chloride concentrations were normalized to the value at 5 mM Cl^−^. **g**, Normalized R/G of **1** as a function of Cl^−^ concentration at the indicated pH value. **h**-**k**, Calibration surface plots of normalized R/G (**h**) and G/R (**k**) of **1** as a function of Cl^−^ and pH. **i**, Normalized G/R of **1** in the presence of 20, 40, 100, and 140 mM of chloride in 5 mM UB4 buffer with different pH. **j**, Normalized fluorescence intensity ratio (G/R) of **1** as a function of Cl^−^ concentration at the indicated pH value. G/R values at different chloride concentrations were normalized to the value at 5 mM Cl^−^. **l,** Selectivity profile of **1**. Fluorescence intensities of **1** in various anions (100 mM KF, KBr, KI, Mg(NO_3_)_2_, Ca(NO_3_)_2_, KNO_3_, NaNO_3_, NaHPO_4_, NaH_2_PO_4_, MgSO_4_, 5 mM NaOAc, 1 mM NaHCO_3_) with and without 100 mM KCl in 5 mM UB4 buffer at pH 7.0. **m,** Fluorescence lifetime decay of 20 μM **1** in 5 mM sodium phosphate buffer, pH 7.0, at various chloride concentrations **n,** Lifetime based Stern–Volmer plot of **1**. Error bars indicate the mean ± standard error of the mean (s.e.m.) of three independent measurements. Non-visible error bars represent errors too small to be seen.

## Results

### Design and *in vitro* Characterization of CytoCl Dyes

Existing chloride-sensitive fluorophores detect chloride through collisional quenching^56,57^ to achieve pH-independent chloride detection. This process necessitates a dye with a positively charged acridinium or quinolinium core, which cannot undergo protonation or deprotonation in physiological pH. These positively charged fluorophores are highly polar, resulting in low cell permeability. To overcome this limitation, we designed and synthesized a series of chloride-sensitive fluorophores with enhanced cell permeability, we called CytoCl dyes (Figure 1a). We hypothesize that optimizing the overall hydrophobicity of the fluorophore is critical for achieving enhanced cell permeability. To this end, we introduced phenyl groups into the cell-impermeable BAC to enhance its hydrophobicity (**1**). This modification makes the dyes cell-permeable, without affecting the acridinium core, which is critical for collisional quenching. The incorporation of phenyl group also enables an extended π system of the fluorophore, achieving a bathochromic shift. Additionally, we designed and synthesized compounds **2**‒**6**, incorporating different electron-donating and electron-withdrawing groups to fine-tune the emission wavelength and chloride sensitivity of the fluorophore.

Replacing the carboxyl group in BAC with a phenyl group (**1**) resulted in a shift of the maximum emission from 505 nm to 518 nm (Figure 1b‒c). The introduction of electron-donating groups, such as methoxy (**4**) and *t*-butyl (**5**), into the acridinium core caused a red shift in emission. We then modified the structure with electron-withdrawing groups, such as fluoride. This countered the effects of electron-donating groups resulting in a relative blue shift in emission (from **4** to **3**) (Figure 1b‒c). Under UV illumination, the fluorescence of compounds **1**‒**6** was quenched upon the addition of potassium chloride (Figure 1e, Supplementary Figure 1). The fluorescence response characteristics of **1**‒**6** was then investigated as a function of pH and [Cl^−^] in order to determine their pH and [Cl^−^] sensitive regimes (Figure 1f‒k, Supplementary Figure 3‒7). We investigated the response of **1**‒**6** by examining the fluorescence intensity ratio (G/R) between each compound (**1**‒**6,** G) and Cyanine5 (R) in the varying environments (working scheme is depicted in Supplementary Figure 3). The G/R ratio of **1** decreased exponentially as a function of [Cl^−^] (Figure 1j, Supplementary Figure 4). Subsequently, the response of **1**‒**6** was analyzed by plotting R/G against [Cl^−^] (Stern-Volmer plot), which showed a linear relationship (Figure 1g, Supplementary Figure 5).

To further validate the pH-independence, the fluorescence intensity of **1** in various pH was investigated (Supplementary Figure 6). No significant variation was seen across physiologically relevant pH 4.5‒7.0. The G/R ratio of **1** in the presence of 20, 40, 100, and 140 mM of chloride in pH 4.5−7.5 was shown in Figure 1i and Supplementary Figure 7. Notably, the fold change in G/R upon addition of chloride remained invariant for **1** across a pH range of 4.5−7.5. The Stern–Volmer constant (*K*sv) in the R/G plot stayed constant as a function of pH from pH 4.5‒7.5 (Figure 1g). These results can also be illustrated by 3D surface plots of R/G (Figure 1h) and G/R (Figure 1k) as a function of [Cl^−^] along the pH range from 4.5 to 7.5. These results demonstrated that **1**‒**6** detect physiological chloride range from 10 mM to 120 mM in a pH-independent manner.

The effects of physiological anions on **1**‒**6** were also investigated (Figure 1l, Supplementary Figure 8). Minor fluorescence quenching was observed in the presence of highly excess anions such as fluoride, nitrate, phosphate, and sulfate, as well as physiological levels of carbonate and acetate.^58^ We observed that halide ions such as bromide and iodide also quenched the fluorescence of **1**‒**6**. However, the combined concentrations of all these ions in cells are < 0.1% the value of chloride.^59–66^ At this low bioavailability, their contribution to the fluorescence changes of **1**‒**6** in biological systems is negligible. Additionally, the fluorescence lifetime of **1** decreased dose-dependently in the presence of increasing concentrations of chloride (Figure 1m-n), demonstrating its potential for fluorescence lifetime-based detection. The cytotoxicity of these dyes was also evaluated in HDF cells. With the labeling protocol for imaging, **1**−**4** and **6** exhibited no significant change in cell viability in HDF cells. A notable decrease in cell viability was observed when **5** was incubated at a concentration of 25 µM (Supplementary Figure 9). Thus, the labeling protocol revealed no significant impact on cell viability for compounds **1**−**6**.

### Cellular Uptake of CytoCl Dyes

Next, we investigated the cellular uptake and distribution of **1**‒**6**. Cells were incubated with 100 µM of **1**‒**6** at 37°C for 1 h, washed with PBS, and then incubated in complete cell culture medium for 30 min (referred to as 1-hour ‘pulse’ and 30-minute ‘chase’). After labeling, **1**, **2**, and **5** labeled cells exhibited high fluorescence signal (Supplementary Figure 10) but no fluorescence was observed for cells labeled with **3**, **4**, and **6** (Supplementary Figure 11). This may be due to the low brightness of these compounds that was observed in *in vitro* studies (Supplementary Figure 1). To further examine the cellular distribution of **1**, we co-stained the cells with **1** and organelle markers such as MitoView, LipidSpot, and LysoTracker (Supplementary Figure 12-13). The images of **1** did not specifically overlap with specific organelle marker. Instead, **1** stained the entire cell excluding the nucleus, making it suitable for cytosolic chloride imaging.

### Dose-Dependent Response of CytoCl Dyes to Intracellular Halide

Chloride channels typically exhibit a limited ability to effectively discriminate between chloride and other halides.^1,67,68^ Iodide stimulation is commonly used to assess whether a chloride channel is open during chloride channel characterization.^69^ Given that **1**‒**6** also exhibit sensitivity to iodide *in vitro*, we investigated the intracellular halide sensing ability of compounds **1**‒**6**. Upon iodide stimulation, fluorescence intensity of the labeled HDF was significantly attenuated (Figure 2a, Supplementary Figure 10). To illustrate the versatility of these dyes across various cell types, we extended our analysis to include RAW 264.7 macrophages, HEK-293 human embryonic kidney cells, and HeLa cells, which represent distinct cell types (Figure 2a, Supplementary Figure 14−16).

**Figure 2.**
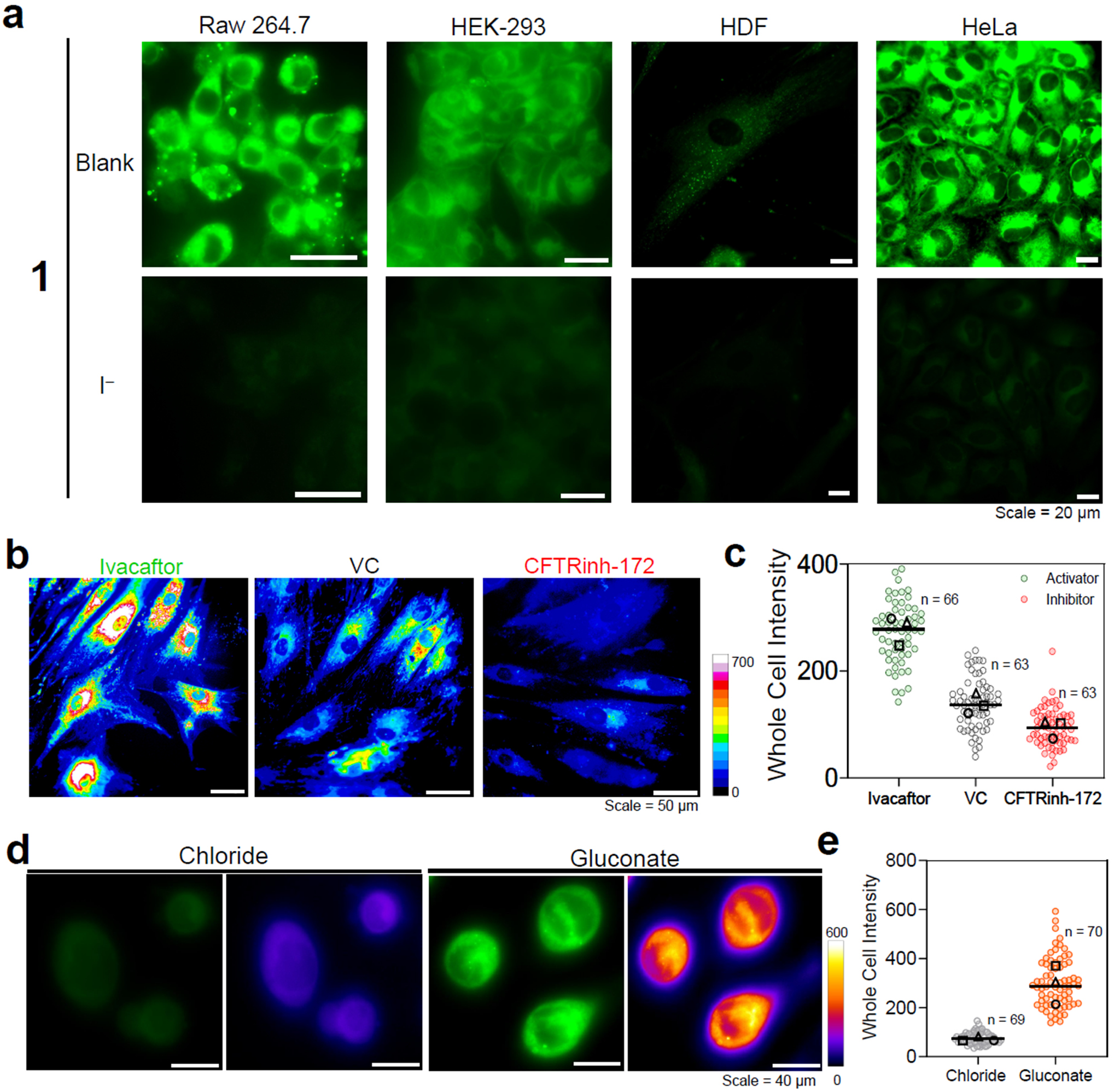
CytoCl Dyes Monitor the Intracellular Halide. **a,** Representative fluorescence images of **1**-labeled cells, with and without incubation of 75 mM iodide. **b**-**c.** CytoCl monitors the cytosolic chloride change induced by CFTR modulator. **b**, Representative fluorescence images of **1**-labeled HDF cells pre-treated with vehicle control (DMSO), 2 µM ivacaftor (CFTR activator), and 20 µM CFTRinh-172 (CFTR inhibitor). **c**, Quantification of whole cell intensity at each condition. **d**-**e,** CytoCl monitors the cytosolic chloride change induced by chloride substitution. **d**, Representative fluorescence images of **1**-labeled HeLa cells incubated with chloride-containing and chloride-free buffer. **e**, Quantification of whole cell intensity of **1**-labeled cells at each condition. Experiments were performed in triplicate. The median value of each trial is given by a square, circle and triangle symbol (n = number of cells).

We then stimulated the labeled cells with different [l^−^]. The fluorescence intensity of the **1**-labeled cells responded to [l^−^] in a dose-dependent manner (Supplementary Figure 17). Supplementary figure S17a presents representative fluorescence images of cells stimulated with different [l^−^]. Histograms depicting the whole cell fluorescence intensity across different [l^−^] are shown in Supplementary Figure 17b-c. Similar results were observed for **2** and **5** (Supplementary Figures 18−19). These results demonstrate that **1**, **2**, and **5** can detect cytosolic halide change in live cells.

### CytoCl Dyes Monitor Changes in Cytosolic Chloride

Next, we sought to investigate the in-cell chloride sensing characteristics of **1**, **2**, and **5**. We clamped the cells with the indicated [Cl^−^]. We fixed the labeled HDF cells and incubated them with a clamping buffer containing the specified [Cl^−^], along with monensin, nigericin, and a squaramide-based synthetic chloride transporter (Supplementary Figure 20).^13,70,71^ The whole-cell fluorescence intensity decreased with increasing concentrations of clamped chloride. To determine whether CytoCl dyes can monitor cytosolic chloride changes in live cells, we activated and inhibited cystic fibrosis transmembrane conductance regulator (CFTR) by ivacaftor (a CFTR activator)^72^ and CFTRinh-172 (a CFTR inhibitor).^73^ CFTR is a transmembrane protein that exports chloride across the cell membrane. Upon incubation with the CFTRinh-172, the cells exhibited a notably diminished fluorescence intensity compared to the control. Conversely, cells treated with the ivacaftor displayed a markedly elevated fluorescence intensity (Figure 2b-c, Supplementary Figure 21). Furthermore, the substitution of chloride in the extracellular cell culture medium has been widely used to deplete the intracellular chloride when studying chloride physiology.^15,74,75^

We then applied **1** to monitor the cytosolic chloride following chloride substitution. **1**-labeled cells upon chloride substitution displayed significantly higher fluorescence intensity than the control sample (Figure 2d-e), showing a low cytosolic chloride level following chloride substitution. These results demonstrate that our tools can monitor intracellular chloride changes in live cells.

### CytoCl Dyes Capture Single-Cell Level Halide Flux in Real-Time

Fluorescence imaging constitutes a crucial method for directly visualizing ion channel dynamics under physiological conditions. Therefore, we sought to investigate the capability of **1** for imaging real-time halide influx in live cells. In Figure 2a, we observed the halide uptake upon iodide stimulation in HeLa, HEK-293, RAW 264.7 macrophages, and primary human fibroblasts through steady-state imaging (Supplementary Figures 14−15). We then monitored the iodide uptake processes by real-time imaging (Figure 3a). The fluorescence intensity of the labeled cells decreased rapidly upon iodide stimulation, with the fluorescence being quenched in less than half a second. In contrast, there was no significant change in the fluorescence intensity of the control sample, indicating that the decrease in fluorescence intensity is not due to photobleaching (Figure 3a, and video 1).

**Figure 3.**
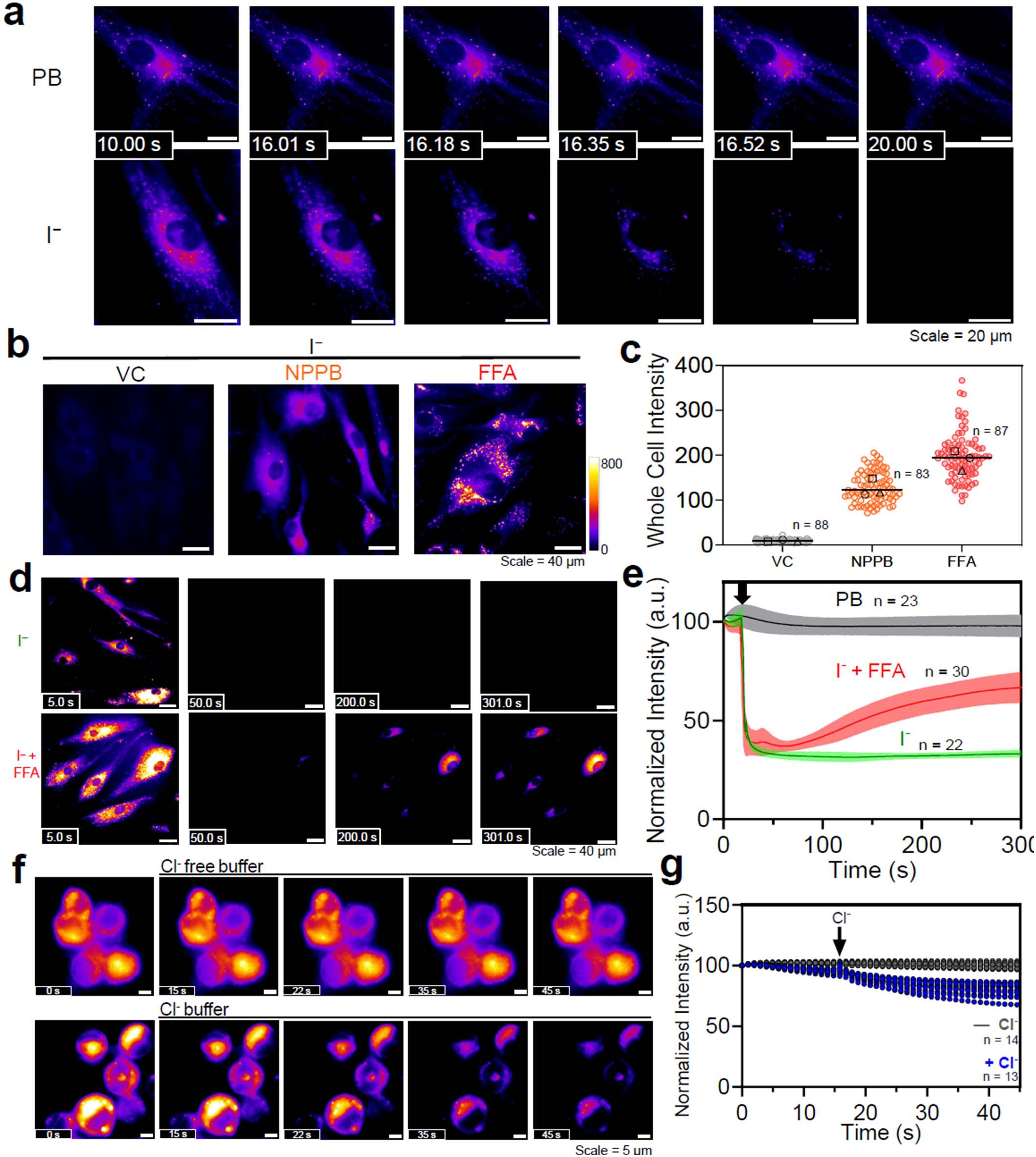
CytoCl Dyes Capture the Rapid Changes and Transient Events of Halide Flux in Real Time. **a, 1**-labeled HDF cells were subjected to iodide stimulation. Representative fluorescence images of **1**-labeled HDF cells at different time points. Images were recorded at a frame rate of 6 fps. PB, photobleaching. **b**-**e**, Visualizing the impact of chloride channel blockers on iodide import using (**b**-**c**) steady state imaging and (**d**-**e**) real time imaging. **b**-**c**, Steady state imaging. Primary HDF cells pre-treated with vehicle control (DMSO), 100 µM NPPB, and 150 µM FFA, were labeled with **1** and then subjected to iodide stimulation for 1 hour. **b**, Representative fluorescence images of **1**-labeled HDF cells. **c**, Quantification of whole cell intensity of **1**-labeled cells at each condition. Experiments were performed in triplicate. The median value of each trial is given by a square, circle and triangle symbol (n = number of cells). **d**-**e**, Real time imaging. Primary HDF cells pre-treated with vehicle control (DMSO) and 150 µM FFA, were labeled with **1** and then subjected to iodide stimulation. The images were recorded at a frame rate of 1 fps. **d**, Representative fluorescence images of **1**-labeled HDF cells at different time points. **e**, Normalized whole cell intensity of **1**-labeled cells at each condition. The dark arrow indicates the time of iodide stimulation. Error shade indicates the median ± standard error of the mean (s.e.m.) of three independent measurements (n = number of cells). **f**-**g**, To induce chloride influx, chloride-depleted **1**-labeled HeLa cells were incubated with chloride-containing and chloride-free buffer. **f**, Representative fluorescence images of **1**-labeled HeLa cells at different time points. **g**, Normalized whole cell intensity of **1**-labeled cells at each condition. The dark arrow indicates the time of stimulation. The images were recorded at a frame rate of 1 fps. Experiments were performed in triplicate (n = number of cells).

Next, we sought to evaluate the impact of chloride channel blockers on the iodide uptake. Through the steady-state imaging, we observed that the cells exhibited weak fluorescence due to the uptake of iodide (Figure 3b-c, Supplementary Figures 22). On the other hand, cells pretreated with chloride channel blockers (NPPB and FFA) displayed relatively high fluorescence, which indicates less intracellular iodide. Based on the results of steady-state imaging, we can hypothesize that chloride channel blockers blocked the import of iodide into the cells. However, a different conclusion could arise if we monitor this ion influx through real-time imaging. The fluorescence of **1**-labeled cells was significantly quenched upon iodide stimulation. However, in cells pretreated with chloride channel blockers, upon iodide stimulation, the fluorescence intensity was initially quenched significantly, then partially recovered within the 5-minute period following iodide stimulation (Figure 3d-e and video 2). This indicates that, with the treatment of chloride channel blockers, iodide was still initially imported into the cells and then partially exported out of the cells. This phenomenon can only be observed using real-time imaging. These results demonstrate that CytoCl dyes can capture rapid changes and transient events by real-time imaging, which provides the temporal information for studying ion flux dynamics.

Additionally, we also utilized our tool to visualize the instantaneous chloride influx in live cells. The influx of chloride was triggered by treating the chloride-depleted cells with chloride-rich medium. Upon treatment, the fluorescence of the labeled cell gradually diminished over time which corresponds to the chloride influx (Figure 3f-g, Video 3). This result underscores that our tool is unique in enabling real-time imaging of chloride influx.

### CytoCl Dyes Capture PAC Channel-Mediated Instantaneous Ion Influx at Single-Cell Level

Next, we sought to further demonstrate the capability of **1** to capture the ion influx mediated by a specific chloride channel in real-time. Here, we utilized our tools to monitor acid-induced ion flux in live cells. The PAC channel is a ubiquitously expressed, pH-sensing ion channel encoded by *PACC1* (*TMEM206*).^76^ It is found on both endosomes and cell surfaces, where it becomes activated under pathological conditions related to acidosis. To prevent the passive redistribution of iodide observed in Figure 3d, we reduced the [I^‒^] for stimulation to 10 mM. In this scenario, no significant fluorescence quenching of **1**-labeled cells was observed upon stimulation of 10 mM iodide at pH 7.0. We subsequently lowered the pH to initiate activation of the PAC channel,^31,32,77^ which induced an influx of I^‒^, leading to a rapid decrease in fluorescence of **1**-labeled WT HeLa cells. The extent of ion influx increased as the pH decreased from 7.0 to 4.5 (Figure 4a-b, and video 4). To validate that the observed ion influx occurs through the PAC channel, we conducted the same experiment using PAC-KO HeLa cells. No significant fluorescence quenching was observed between pH 4.5 and pH 7.0 with PAC-KO cells (Figure 4a-b, Supplementary Figure 24, and video 5). It indicates that the proton-triggered iodide influx is mediated by the PAC channel. These results demonstrate that our tool is capable of imaging real-time, instantaneous ion influx through specific chloride channels, such as PAC.

**Figure 4.**
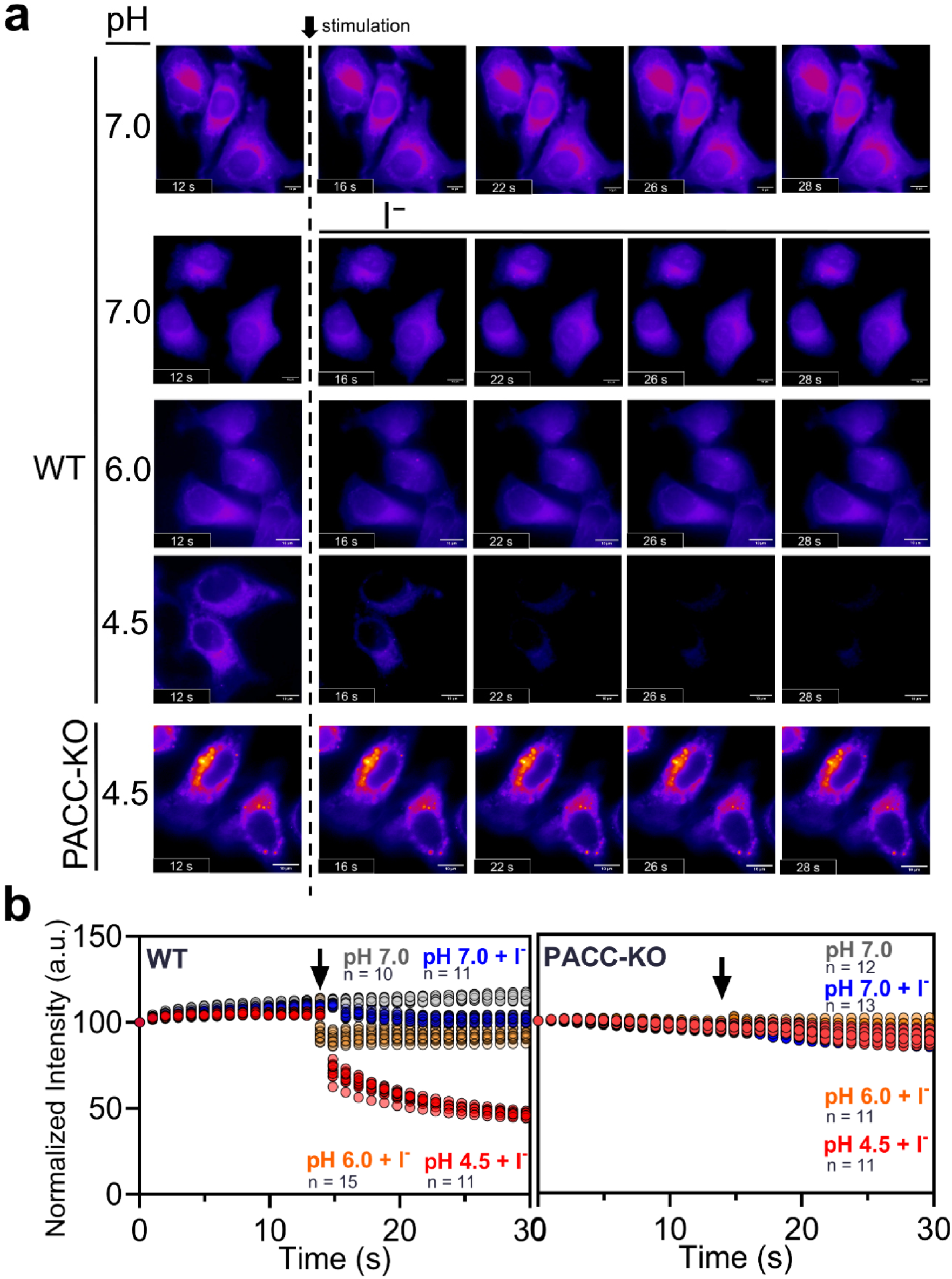
CytoCl Dyes Visualize Ion Flux Through Proton-Activated Chloride (PAC) Channel at Single-Cell Level. **a**-**b, 1**-labeled WT / PAC-KO HeLa cells were stimulated with I^−^ containing buffer at various pH. **a**, Representative fluorescence images of **1**-labeled HeLa cells at different time points. **b**, Normalized whole cell intensity of **1**-labeled cells at different time points. The images were recorded at a frame rate of 1 fps. Experiments were performed in triplicate (n = number of cells).

## Discussion

Fluorescence imaging is one of the most crucial techniques for directly visualizing intracellular ions. The primary methods for cytosolic chloride imaging include the use of small molecules and genetically encodable chloride indicators. However, small molecule-based indicators have limited cell permeability that required pre-reduction of the dye to penetrate the plasma membrane. Genetically encodable chloride indicators are pH-sensitive, so the observed signal changes may be attributed to fluctuations in either pH or chloride levels. Our chloride-sensitive fluorophores, CytoCl dyes, exhibit exceptional cell permeability and are pH-insensitive. These advanced CytoCl dyes serve as effective cytosolic chloride indicators capable of directly visualizing real-time cytosolic chloride change in a pH-independent manner.

Using our tools in real-time imaging, we successfully captured rapid and transient events that are unattainable by steady-state imaging. In Figure 3d, following iodide stimulation in chloride channel blocker-treated cells, we observed an instantaneous iodide influx and subsequent iodide efflux prior to reaching the steady state. This emphasizes the significance of studying ion flux dynamics while our tools are unique in enabling real-time imaging of instantaneous halide flux at single-cell level. We also successfully imaged the acid-induced ion flux through the PAC channel, at single cell level. The PAC channel is an emerging therapeutic target for acidosis^76,78–80^ but the pharmacology of the PAC channel remains poorly understood.^31^ Our new tools will provide a straightforward and rapid method to screen the channel modulators and investigate the PAC channel modulation under biologically relevant conditions. It will also contribute to the discovery of new chloride channels and the exploration of chloride physiology.

We are aware that multi-fluorophore-based reporters have been developed to quantify steady-state lysosomal chloride through multispectral imaging.^10,13,81^ In such cases, the concentration difference of the sensor between organelles/cells could be neglected. In this study, we demonstrate that the fluorescence lifetime of CytoCl responds to changes in chloride levels, showing its potential for fluorescence lifetime imaging. In this method, the lifetime output is independent of sensor concentration, which enables the quantitative measurement of chloride.^82^ We also developed a synthetic pathway to synthesize the conjugable version of CytoCl dyes, allowing for customization across different applications such as constructing a 2-dye based reporter.

In summary, CytoCl dyes are a series of cell-permeable, pH-independent cytosolic chloride indicators with tunable emissions ranging from green to yellow. CytoCl dyes show good photostability, making it suitable for real-time imaging. It could capture instantaneous iodide (Figure 3a-e) and chloride flux (Figure 3f-g) in unmodified cells at single-cell level, a capability that was previously unattainable. With our new tools, we can capture the rapid changes and transient events (Figure 3d-e) by real-time imaging, which cannot be captured by steady-state imaging (Figure 3b-c). As a proof of concept, we also demonstrated that it could image the PAC channel-mediated ion flux in real-time at single-cell level (Figure 4). CytoCl dyes are practical and beneficial for various applications in chloride studies, including chloride channel characterization and modulator screening, and are particularly valuable for studying ion flux dynamics. A conjugable version of CytoCl dyes has also been developed, allowing for customizable applications and immediate practical use by the community.

## Supporting information

supporting information_synthesis

## Resources availability

We are delighted to provide our developed tools upon request.

## Acknowledgements

This work was supported by NIH grants R35GM147112 (K.L.), R35GM147112-02S2 (K.L.), R35GM147112-02S1 (K.L. & M.S.), R35GM124824 (Z.Q.), and Clarkson University start-up fund. We thank Patrick Lutz from St. Lawrence University for providing NMR support. We extend our sincere gratitude to St. Lawrence University for generously providing access to their JEOL 400 MHz nuclear magnetic resonance (NMR) spectrometer. The authors acknowledge the use of facilities and instrumentation (Edinburgh FLS1000 spectrometer) supported by NSF through the Cornell University Materials Research Science and Engineering Center DMR-1719875.

## Author contributions

K.L. and J.M. wrote the manuscript. All authors discussed the results and commented on the manuscript. J.M. and M.R.N. wrote the supporting information. CytoCl dyes were synthesized by M.R.N. and M.S. Cytotoxicity assays, cellular uptake, cellular distribution, intracellular iodide sensing characteristics, intracellular chloride calibration, steady-state cellular chloride imaging (CFTR and chloride substitution), and real-time halide influx imaging (chloride substitution and PAC) were performed by J.M. The photo-physical properties of CytoCl dyes were investigated by J.M., F.T. The *in vitro* calibrations were performed by J.M., and P.G. Fluorescence lifetime measurements were performed by J.M., C.H., and P.G. The selectivity assays were performed by F.T. and C.H.. Z. Q. provided the consultation and support for the PAC ion flux imaging.

