## supporting information_synthesis for "Tunable Cytosolic Chloride Indicators for Real-Time Chloride Imaging in Live Cells"

#### General synthetic materials

Compound **I**, **a1**, **e1** and 3-(3-Bromophenyl)propanoic acid was purchased from Ambeed (IL, US). Unless otherwise specified, all reagents and solvents were obtained from commercial suppliers and used without further purification. All reagents were weighed and handled in air at room temperature. $^{1}$H NMR spectra were recorded at 400 MHz and $^{13}$C NMR spectra were recorded at 100 MHz by using a Bruker Avance 400 MHz spectrometer (Department of Chemistry and Biomolecular Science, Clarkson University, NY) and JOEL 400 MHz spectrometer Department of Chemistry, St. Lawrence University, NY). Chemical shifts were calibrated using residual undeuterated solvent as an internal reference ($^{1}$H NMR: $CDCl_3$ 7.26 ppm, $^{13}$C NMR: $CDCl_3$ 77.0 ppm, $^{1}$H NMR: DMSO-$d_6$ 2.5 ppm, $^{13}$C NMR: DMSO-$d_6$ 45 ppm). NMR data were processed using MestReNova Software. The following abbreviations were used to describe peak splitting patterns when appropriate: s = singlet, d = doublet, t = triplet, q = quartet, m = multiplet, brs = broad singlet. The progress of the reaction was followed with TLC using silica gel SILG/UV 254 and 365 plates. High-resolution mass spectra (HRMS) were recorded using a SCIEX X500B QTOF mass spectrometer (at Center for Air and Aquatic Resources Engineering and Sciences, CAARES, Clarkson university, NY) which operated in positive ion mode (+ve ESI).

#### General Procedure for the Synthesis of 2-(phenylamino)benzoic acid derivatives: (GP1)

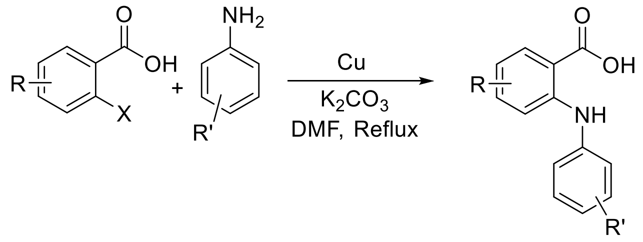


Phenyl aminobenzoic acids were synthesized using modified reported literature.[1] A mixture of 2-iodo/bromo benzoic acid derivatives (1 Equiv.), aniline derivatives (1.5 equiv.), $K_2CO_3$ (2 equiv.) and copper powder (25 mol%) were stirred at reflux conditions in DMF for 24 hours. The progress of the reaction was monitored by thin-layer chromatography. After completion of the reaction, water was added, and the precipitated product was collected and filtered, dried on a vacuum for overnight, and used in the next step without further purification.

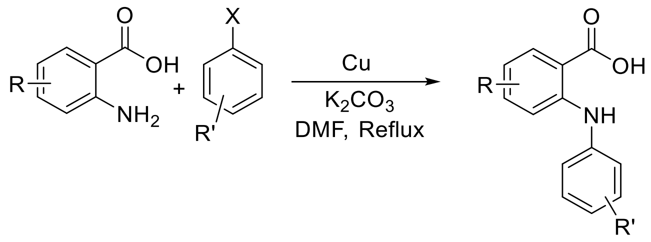


X= Br/I

A mixture of 2-aminobenzoic acid derivatives (1 Equiv.), aryl halide derivatives (1.5 Equiv.), $K_2CO_3$ (2 Equiv.) and copper powder (25 mol%) were stirred at reflux conditions in DMF for 24 hours. The progress of the reaction was monitored by thin-layer chromatography. After completion of the reaction, water was added, and the precipitated product was collected and filtered, dried on a vacuum for overnight, and used in the next step without further purification.

**General Procedure for the Synthesis of Acridones: (GP2)**

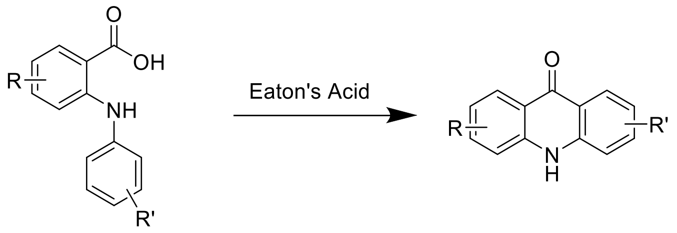


A mixture of 2-(phenylamino)benzoic acid derivatives (1 Equiv.) and Eaton's reagent (2 Equiv.) were stirred at 80 °C for 1-2 hours. The progress of the reaction was monitored by thin-layer chromatography. After completion of the reaction, the residue was transferred to ice-cold water, stirred for 10 minutes, and filtered. The obtained residue for the compounds was used as it is for the next step, for the compounds, purified with column chromatography from n-hexane/EtOAc (5:1) to afford the compounds **b1**, **c1**, **d1** and **e1** in 66-88%.

**General Procedure for N-Phenylation: (GP3)**

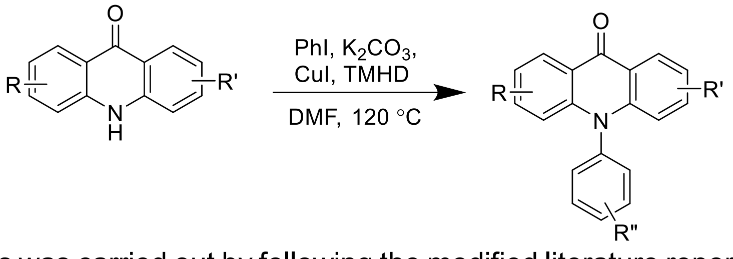


Phenylation of Acridones was carried out by following the modified literature report.[2] A mixture of acridone derivatives (1 Equiv.), Iodobenzene (1.2 Equiv.), $K_2CO_3$ (2 Equiv.), CuI (25 mol%), and TMHD (25 mol%) in DMF were stirred at 150 °C for 20 hours. The progress of the reaction was monitored by thin-layer chromatography. After completion of the reaction, diluted with ethyl acetate, filtered through celite, and the filtrate was washed with 1N HCl, the residue was purified with column chromatography from n-hexane/EtOAc (9:1) to afford the compounds **a2–f2** and **IIb** in 40-50%.

**General Procedure for Ketone self-coupling followed by oxidation: (GP4)**

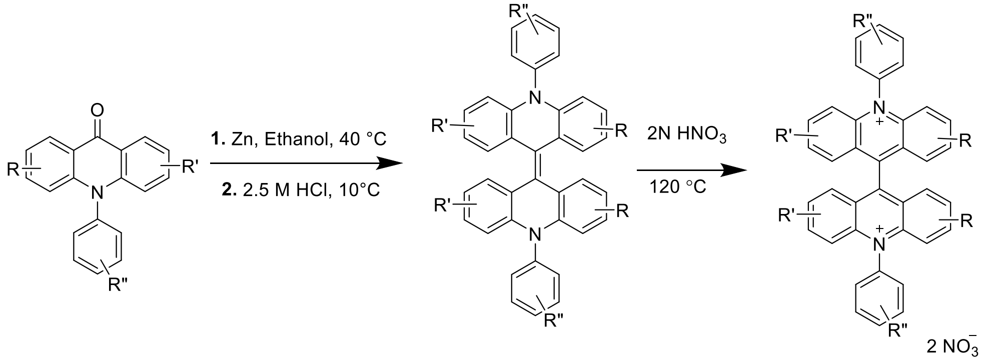


Dimerization followed by oxidation of acridone derivatves was carried out by following modified Literature.[3] Acridone derivatives were dissolved in ethanol and Zn (10 Equiv.) was added and stirred for 20 min at 40 ℃. Then cooled to 10 ℃ and 2.5 M HCl in ethanol was added dropwise, continued stirring at room temperature for a further 1 hour, and reaction progress was monitored by TLC. After completion of the reaction, diluted with 10 Equiv. of water and filtered the precipitate. To the crude alkene derivative, add 2N $HNO_3$ (10 Volumes) and stir at 120 ℃, for 2 hours. Then cooled to room temperature and filtered. The obtained precipitate was dried under a vacuum to obtain the compounds **1−6** and **1b.**

**Synthesis of BAC:**

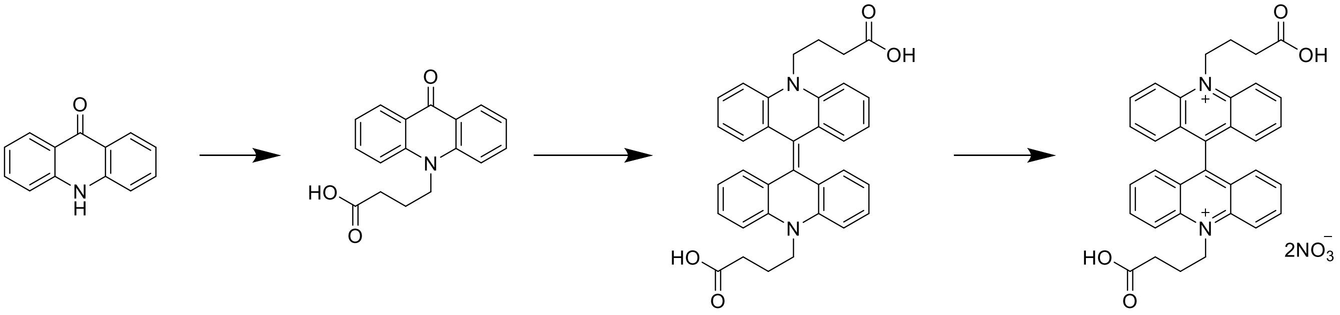

(I) (II) (III) BAC

**Scheme 1.** Synthesis of BAC.

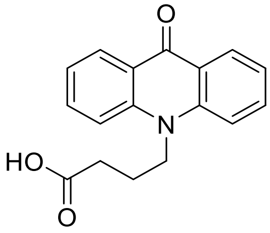

(II)

4-(9-oxoacridin-10(9H)-yl)butanoic acid (**II**)

Compound **II** was synthesized by using the reported literature procedure.[4,5] $^1$H NMR (400 MHz, $CDCl_3$) δ 12.34 (s, 1H), 8.36 (d, *J* = 7.9 Hz, 2H), 8.01 – 7.76 (m, 4H), 7.45 – 7.25 (m, 2H), 4.50 (d, *J* = 7.0 Hz, 2H), 2.64 – 2.56 (m, 2H), 2.04 (s, 2H). HRMS (+ TOF MS) calculated for $C_{17}H_{16}NO_3$ $[M + H]^+$, 282.1052; found: 282.1121

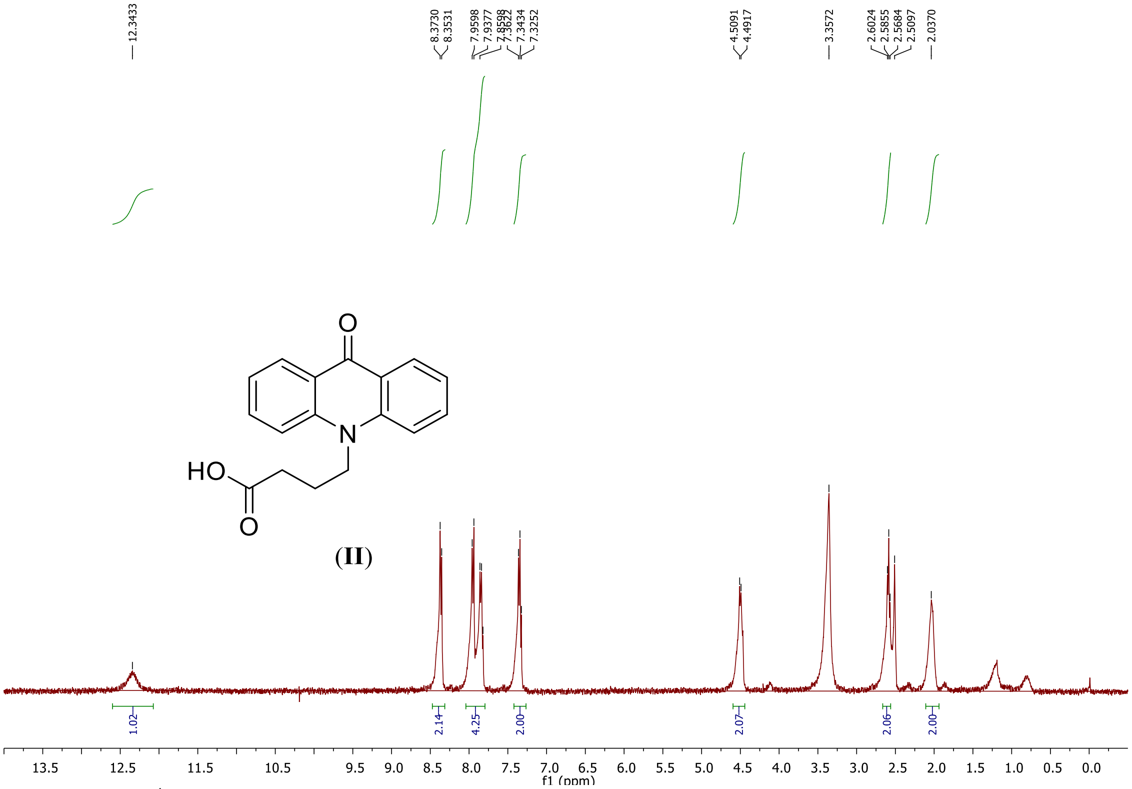

**Figure S1.** $^1$H NMR of compound **II**

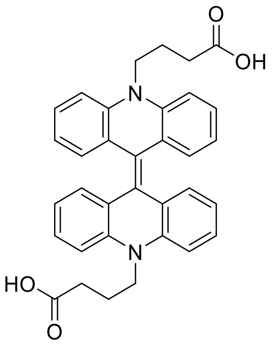

(III)

4,4'-(10H,10'H-[9,9'-biacridinylidene]-10,10'-diyl)dibutyric acid (**III**)
**III** was synthesized by using the reported literature procedure.[4,5] HRMS (+ TOF MS) calculated for $C_{17}H_{15}NO_2$, m/z, 282.1052 ; found: 282.1121

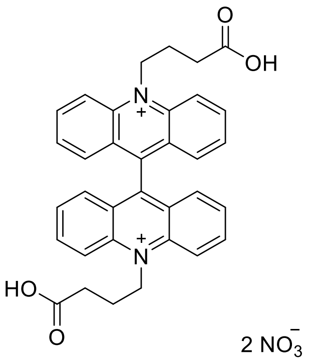

**BAC**

10,10'-bis(3-carboxypropyl)-[9,9'-biacridine]-10,10'-diium (**BAC**)
BAC was synthesized by using the reported literature procedure.[4,5] $^1$H NMR (400 MHz, DMSO-$d_6$) δ 12.48 (s, 1H), 9.10 (d, *J* = 9.5 Hz, 2H), 8.66 – 8.33 (m, 3H), 7.92 – 7.52 (m, 4H), 5.65 (d, *J* = 5.9 Hz, 3H), 3.03 – 2.80 (m, 2H), 2.53 (s, 2H). $^{13}$C NMR (101 MHz, $CDCl_3$) δ 174.74, 152.04, 141.64, 139.69, 129.73, 129.32, 127.26, 119.70, 50.95, 30.74, 24.48. HRMS $[M-3H]^-$ (- TOF MS) calculated for $C_{34}H_{30}N_2O_4$ 527.2195 found: 527.1989

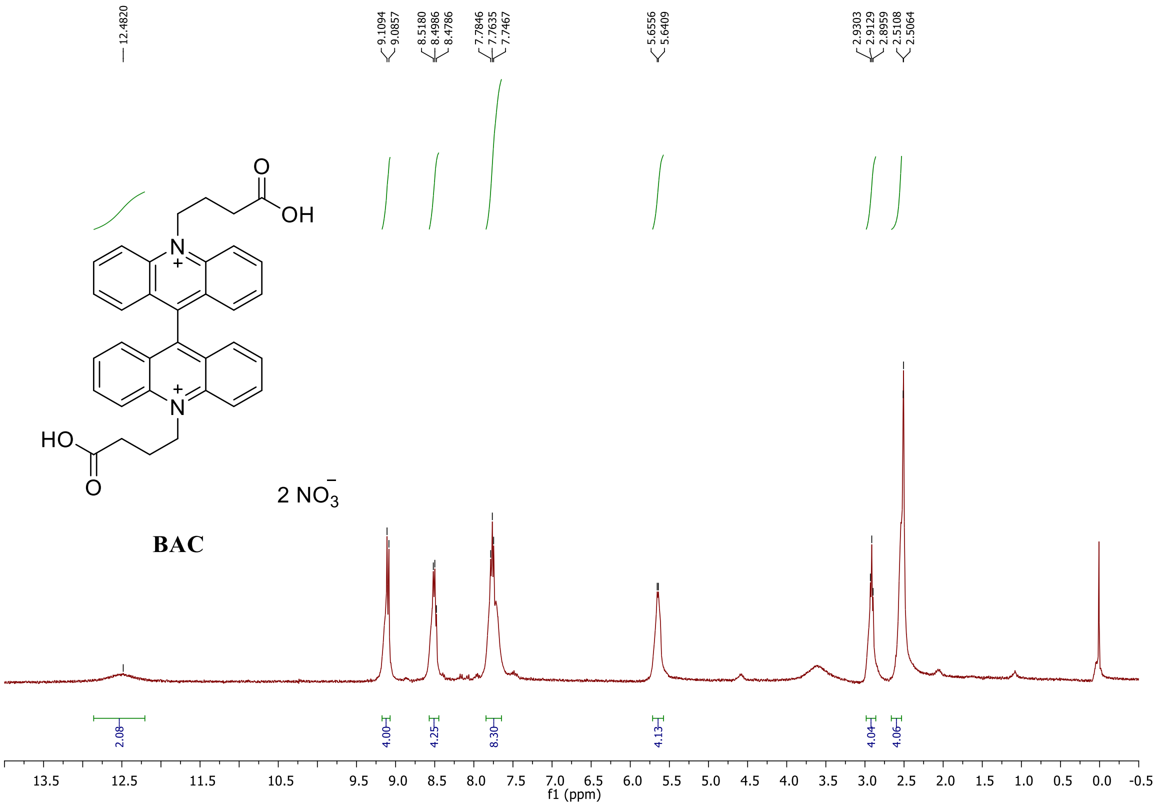


**Figure S2.** $^{1}H$ NMR of BAC.

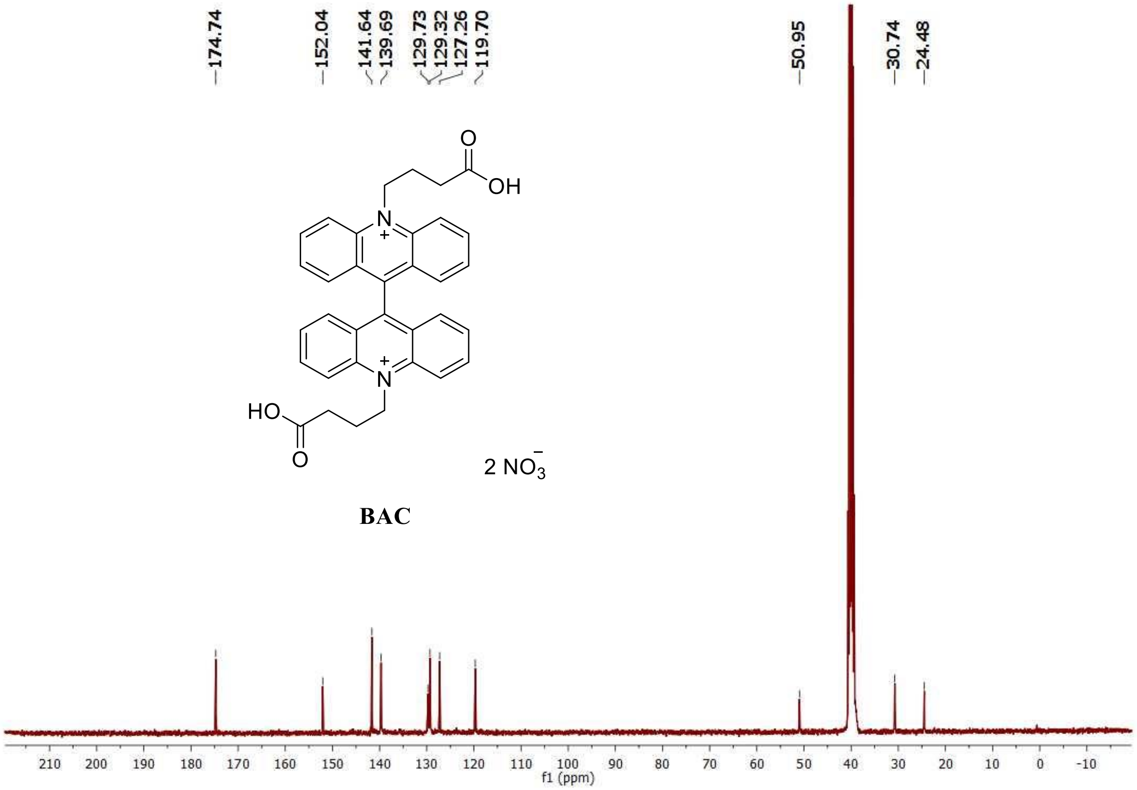


**Figure S3.** $^{13}C$ NMR of BAC.

### Synthesis of compound 1

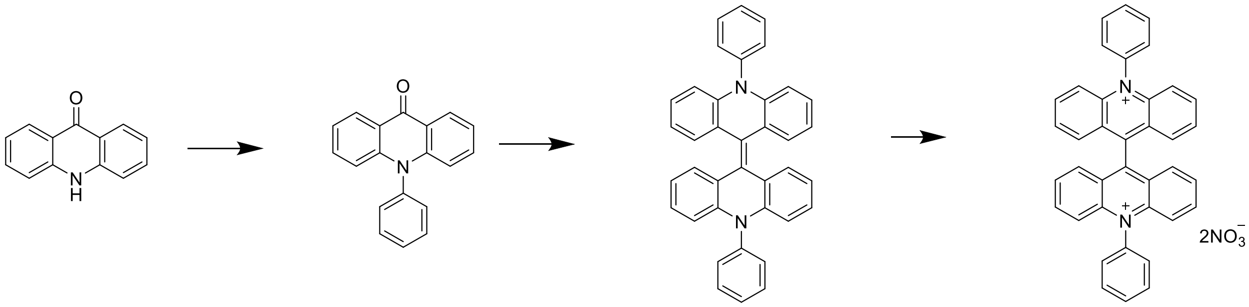


a1 a2 a3 1

**Scheme 2.** Synthesis of compound **1**.

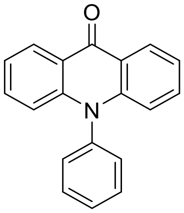


**a2**

10-phenylacridin-9(10H)-one (**a2**)

Compound **a2** was synthesized following general procedure GP3. $^1$H NMR (400 MHz, $CDCl_3$) δ 8.63 (d, *J* = 8.0 Hz, 2H), 7.74 (t, *J* = 7.4 Hz, 2H), 7.68 (d, *J* = 7.1 Hz, 1H), 7.52 (t, *J* = 7.9 Hz, 2H), 7.40 (d, *J* = 7.1 Hz, 2H), 7.34 – 7.28 (m, 2H), 6.78 (d, *J* = 8.6 Hz, 2H). HRMS $[M+H]^+$ (+ TOF MS) calculated for $C_{19}H_{14}NO$ 272.0997 found: 272.1053

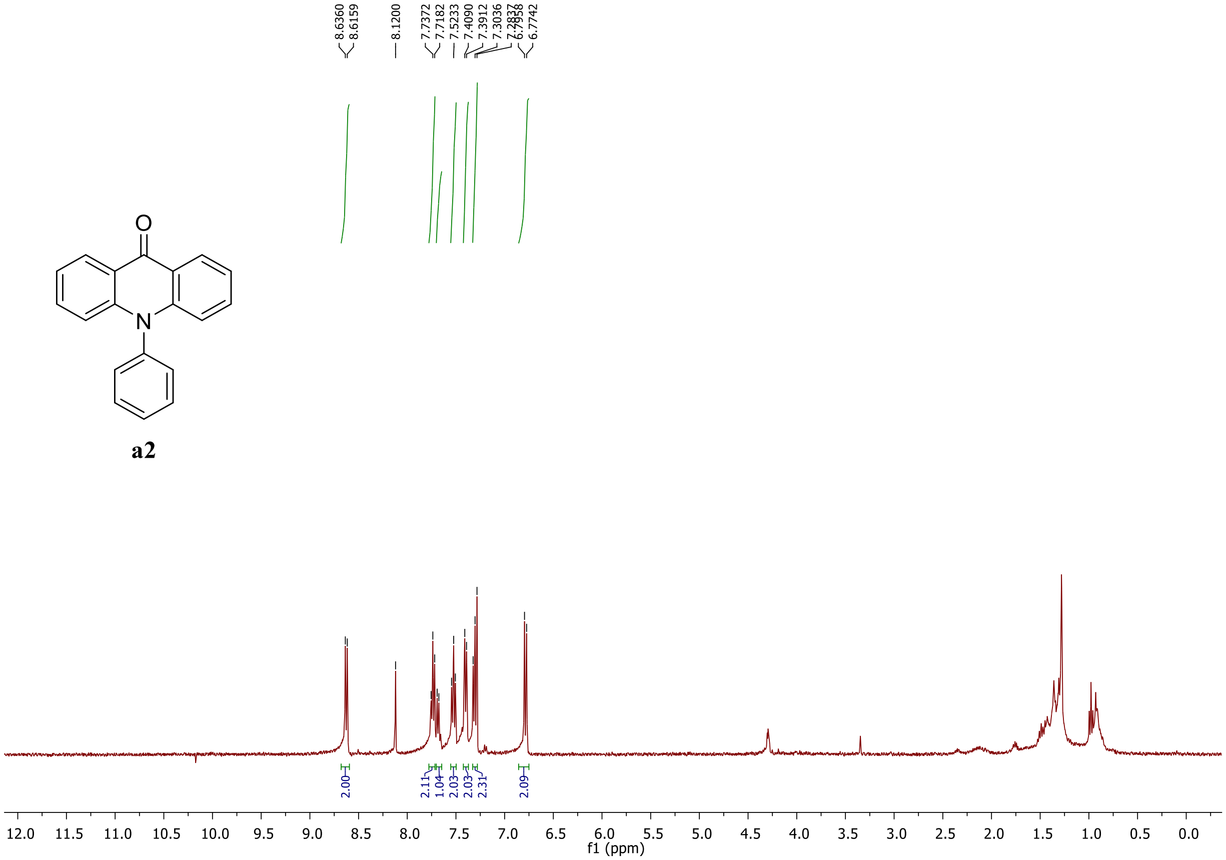


**Figure S4.** $^1$H NMR of compound **a2**.

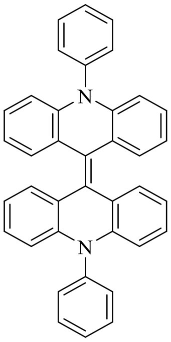

**a3**

10,10'-diphenyl-10H,10'H-9,9'-biacridinylidene (**a3**)

Compounds **a3** was synthesized following general procedure GP4. HRMS [M+H] (+ TOF MS) calculated for $C_{38}H_{26}N_2$ 511.2096 found: 511.2147

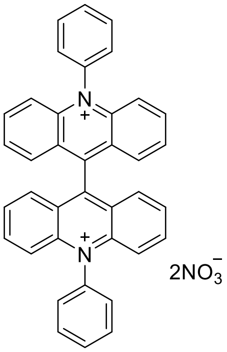

**1**

10,10'-diphenyl-[9,9'-biacridine]-10,10'-diium (**1**)

Compounds **1** was synthesized following general procedure GP4. $^1$H NMR (400 MHz,DMSO-$d_6$) δ 8.33 (dd, *J* = 8.4, 6.9 Hz, 4H), 8.05 (dt, *J* = 14.3, 7.6 Hz, 6H), 7.94 (d, *J* = 7.4 Hz, 4H), 7.91 – 7.81 (m, 8H), 7.73 (d, *J* = 9.1 Hz, 4H). $^{13}$C NMR (100 MHz, ) δ 162.22, 152.91, 142.70, 140.06, 137.90, 132.10, 129.86, 129.15, 128.26, 127.12, 120.83. HRMS [M] (+ TOF MS) calculated for $C_{38}H_{26}N_2$ 510.2085 found: 527.2094.

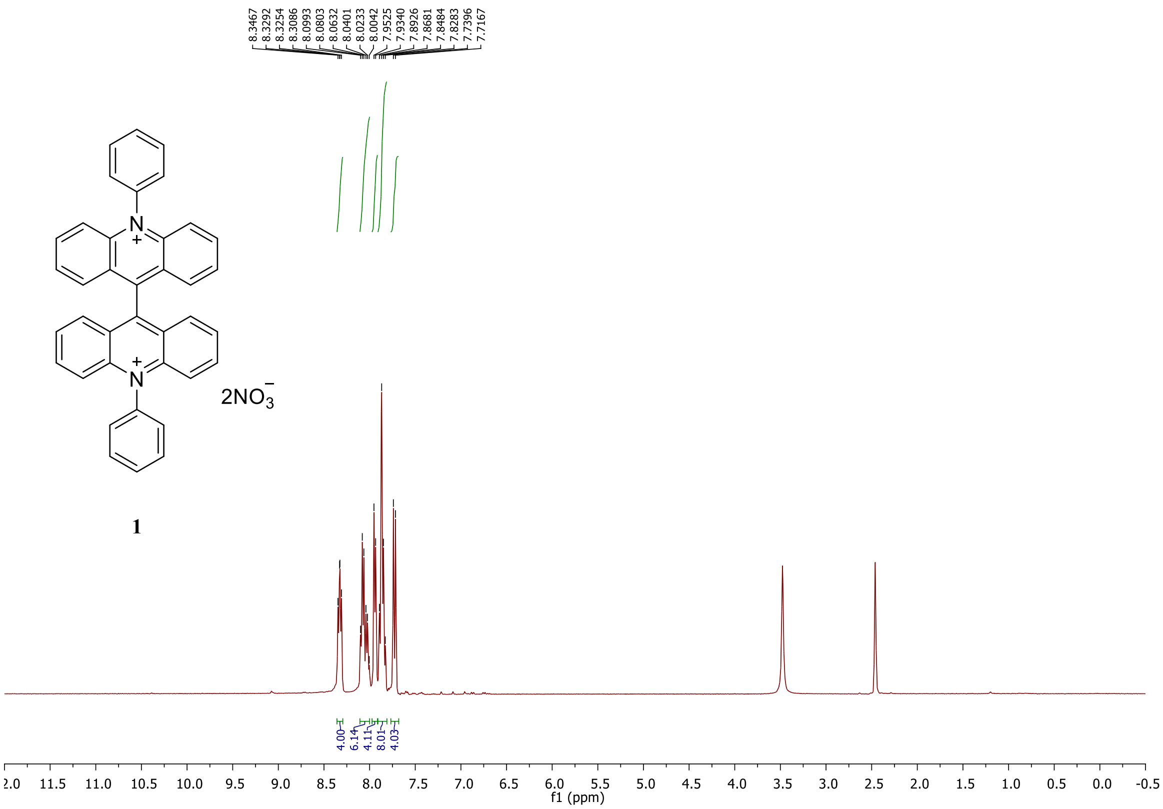


**Figure S5.** $^1$H NMR of compound **1**.

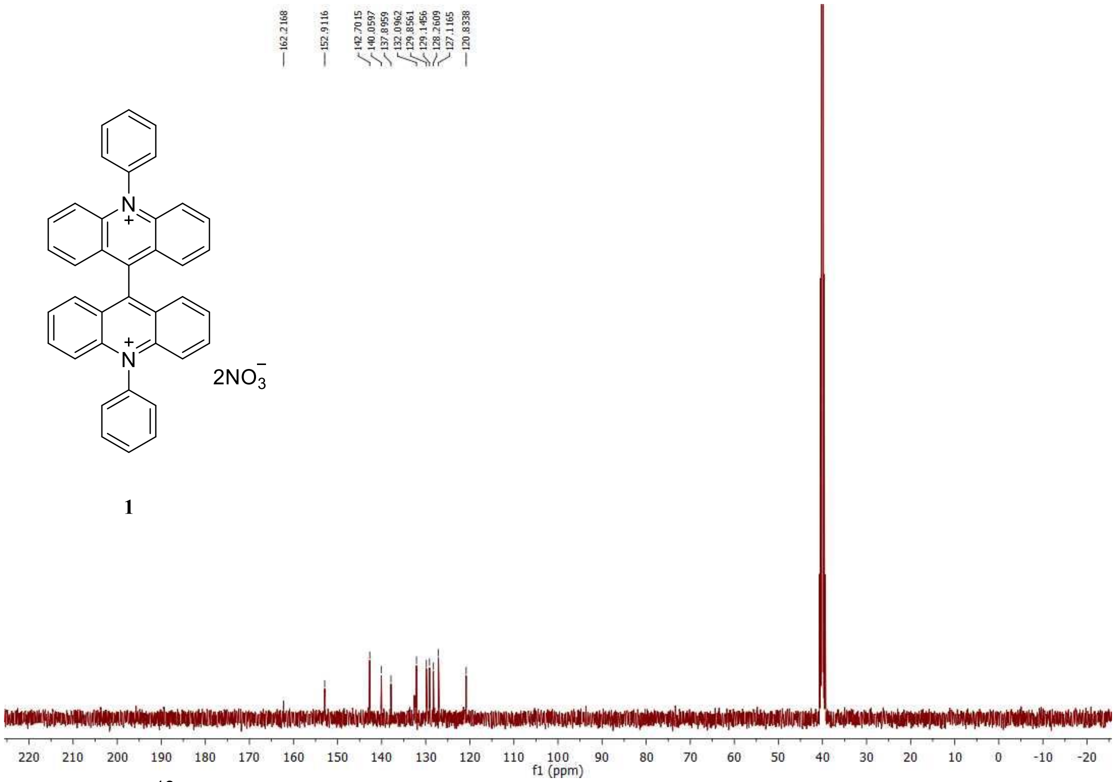

**Figure S6.** $^{13}C$ NMR of compound **1**

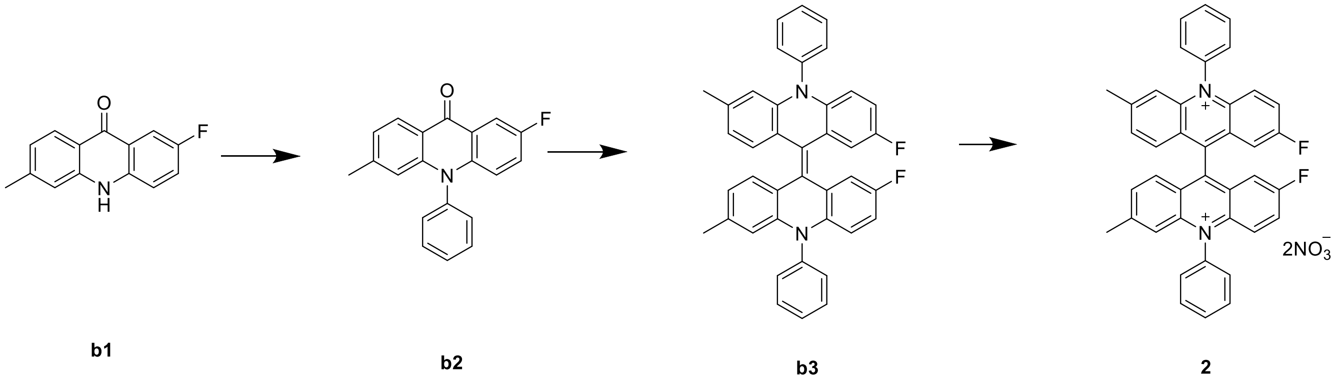

**Scheme 4.** Synthesis of compound **2**

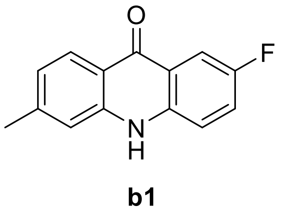

2-fluoro-6-methylacridin-9(10H)-one (**b1**)

Compound **b1** was synthesized following general procedures GP1 and GP2

$^1$H NMR (400 MHz, ) δ 7.93 (d, *J* = 8.0 Hz, 2H), 7.27 (d, *J* = 19.3 Hz, 2H), 7.14 (d, *J* = 7.9 Hz, 2H), 2.38 (s, 3H). HRMS $[M+H]^+$ (+ TOF MS) calculated for $C_{14}H_{10}NO$ 228.0746 found: 228.0820

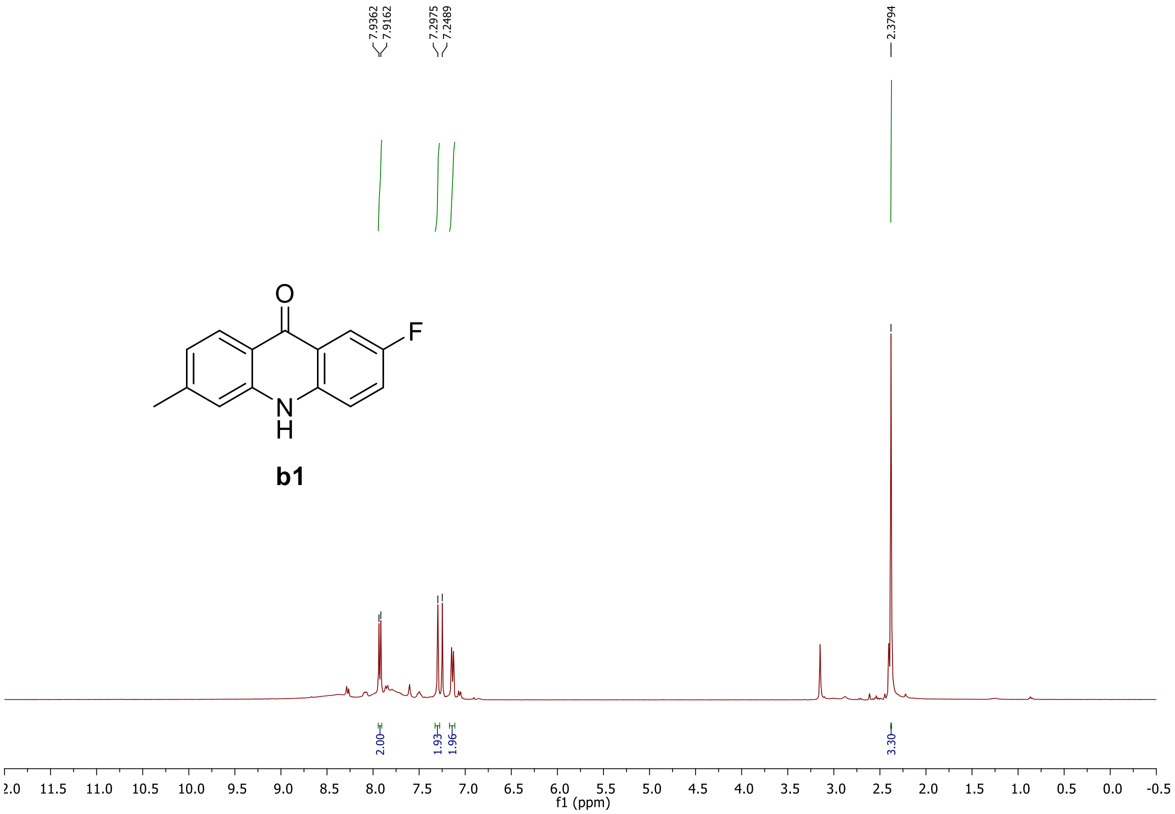


**Figure S7.** $^{1}H$ NMR of compound **b1**

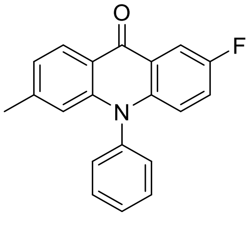


**b2**

2-fluoro-6-methyl-10-phenylacridin-9(10H)-one (**b2**)
Compound **b2** was synthesized following general procedure GP3. $^{1}H$ NMR (400 MHz, $CDCl_3$) δ 8.47 (d, *J* = 7.7 Hz, 1H), 8.21 (d, *J* = 6.1 Hz, 1H), 7.79 – 7.61 (m, 3H), 7.35 (d, *J* = 6.5 Hz, 2H), 7.22 (s, 1H), 7.11 (d, *J* = 7.2 Hz, 1H), 6.72 (d, *J* = 5.0 Hz, 1H), 6.51 (s, 1H), 2.32 (s, 1H). HRMS [M+H] (+ TOF MS) calculated for $C_{20}H_{14}FNO$ 304.1059 found: 304.1119

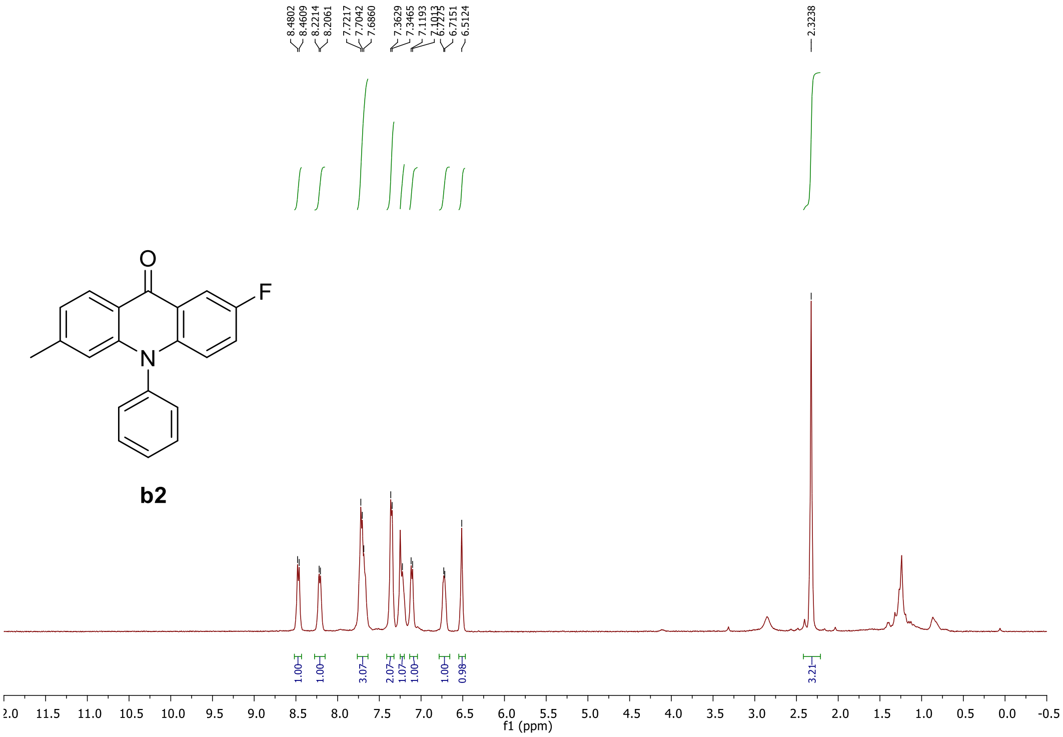


**Figure S8.** $^1$H NMR of compound **b2**

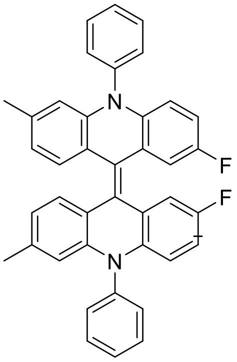

**b3**

(Z)-2,2'-difluoro-6,6'-dimethyl-10,10'-diphenyl-10H,10'H-9,9'-biacridinylidene (**b3**) Compounds **b3** and **2** was synthesized following general procedure GP4. HRMS m/z (+ TOF MS) calculated for $C_{42}H_{32}N_2O_4$ 573.2137 found: 573.2132

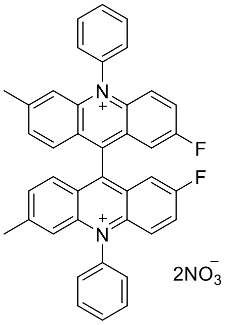

**2**

2,2'-difluoro-6,6'-dimethyl-10,10'-diphenyl-[9,9'-biacridine]-10,10'-diium (**2**) $^1$H NMR (400 MHz, $CDCl_3$ ) δ 8.24 (t, J = 7.7 Hz, 2H), 8.13 – 7.98 (m, 6H), 7.87 (dd, J = 19.5, 7.0 Hz, 4H), 7.74 (t, 6H), 7.70 (dd, 2H), 7.46 (s, 2H), 2.60 (s, 6H). $^{13}$C NMR (101 MHz, $CDCl_3$) δ 152.84, 142.64, 140.07, 137.83, 132.51, 132.02, 129.80, 129.09, 128.21, 127.05, 120.78. HRMS m/z (+ TOF MS) calculated for $C_{42}H_{32}N_2O_4$ 333.1160 found: 333.1173

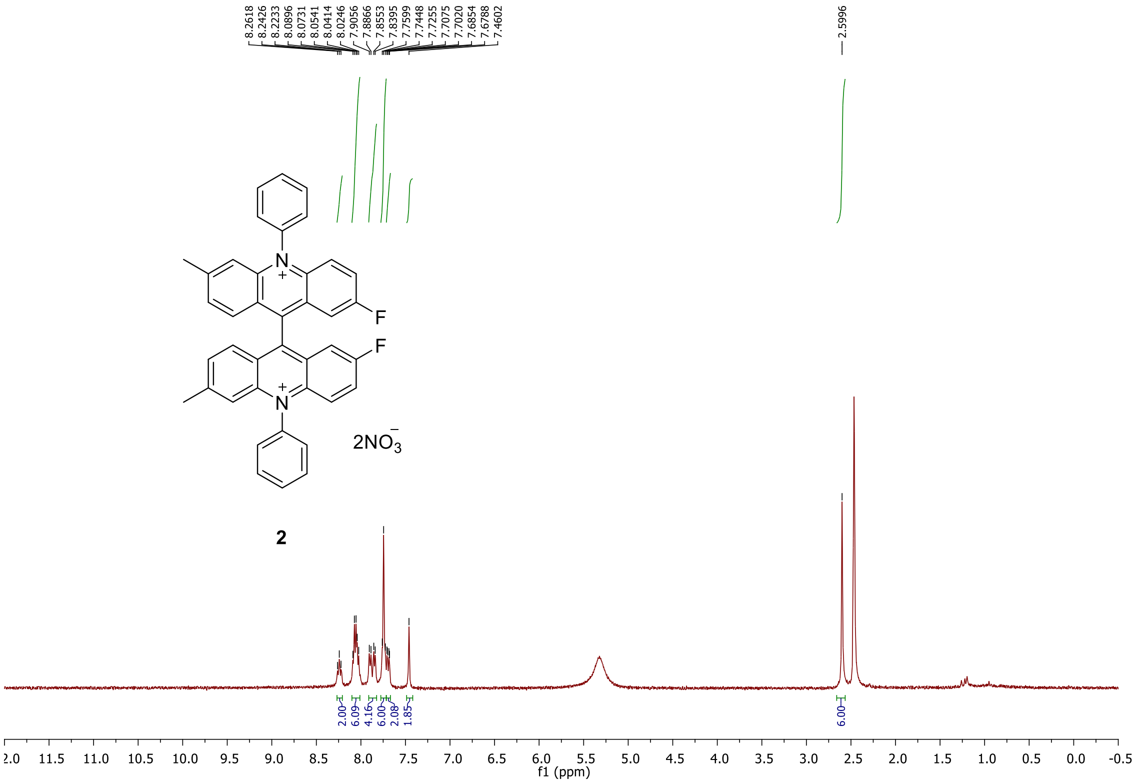


**Figure S9.** $^{1}$H NMR of compound **2**.

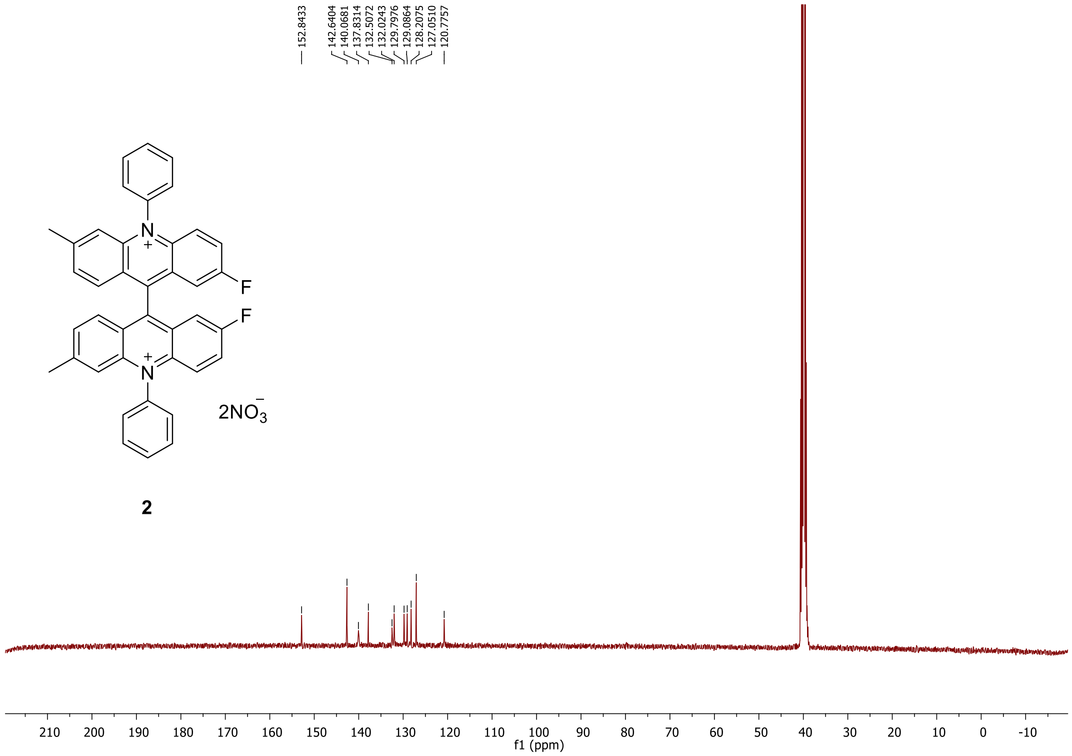


**Figure S10.** $^{13}$C NMR of compound **2**.

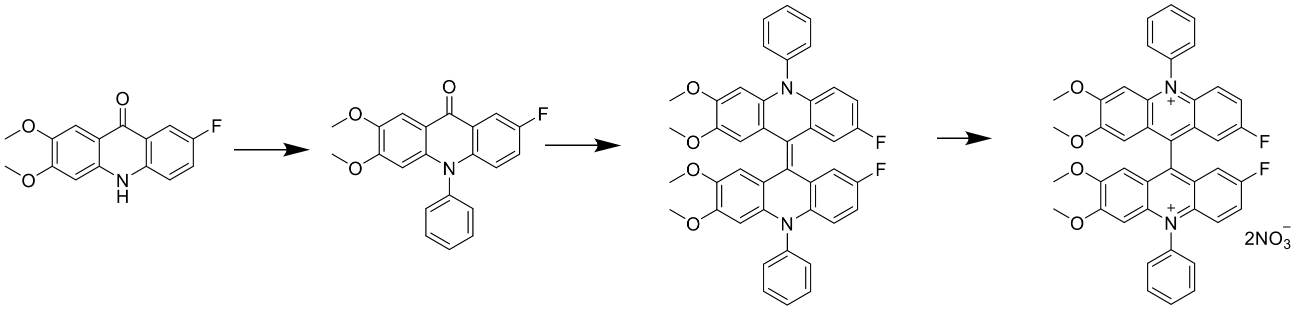

**c1** **c2** **c3** **3**

**Scheme 5.** Synthesis of compound **3**

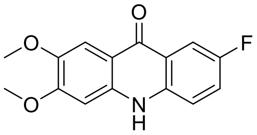

**c1**

7-fluoro-2,3-dimethoxyacridin-9(10H)-one (**c1**)
Compound **c1** was synthesized following general procedures GP1 and GP2. $^{1}H$ NMR (400 MHz, DMSO-$d_6$) δ 11.74 (s, 1H), 7.82 – 7.71 (m, 1H), 7.59 – 7.44 (m, 3H), 6.89 (s, 1H), 3.88 (s, 3H), 3.81 (s, 3H). HRMS [M+H] (+ TOF MS) calculated for $C_{15}H_{12}FNO_3$ 274.0801 found: 274.0861

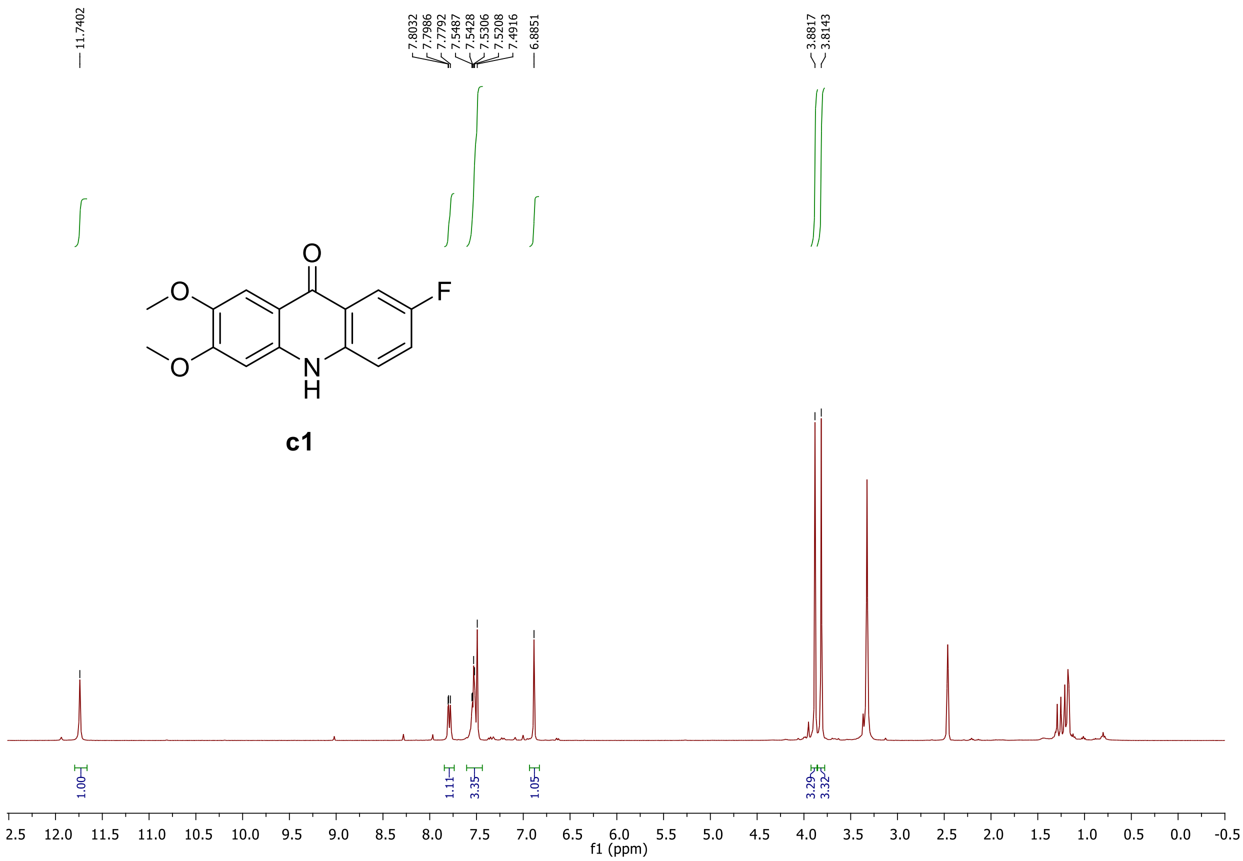


**Figure S11.** $^{1}H$ NMR of compound **c1**

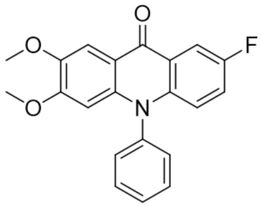

**c2**

7-fluoro-2,3-dimethoxy-10-phenylacridin-9(10H)-one (**c2**)
Compound **c2** was synthesized following general procedure GP3. $^1$H NMR NMR (400 MHz, $CDCl_3$) δ 8.20 (dd, *J* = 9.0, 3.0 Hz, 1H), 7.97 (s), 7.70 (dq, *J* = 14.6, 7.3 Hz, 1H), 7.38 (d, *J* = 7.2 Hz, 3H), 7.21 (ddd, *J* = 10.3, 7.6, 3.1 Hz, 2H), 6.75 (dd, *J* = 9.4, 4.3 Hz, 1H), 6.09 (s, 1H), 4.01 (s, 3H), 3.64 (s, 3H). HRMS [M+H] (+ TOF MS) calculated for $C_{21}H_{16}FNO_3$ 350.1114 found: 350.1176

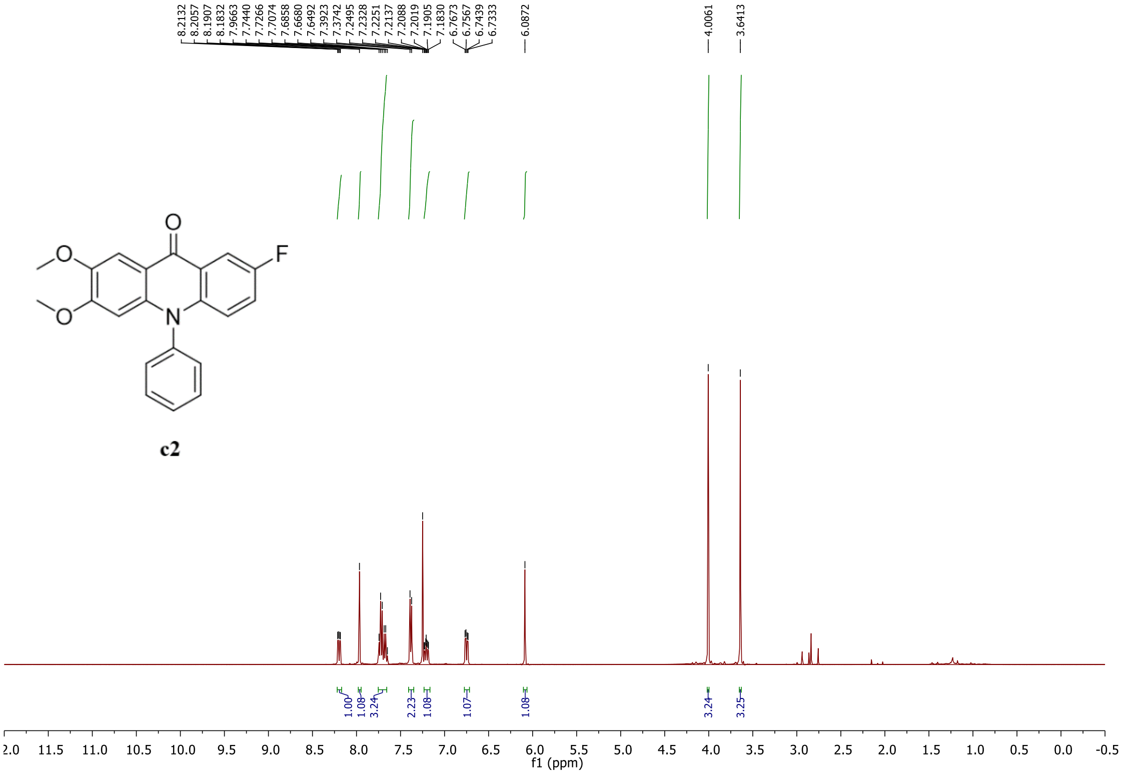


**Figure S11.** $^1$H NMR of compound **c2**

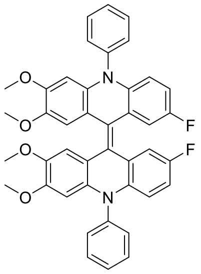

**c3**

(Z)-7,7'-difluoro-2,2',3,3'-tetramethoxy-10,10'-diphenyl-10H,10'H-9,9'-biacridinylidene (**c3**)
Compounds **c3** was synthesized following general procedure GP4. HRMS [M+H] (+ TOF MS) calculated for $C_{42}H_{34}N_2O_4$ 630.2519 found: 630.2611

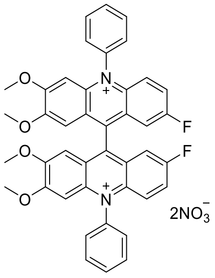

**3**

7,7'-difluoro-2,2',3,3'-tetramethoxy-10,10'-diphenyl-[9,9'-biacridine]-10,10'-diium (**3**) Compounds **c3** was synthesized following general procedure GP4. $^{1}H$ NMR (400 MHz, DMSO-$d_6$) δ 8.12 – 7.97 (m 8H), 7.90 (dd, *J* = 14.7, 7.8 Hz, 4H), 7.63 (dd, *J* = 9.8, 4.6 Hz, 2H), 7.59 – 7.51 (m, 2H), 6.82 (s, 2H), 6.67 (s, 2H), 3.85 (s, 6H), 3.59 (s, 6H). $^{13}C$ NMR (101 MHz, DMSO-$d_6$ ) δ 161.63, 153.19, 143.45, 138.54, 138.31, 132.62, 128.68, 125.67, 105.11, 99.40, 58.25. HRMS [M] (+ TOF MS) calculated for $C_{42}H_{32}F_2N_2O_4$ 666.2319 found: 666.2328

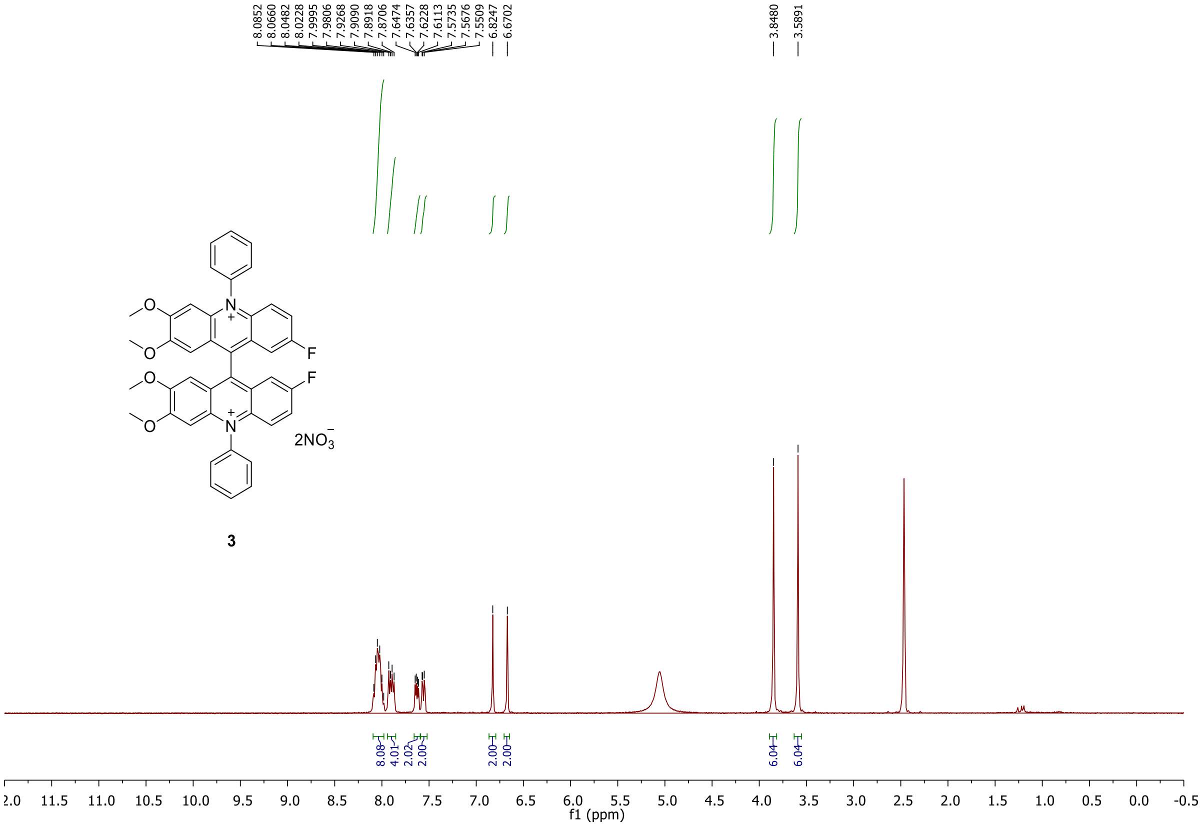


**Figure S12.** $^{1}H$ NMR of compound **3**

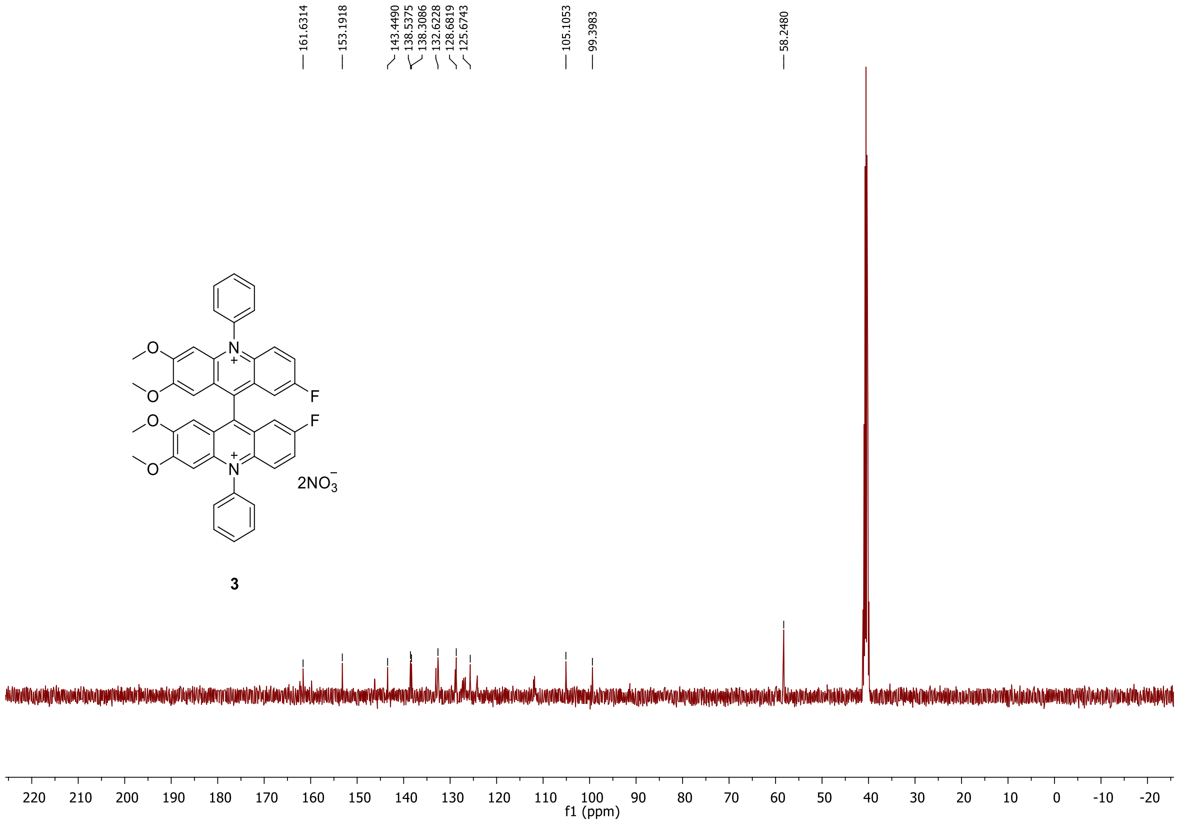


**Figure S13.** $^{13}$C NMR of compound **3**

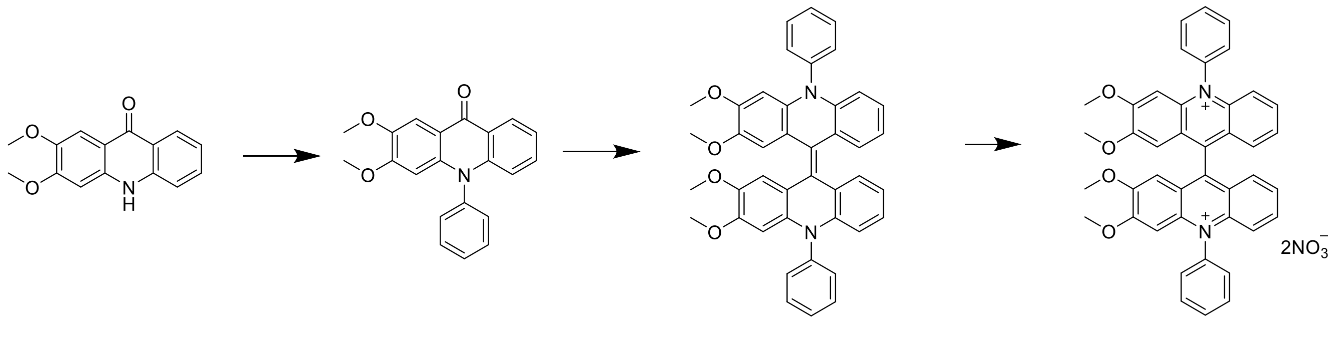


**d1** **d2** **d3** **4**

**Scheme 6.** Synthesis of compound **4**

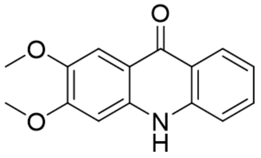

**d1**

2,3-dimethoxyacridin-9(10H)-one (**d1**)
Compound **d1** was synthesized following general procedures GP1 and GP2
$^{1}$H NMR NMR (400 MHz, DMSO-d$_6$ ) δ 11.63 (s, 1H), 8.17 (d, *J* = 7.7 Hz, 1H), 7.62 (t, *J* = 7.3 Hz, 1H), 7.53 (s, 1H), 7.46 (d, *J* = 8.2 Hz, 1H), 7.19 (d, *J* = 7.1 Hz, 1H), 6.92 (s, 1H), 3.88 (s, 3H), 3.82 (s, 3H).HRMS [M+H] (+ TOF MS) calculated for $C_{15}H_{13}NO_3$ 256.0895 found: 256.0959

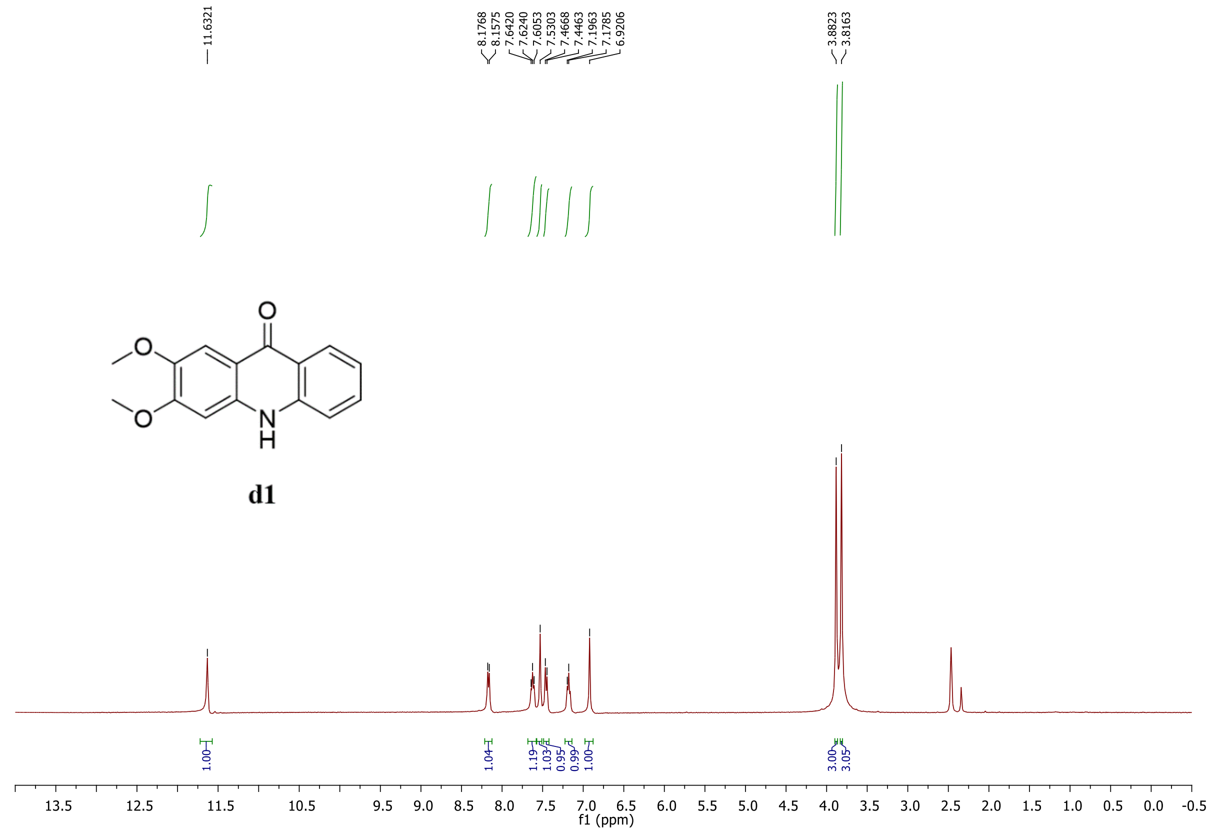


**Figure S14.** $^{1}$H NMR of compound **d1**

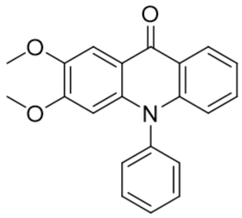

**d2**

Synthesis of 2,3-dimethoxyacridin-9(10H)-one (**d2**)
Compound **d2** was synthesized following general procedure GP3.
$^{1}$H NMR (400 MHz, $CDCl_3$ ) δ 8.65 (d, J = 8.1 Hz, 1H), 8.10 (s, 1H), 7.76 – 7.64 (m, 3H), 7.50 (dd, J = 8.4, 7.2 Hz, 1H), 7.38 (d, J = 7.8 Hz, 2H), 7.32 (t, J = 7.6 Hz, 1H), 6.78 (d, J = 8.7 Hz, 1H), 6.12 (s, 1H), 4.04 (s, 3H), 3.65 (s, 3H).
HRMS [M+H] (+ TOF MS) calculated for $C_{21}H_{18}NO_3$ 332.1208 found: 332.1248

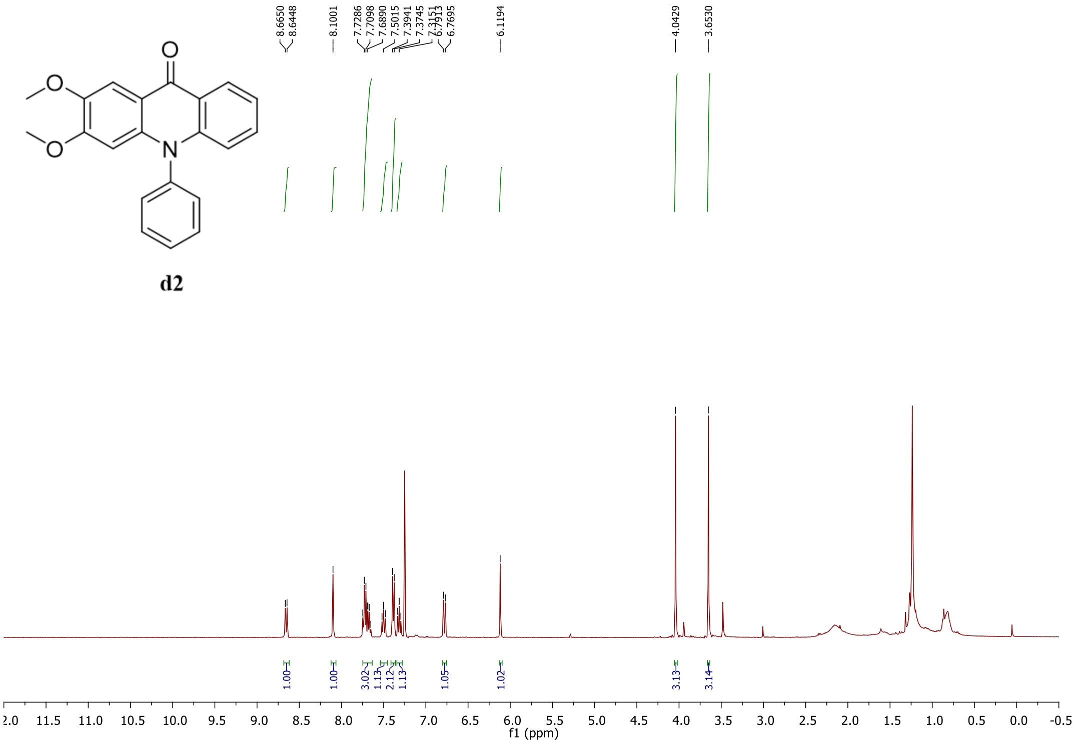


**Figure S15.** $^{1}$H NMR of compound **d2**

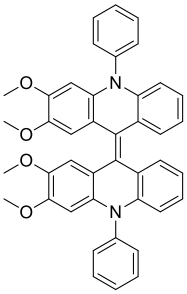

**d3**

Synthesis of (Z)-2,2',3,3'-tetramethoxy-10,10'-diphenyl-10H,10'H-9,9'-biacridinylidene (**d3**)
Compounds **d3** and **4** were synthesized following general procedure GP4. HRMS $[M+H]^+$ (+ TOF MS) calculated for $C_{42}H_{35}N_2O_4$ 631.2519 found: 631.2611

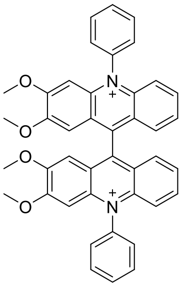

**4**

2,2',3,3'-tetramethoxy-10,10'-diphenyl-[9,9'-biacridine]-10,10'-diium (**4**)
$^{1}$H NMR (400 MHz, DMSO-$d_6$) δ 8.14 – 8.09 (m, 2H), 8.08 – 7.99 (m, 6H), 7.96 (d, *J* = 7.8 Hz, 4H), 7.71 (t, *J* = 6.9 Hz, 4H), 7.54 (d, *J* = 9.1 Hz, 2H), 6.90 (s, 2H), 6.70 (s, 2H), 3.85 (s, 3H), 3.57 (s, 3H). $^{13}$C NMR (101 MHz, DMSO-$d_6$) δ 160.93, 152.07, 147.22, 142.86, 140.25, 137.96, 136.67, 132.35, 132.04, 128.57, 128.36, 128.22, 125.29, 124.25, 120.14, 57.68. HRMS m/z (+ TOF MS) calculated for $C_{21}H_{17}NO_2$ 315.1254 found: 315.1270.

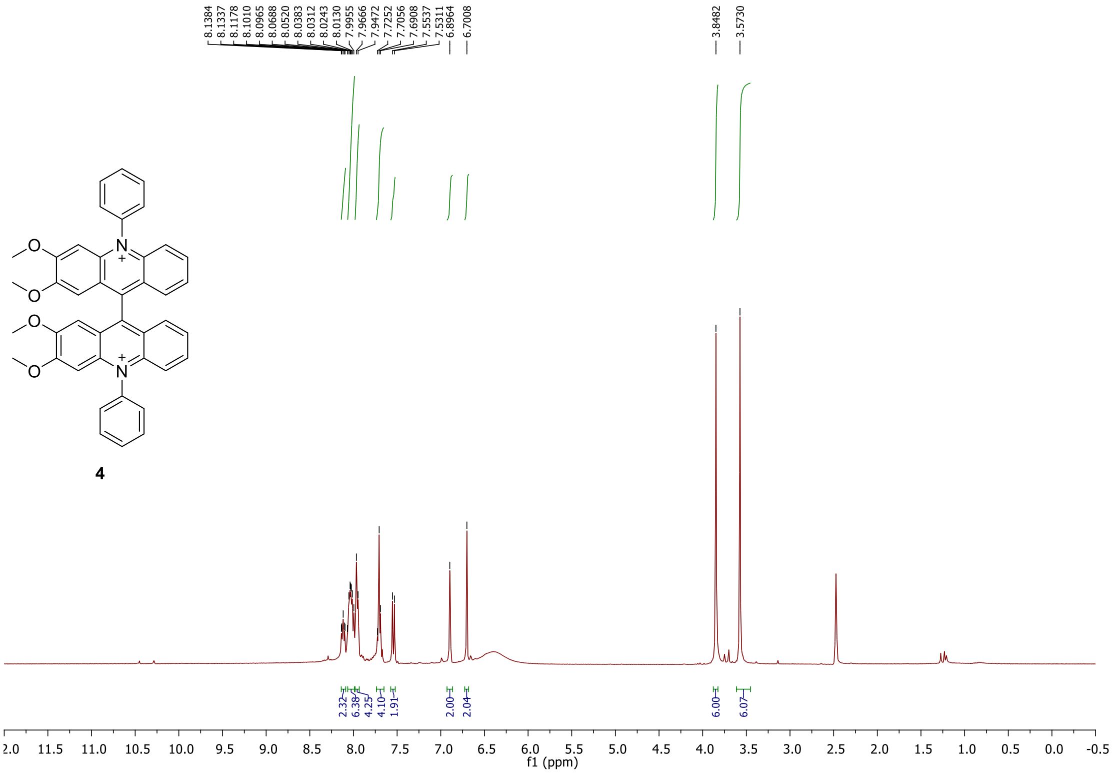

**Figure S16.** $^{1}H$ NMR of compound **4**

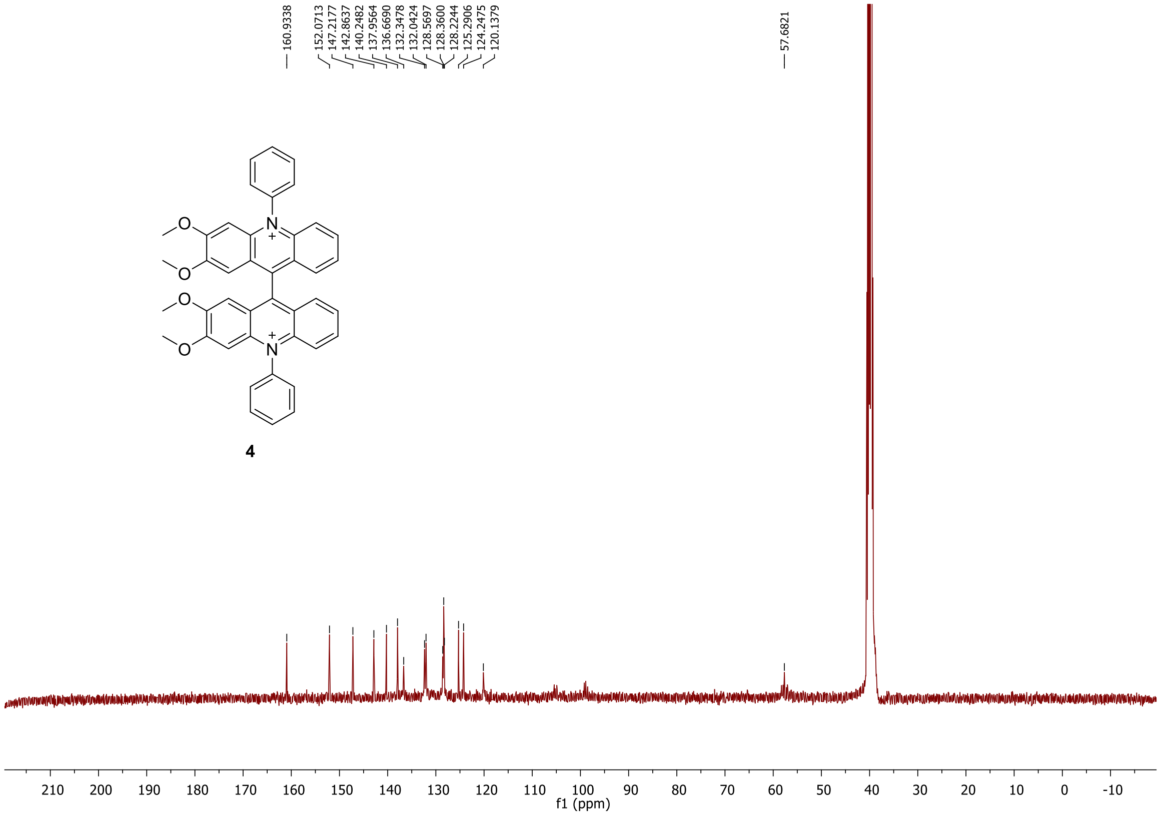

**Figure S17.** $^{13}C$ NMR of compound **4**

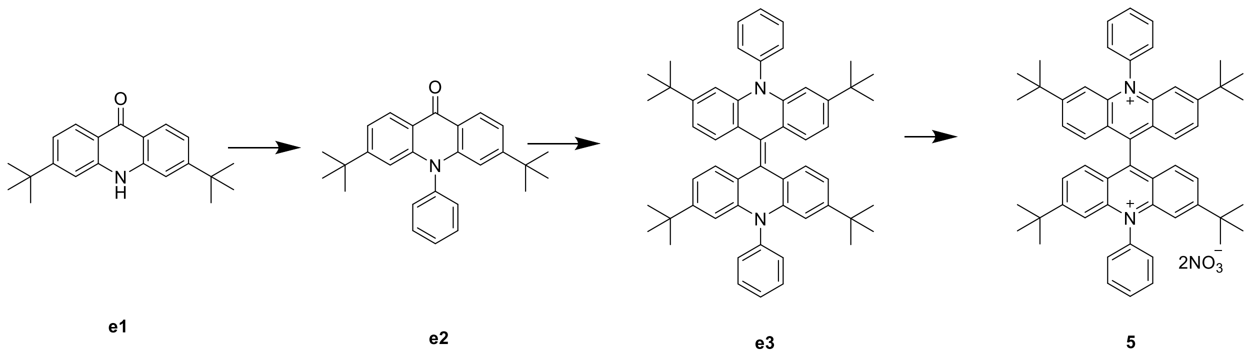


**Scheme 7.** Synthesis of compound **5**

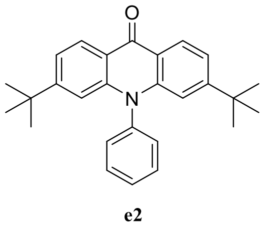


Synthesis of 3,6-di-tert-butyl-10-phenylacridin-9(10H)-one (**e2**)

Compound **e2** was synthesized following general procedure GP3. $^1$H NMR (400 MHz, $CDCl_3$) δ 8.53 (d, *J* = 8.5 Hz, 2H), 7.69 (tt, *J* = 14.7, 7.3 Hz, 3H), 7.35 (t, *J* = 9.2 Hz, 4H), 6.71 (s, 2H), 1.18 (d, *J* = 0.5 Hz, 18H). HRMS [M+H] (+ TOF MS) calculated for $C_{27}H_{30}NO$ 384.2249 found: 384.2305

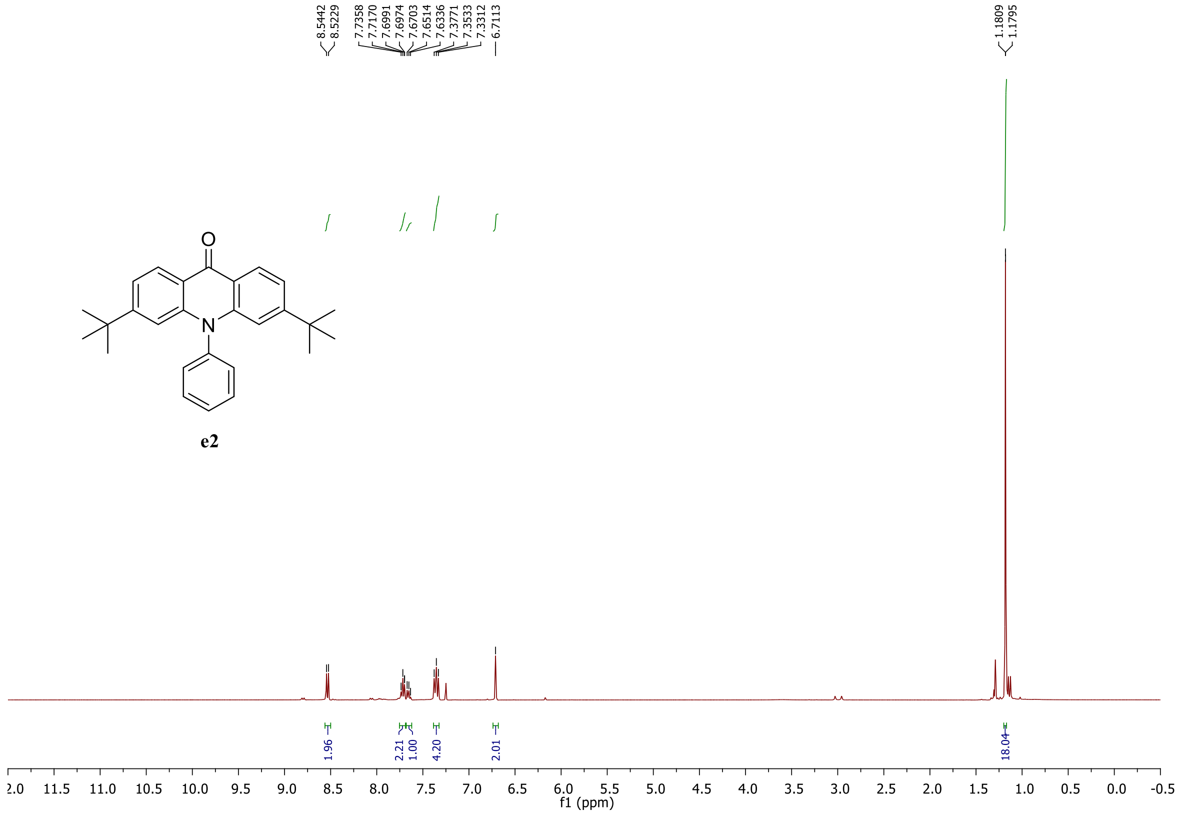


**Figure S18.** $^1$H NMR of compound **e2**

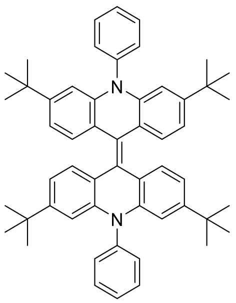

e3

Synthesis of 3,3',6,6'-tetra-tert-butyl-10,10'-diphenyl-10H,10'H-9,9'-biacridinylidene (**e3**) was synthesized following general procedure GP4. HRMS m/z (+ TOF MS) calculated for $C_{27}H_{29}N$ 367.2300 found: 367.2287

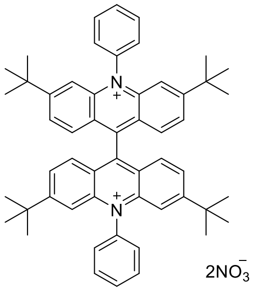

**5**

Synthesis of 3,3',6,6'-tetra-tert-butyl-10,10'-diphenyl-[9,9'-biacridine]-10,10'-diium (**5**)
$^{1}$H NMR (400 MHz, DMSO-d$_6$ ) δ 8.12 – 8.02 (m, 6H), 7.98 (dd, *J* = 9.2, 1.4 Hz, 4H), 7.93 (d, *J* = 7.1 Hz, 4H), 7.78 (d, *J* = 9.1 Hz, 4H), 7.41 (d, *J* = 1.1 Hz, 4H), 1.22 (s, 36 H). $^{13}$C NMR (101 MHz, DMSO-d6) δ 163.25, 151.06, 142.70, 137.58, 132.06, 128.98, 128.15, 125.35, 36.98, 30.05, 29.81.HRMS m/z (+ TOF MS) calculated for $C_{27}H_{29}N$ 367.2295 found: 367.2288

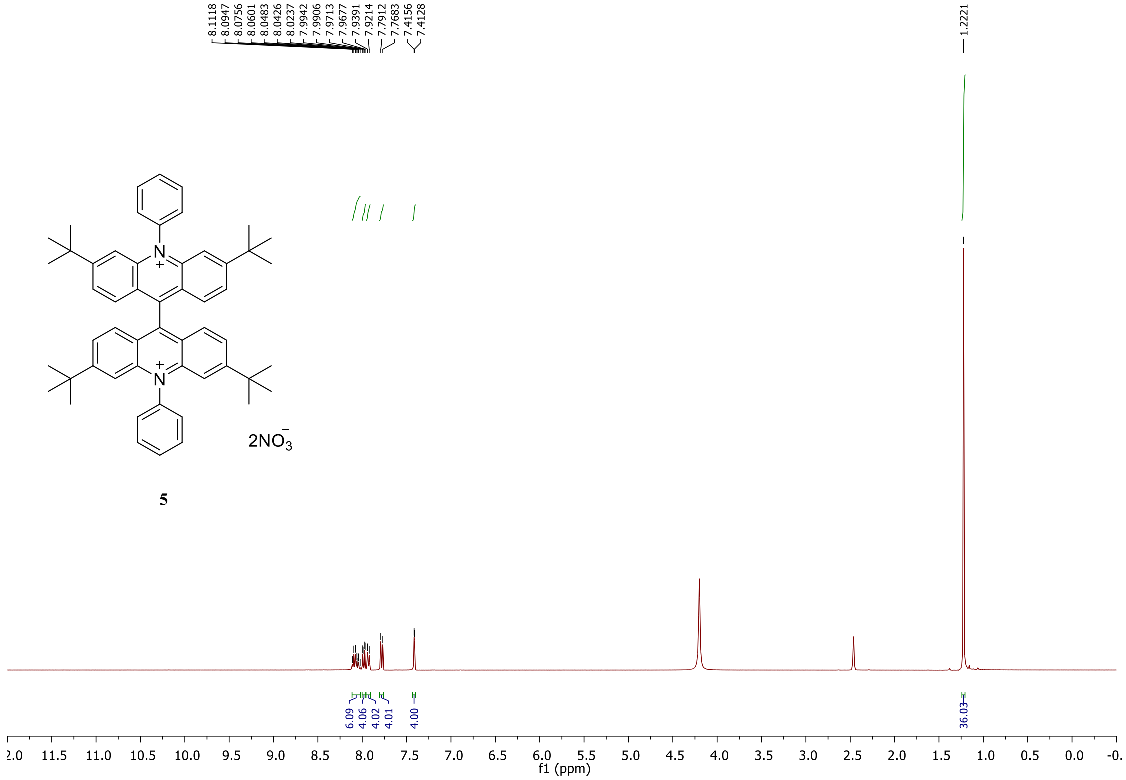


**Figure S19.** $^{1}$H NMR of compound **5**.

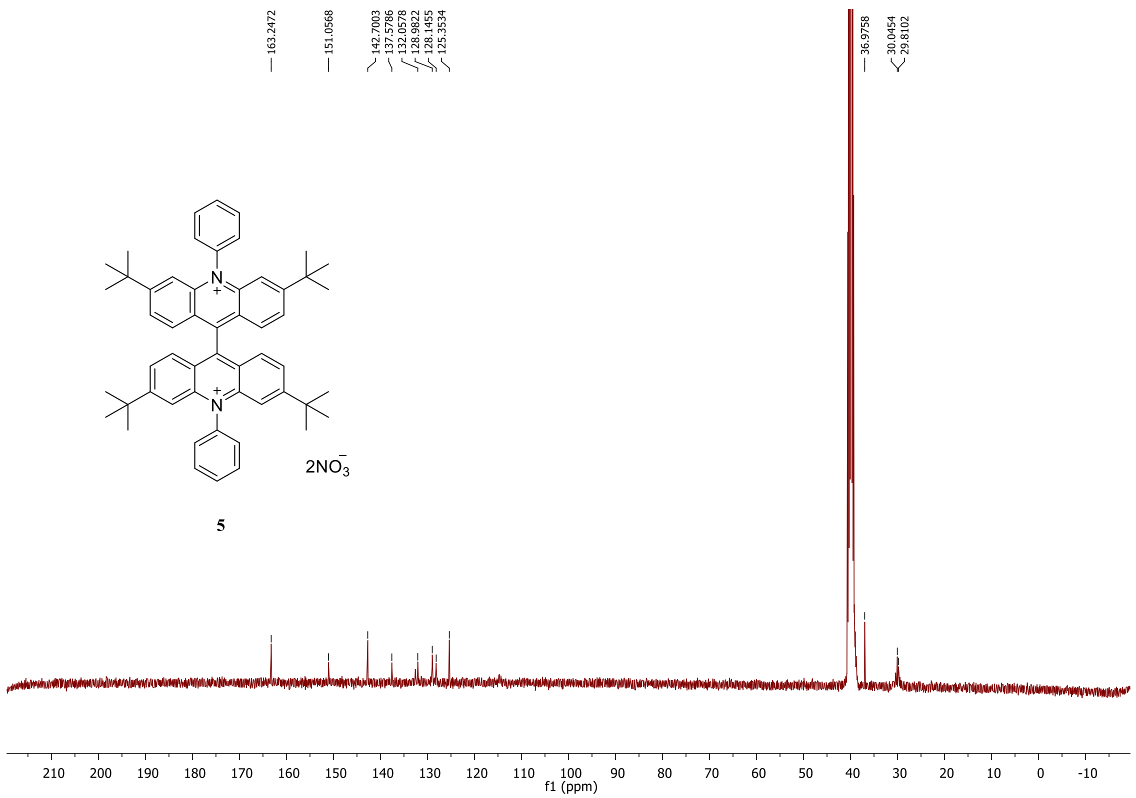


**Figure S20.** $^{13}C$ NMR of compound **5**.

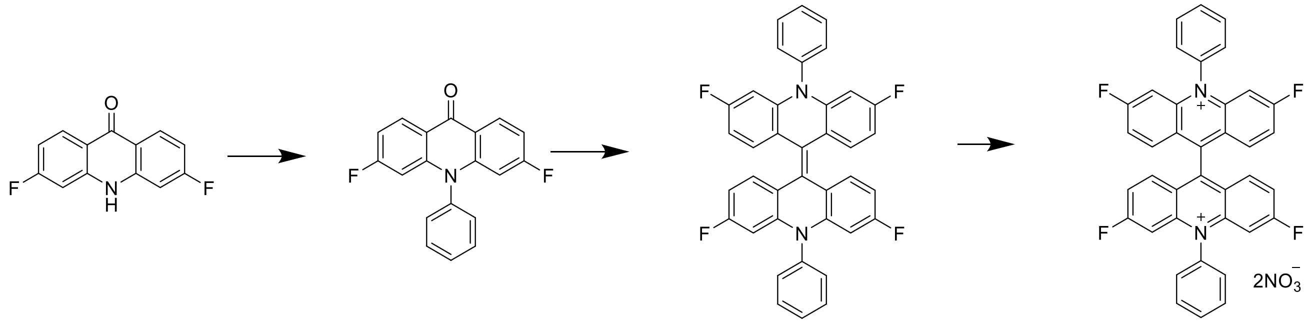


**Scheme 8.** Synthesis of compound **6**

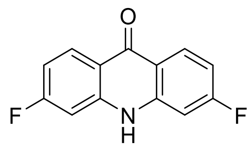

**f1**

Synthesis of 3,6-difluoroacridin-9(10H)-one (**f1**)

Compound **f1** was synthesized following general procedures GP1 and GP2. $^{1}H$ NMR (400 MHz, DMSO-d6 ) δ 11.86 (s, 1H), 8.41 – 8.03 (m, 2H), 7.40 – 6.80 (m, 4H). HRMS $[M+H]^+$ (+ TOF MS) calculated for $C_{13}H_8F_2NO$ 232.0496 found: 232.0558

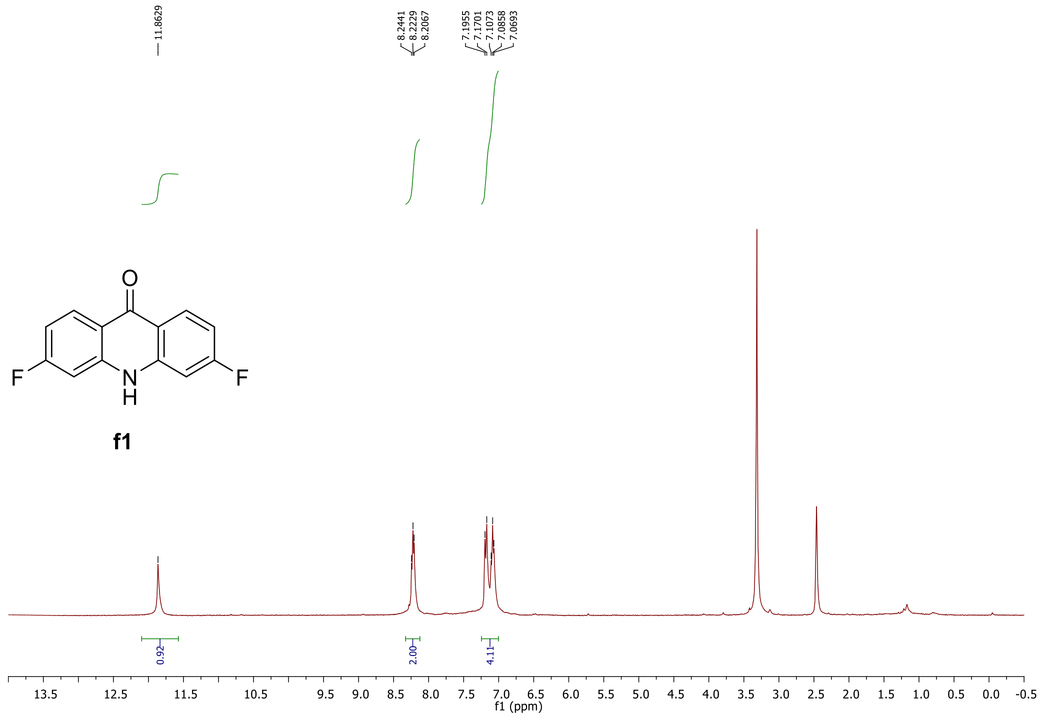


**Figure S21.** $^{1}H$ NMR of compound **f1**.

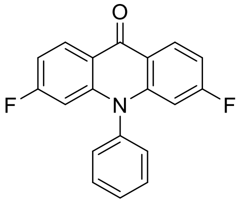

**f2**

Synthesis of 3,6-difluoro-10-phenylacridin-9(10H)-one (**f2**)

Compound **f2** was synthesized following general procedure GP3. $^{1}H$ NMR (400 MHz, $CDCl_3$) δ 8.59 (dd, *J* = 8.8, 6.7 Hz, 2H), 7.72 (dt, *J* = 13.2, 6.8 Hz, 3H), 7.35 (d, *J* = 7.5 Hz, 2H), 7.04 – 6.97 (m, 2H), 6.37 (dd, *J* = 11.2, 2.0 Hz, 2H).

HRMS $[M+H]^+$ (+ TOF MS) calculated for $C_{19}H_{12}F_2NO$ 308.0809 found: 308.0860

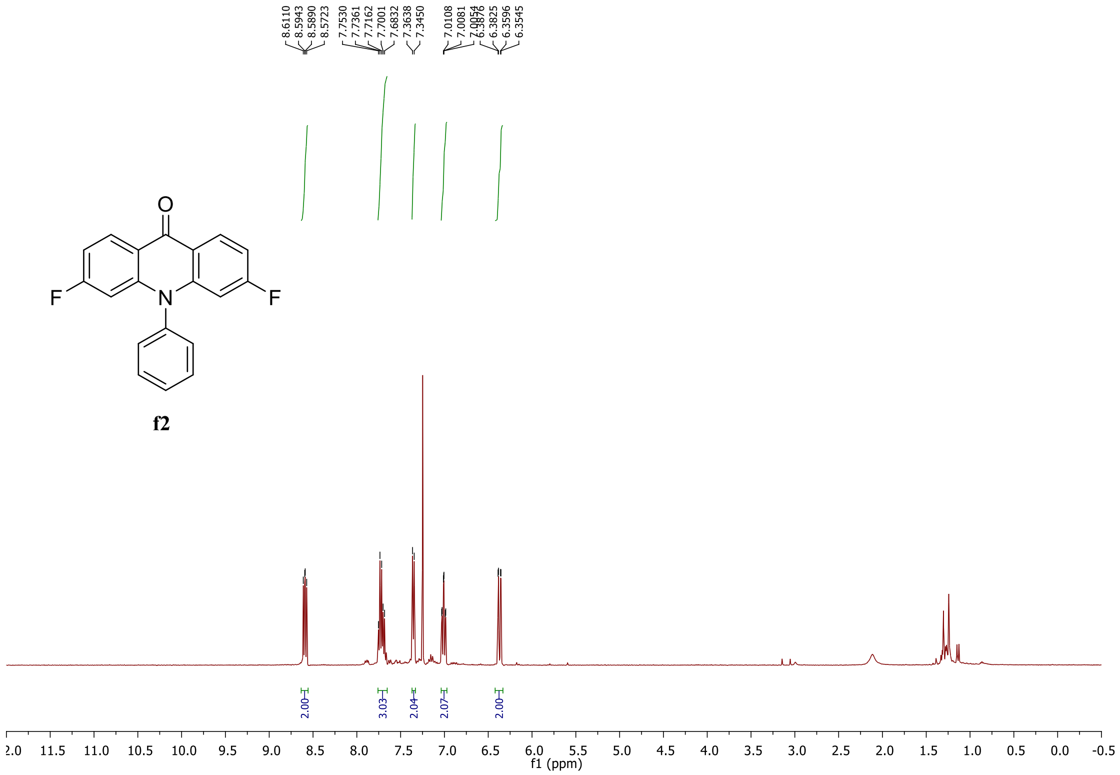


**Figure S22.** $^1$H NMR of compound **f2**.

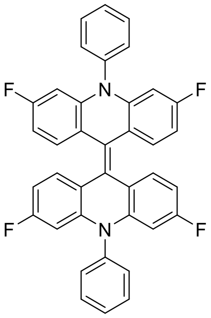

f3

Synthesis of 3,3',6,6'-tetrafluoro-10,10'-diphenyl-10H,10'H-9,9'-biacridinylidene (**f3**)
Compounds **f3** was synthesized following general procedure GP4.
HRMS [M]+ (+ TOF MS) calculated for C38H22F4N 582.1719 found: 582.1698

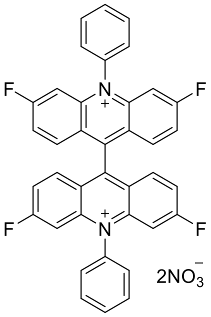

**6**

Synthesis of 3,3',6,6'-tetrafluoro-10,10'-diphenyl-[9,9'-biacridine]-10,10'-diium (**6**)
Compounds **6** was synthesized following general procedure GP4. $^{1}$H NMR (400 MHz, DMSO-d6) δ 8.10 – 8.00 (m, 8H), 7.91 (t, *J* = 8.5 Hz, 4H), 7.84 (dd, *J* = 14.7, 7.6 Hz, 6H), 7.42 (d, *J* = 9.7 Hz, 4H). $^{13}$C NMR (100 MHz, DMSO-d6δ 169.71, 169.21, 168.41, 167.06, 165.81, 152.19, 150.46, 146.88, 145.51, 145.37, 143.79, 137.74, 137.28, 133.59, 132.89, 132.40, 132.09, 128.11, 127.86, 124.45, 122.79, 122.05, 121.65, 121.39, 118.77. HRMS m/z (+ TOF MS) calculated for $C_{19}H_{22}F_2N$ 291.0854 found: 291.0844

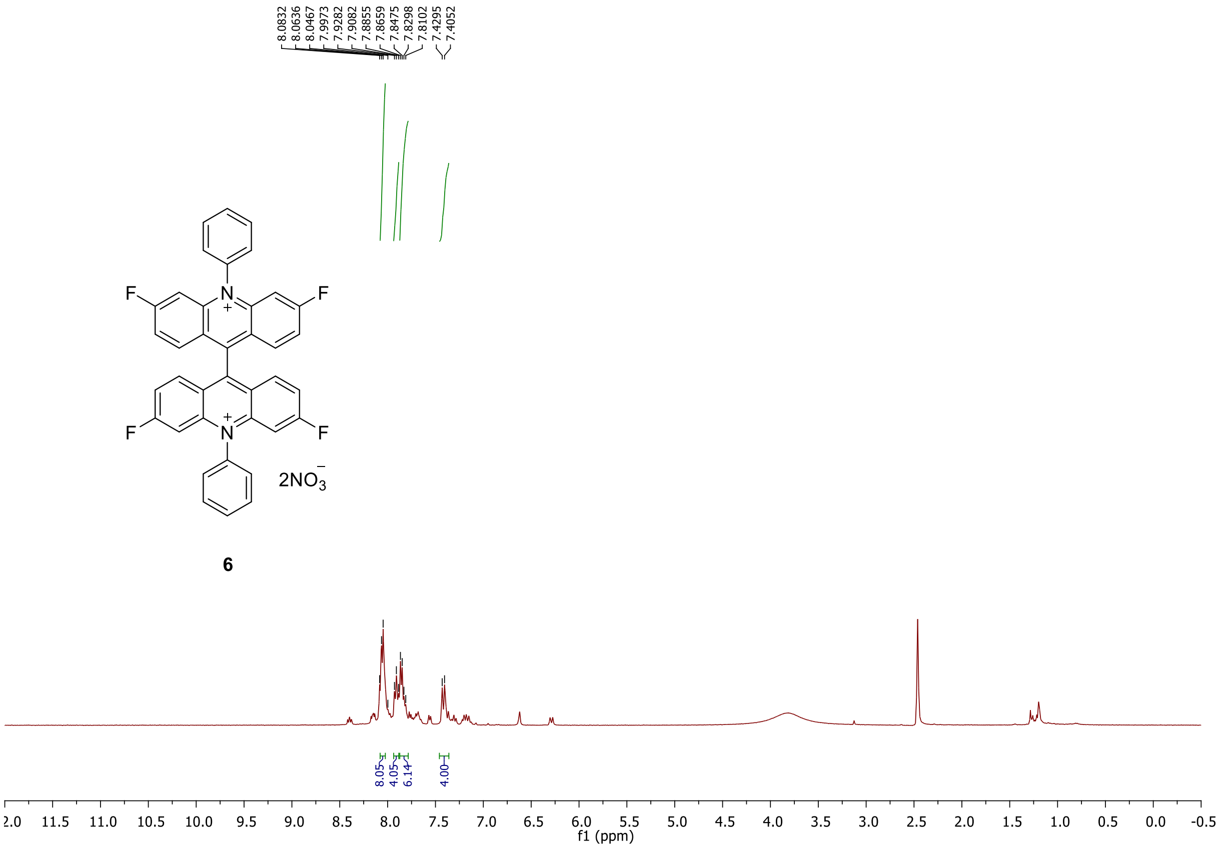


**Figure S23.** $^{1}$H NMR of compound **6**

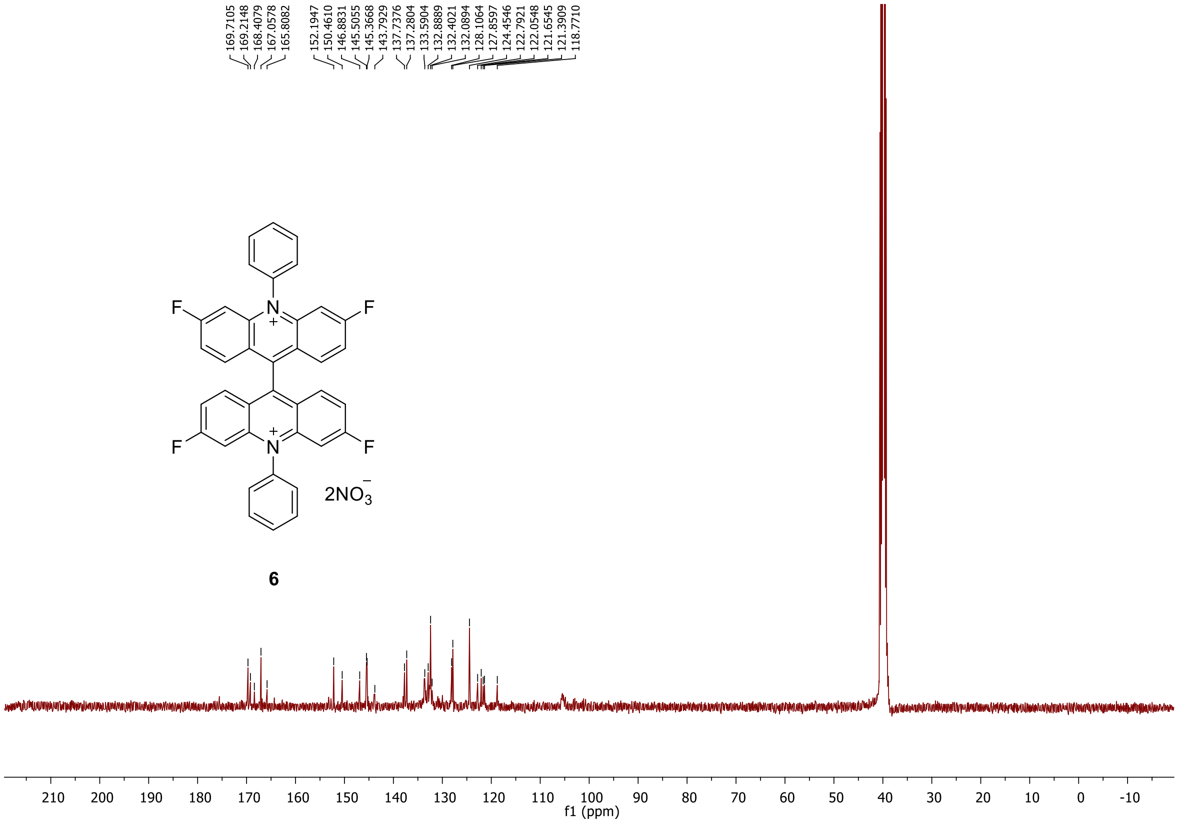


**Figure S24.** $^{13}$C NMR of compound **f1**.

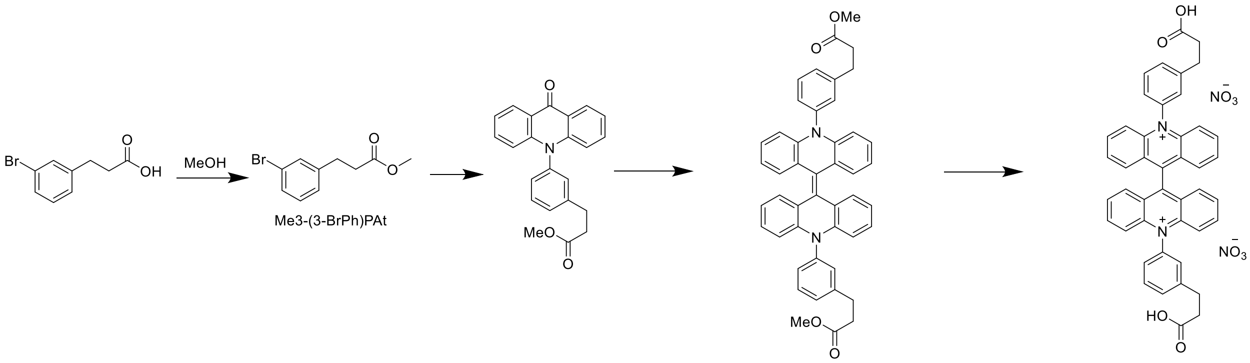


Ib IIb IIIb 1b

**Scheme 1.** Synthesis of compound **1b**.

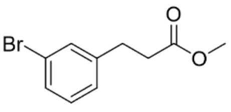


Synthesis of methyl 3-(3-bromophenyl)propanoate (**Ib**)

Methyl 3-(3-bromophenyl)propanoate was synthesized following the reported literature procedure.[6] $^{1}$H NMR (400 MHz, $CDCl_3$) δ 7.33 (dd, $J$ = 5.6, 3.9 Hz, 2H), 7.18 – 7.10 (m, 2H), 3.66 (s, 3H), 2.91 (t, $J$ = 7.7 Hz, 2H), 2.61 (t, $J$ = 7.8 Hz, 2H).

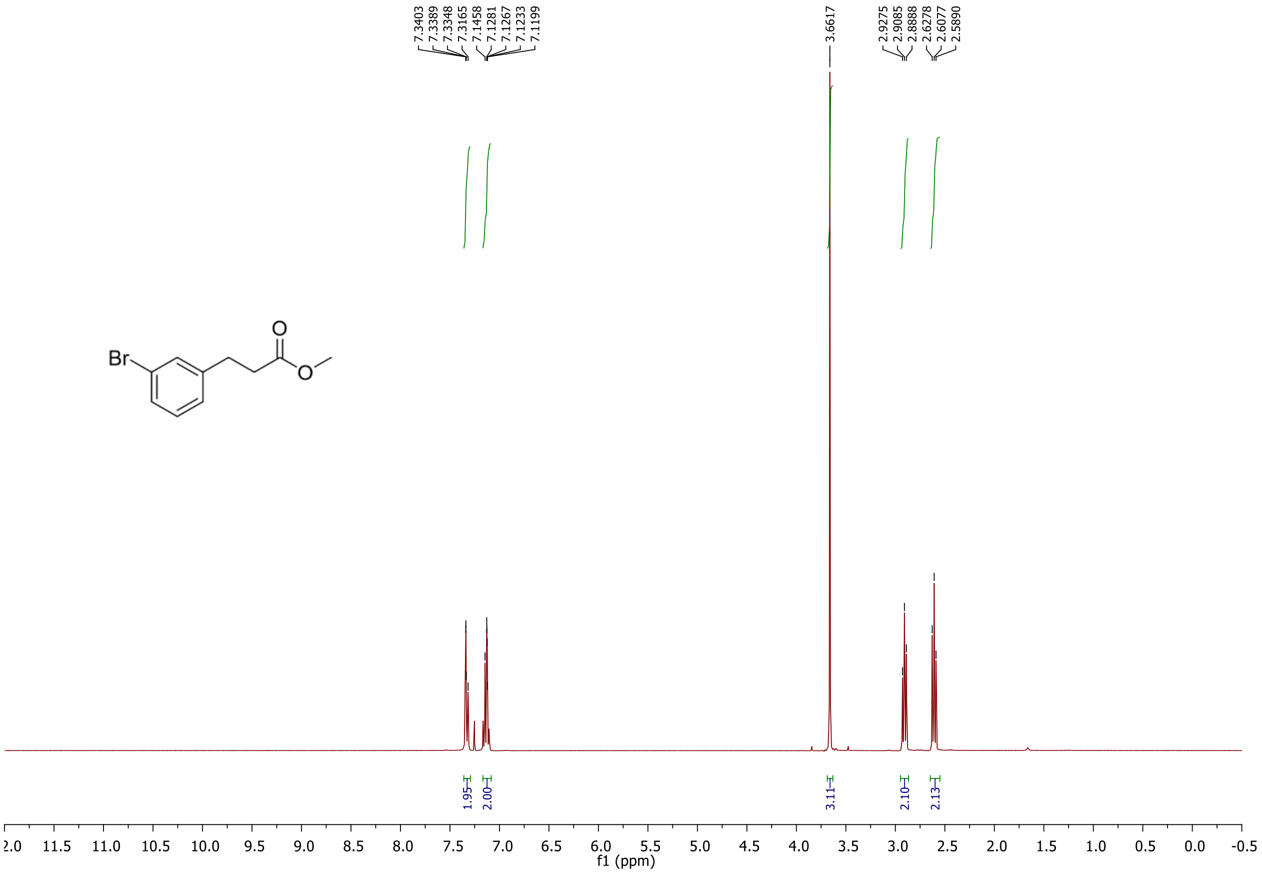

**Figure S25.** $^1$H NMR of methyl 3-(3-bromophenyl)propanoate.

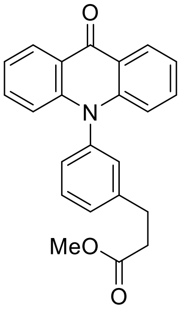

**IIb**

Synthesis of methyl 3-(3-(9-oxoacridin-10(9H)-yl)phenyl)propanoate (**IIb**)
Compound was synthesized following general procedure GP3. $^1$H NMR (400 MHz, $CDCl_3$ ) δ 8.57 (dd, J = 8.0, 1.3 Hz, 2H), 7.61 (t, J = 7.6 Hz, 1H), 7.49 (ddd, J = 8.5, 7.1, 1.5 Hz, 3H), 7.28 – 7.24 (m, 2H), 7.23 (d, J = 9.7 Hz, 1H), 7.20 (s,1H), 6.73 (d, J = 8.6 Hz, 2H), 3.62 (s, 3H), 3.06 (t, J = 7.6 Hz, 2H), 2.69 (t, J = 7.6 Hz, 2H). HRMS $[M+H]^+$ (+ TOF MS) calculated for $C_{23}H_{20}NO_3$ 358.1365 found: 358.1423

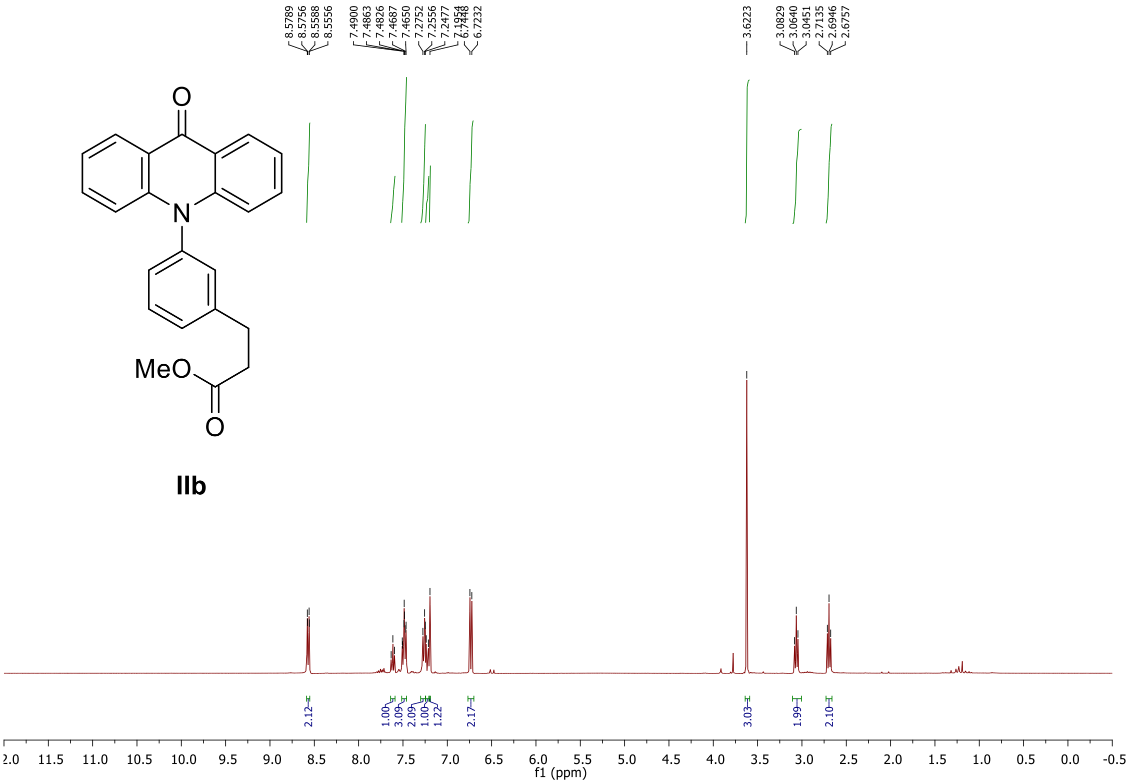


**Figure S26.** $^{1}H$ NMR of methyl 3-(3-(9-oxoacridin-10(9H)-yl)phenyl)propanoate.

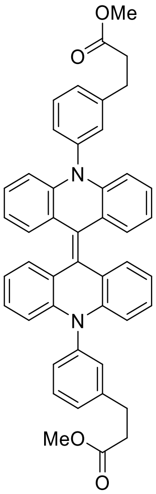


Synthesis of Dimethyl 3,3'-(10H,10'H-[9,9'-biacridinylidene]-10,10'-diylbis(3,1-phenylene))dipropionate (**IIIb**)

Acridone derivatives were dissolved in acetone and Zn (10 Equiv.) was added and stirred for 20 min at 40 ℃. Then cooled to 10 ℃ and 37-34% HCl was added dropwise, continued stirring at room temperature for a further 1 hour, and reaction progress was monitored by TLC. After completion of the reaction, diluted with 10 Equiv. of water and filtered the precipitate. Confirmed by HRMS and proceeded to the next step without further purification. HRMS (+ TOF MS) m/z calculated for $C_{46}H_{38}N_2O_4$ 341.1416 found: 341.1397

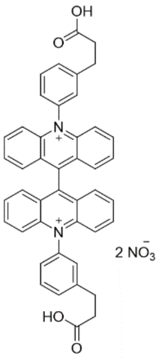


**1b**

Synthesis of 10,10'-bis(3-(2-carboxyethyl)phenyl)-[9,9'-biacridine]-10,10'-diium (**1b**)

[1]H NMR (400 MHz, DMSO-$d_6$) δ 8.31 (dd, $J$ = 11.3, 4.4 Hz, 4H), 7.98 (t, $J$ = 7.7 Hz, 2H), 7.92 – 7.81 (m, 12H), 7.78 (d, $J$ = 7.9 Hz, 2H), 7.72 (d, $J$ = 9.2 Hz, 4H), 3.10 (t, $J$ = 7.3 Hz, 4H), 2.73 (t, $J$ = 7.4 Hz, 4H). [13]C NMR (101 MHz, $CDCl_3$) δ 174.03, 152.77, 145.23, 142.60, 139.99, 137.71, 132.36, 131.82, 129.77, 129.00, 128.00, 127.03, 125.82, 120.77, 35.31, 30.63. HRMS [M] (+ TOF MS) calculated for $C_{44}H_{34}N_2O_4$ 654.2508 found: 654.2512

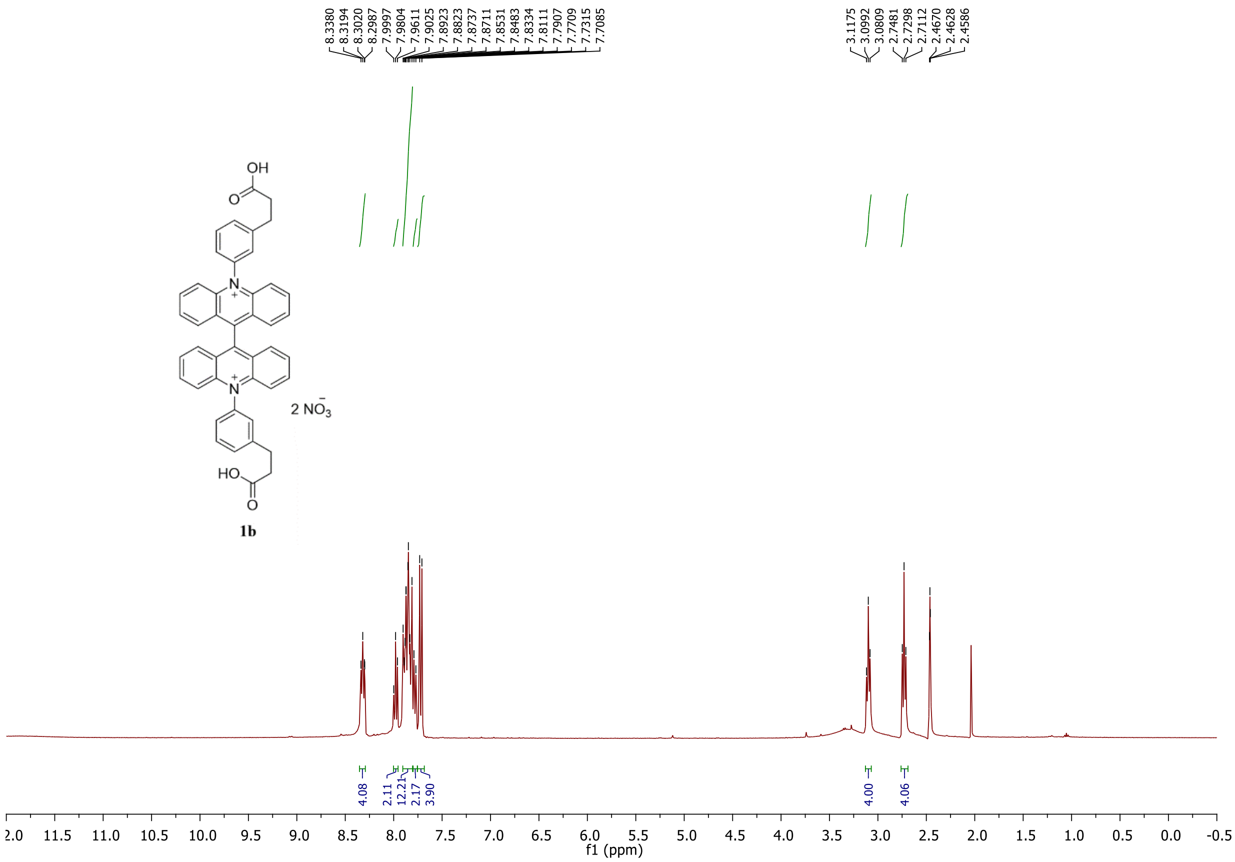


**Figure S27.** [1]H NMR of compound **1b**.

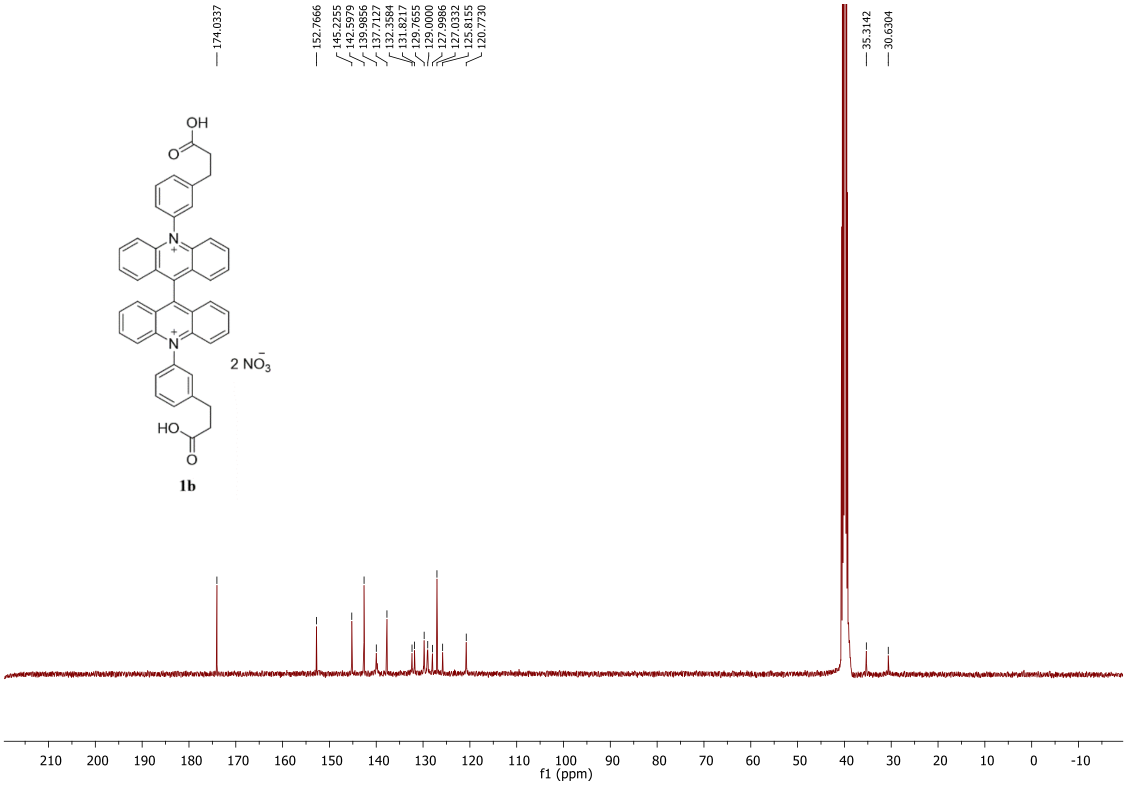


**Figure S28.** $^{13}$C NMR of compound **1b**.

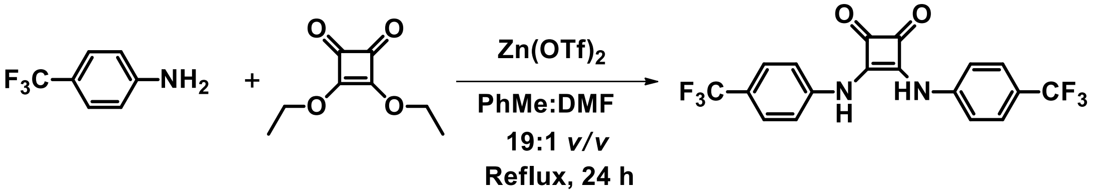


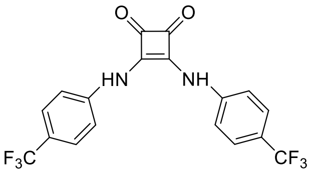

7

3,4-bis((4-(trifluoromethyl)phenyl)amino)cyclobut-3-ene-1,2-dione (**7**) was synthesized with modification according to the reported procedure in literature.[7]

In brief, 4-(trifluoromethyl)aniline (790 μL, 6.29 mmol) was added to a solution of 3,4-diethoxy-3-cyclobutene-1,2-dione (0.5 g, 2.94 mmol) in 5 mL of PhMe/DMF (*v/v* = 19:1). Then $Zn(OTf)_2$ (210 mg, 0.58 mmol, 20 mol%) was added and the mixture was refluxed overnight. The precipitate was collected by filtration and recrystallized using hot MeOH. The precipitate was collected after cooling and dried in air to give a pale yellow solid. Yield: 680 mg, 58%. $^1$HNMR (400 MHz, DMSO-d$_6$) δ 10.24 (s, 2H), 7.73 (d, *J* = 8.6 Hz, 4H), 7.63 (d, *J* = 8.6 Hz, 4H). HRMS $[M+H]^+$ (+ TOF MS) calculated for $C_{44}H_{34}N_2O_4$ 401.0719, found 401.0714.

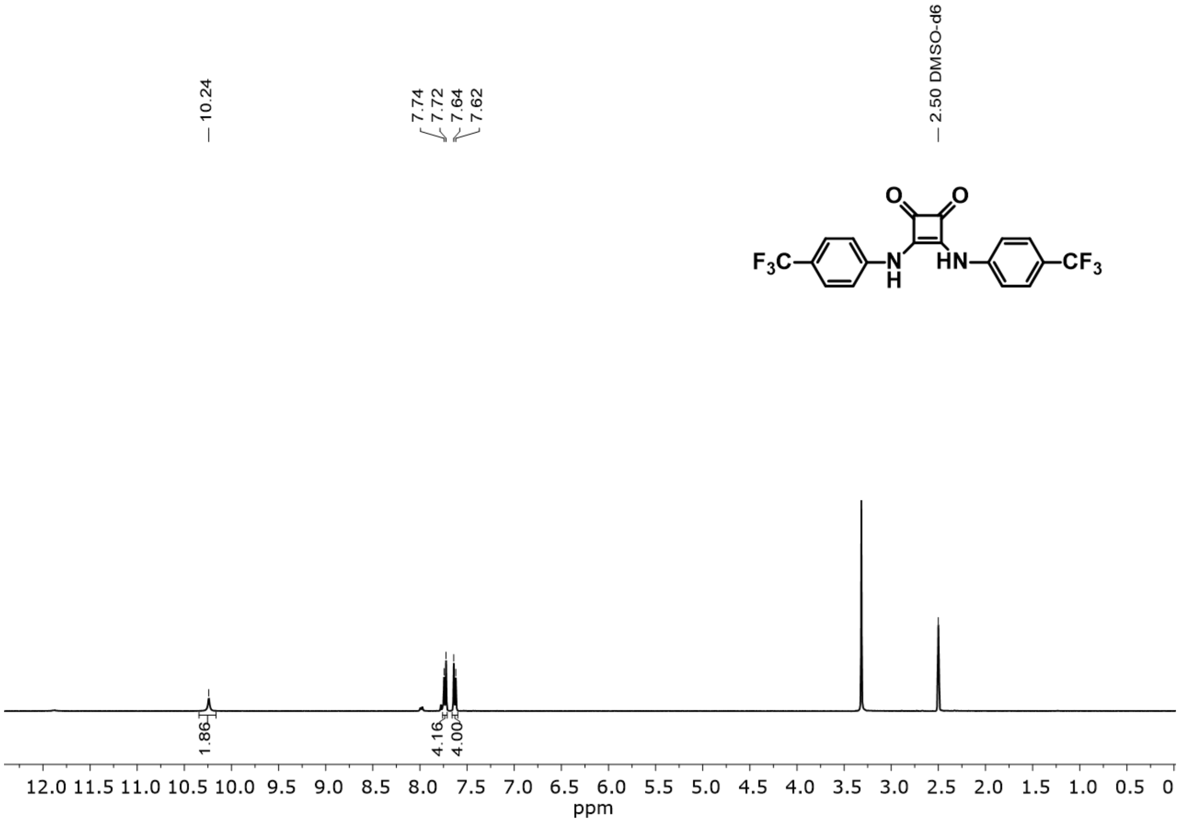


**Figure S29.** $^{1}$H NMR of compound **7**.
