## supporting information_figures for "Tunable Cytosolic Chloride Indicators for Real-Time Chloride Imaging in Live Cells"

### METHODS

#### Chemicals

NPPB, FFA, Monensin, Nigericin, Ivacaftor and CFTRinh172 were purchased from Cayman Chemicals (Ann Arbor, MI, USA). Potassium nitrate, sodium nitrate, calcium nitrate, magnesium nitrate, sodium chloride, magnesium chloride, magnesium chloride and HEPES were purchased from Sigma-Aldrich (St Louis, MA, USA). MitoView™ 650 and Lipidspot™ 650 were purchased from Biotium (Fremont, CA, USA). Lysotracker™ Deep Red, sodium gluconate, and potassium gluconate was purchased from Thermo Fisher Scientific,(CA, USA).

#### *in vitro* fluorescence measurements

UV–Vis absorption spectra was collected on a Hewlett Packard HP 8453 spectrometer and fluorescence spectra were taken using SpectraMaxTM i3/i3x multi-mode plate reader or Varian Cary eclipse fluorescence spectrophotometer. Complex **1** was dissolved in dimethyl sulfoxide (DMSO) to create a 5 mM stock solution. This stock was subsequently diluted to achieve a final concentration of 10 µM, utilizing 20 mM sodium phosphate buffer supplemented with 150 mM $KNO_3$, 5 mM $NaNO_3$, 1 mM $Ca(NO_3)_2$ and $Mg(NO_3)_2$ across a range of pH values. The emission spectra of **1** was acquired by exciting the sample at 365 nm. To study the chloride sensitivity of **1**, final [$Cl^-$] ranging between 0 mM to 150 mM was achieved by adding microlitre aliquots of 4 M KCl to the samples. To study selectivity of **1**, final concentrations of 100 mM of various salts were achieved by adding microlitre aliquots of 0.5−4 M stocks. The analysis of *in vitro* measurements for **1** was conducted by assessing the fold change, as annotated by the ratio of initial intensity ($I_0$) to final intensity ($I_F$). This is from 0 mM to 150 mM of each indicated ions, where $I_0$ is the intensity at 0 mM and $I_F$ at the respected final concentration of analyte.

#### Fluorescence lifetime measurements

Florescence lifetime spectra was collected on an Edinburgh FLS1000 spectrometer. Complex **1** was dissolved in dimethyl sulfoxide (DMSO) to create a 5 mM stock solution. This stock was subsequently diluted to achieve a final concentration of 20 µM in 5 mM sodium phosphate buffer at pH 7. For lifetime-based chloride sensitivity analysis of **1**, final [$Cl^-$] ranging between 0 mM to 100 mM was achieved by adding microlitre aliquots of 4 M KCl to each sample. Samples were excited with a pulsed laser diode (450nm) every 200 ns and decay was monitored between pulses to produce the decay spectrum. To generate τ values, decay spectrums were fit to the "ExpDecay1" function in OriginLab™ software. The subsequent analysis of lifetime (τ) measurements for **1** was conducted by assessing the fold change, as annotated by the ratio of the initial lifetime ($T_0$) to final lifetime ($T_F$). This is from 0 mM to 100 mM of each indicated ions, where $T_0$ is the intensity at 0 mM and $T_F$ at the respected final concentration of analyte.

#### Mammalian cell culture

RAW 264.7, HeLa, and HEK-293 cells were purchased from ATCC (Manassas, VA, USA). PAC-KO HeLa cells were generated previously.[1,2] These cell lines were maintained in Dulbecco's Modified Eagle's Medium (DMEM) supplemented with 10% heat-inactivated fetal bovine serum, Pen-Strep (100 U/mL-100 µg/mL). Primary Human Dermal Fibroblasts (HDF) were purchased from ATCC (Manassas, VA, USA). These cells were routinely maintained in Dulbecco's Modified Eagle's Medium/ Nutrient mixture F-12 (DMEM/F12) supplemented with 10% fetal bovine serum, Pen-Strep (100 U/mL-100 µg/mL). DMEM/F12 was purchased from (Thermo Fisher Scientific, CA, USA). DMEM and fetal bovine serum was purchased from Corning (Corning, NY, USA). All cell lines were cultured in 37 °C with 5% CO2 atmosphere.

**Wide field fluorescence imaging**

All florescence imaging studies was done using an IX-83 inverted microscope (Olympus Corporation of the Americas, Center Valley, PA, USA) and Prime BSI CMOS camera (Photometrics, USA). Filter cubes, shutter and CMOS camera were controlled using Cellsens Dimension 4.1 (Olympus Corporation of the Americas, Center Valley, PA, USA) software.

**Cellular uptake assay**

Primary Human Dermal Fibroblast (HDF) cells were plated on 35mm glass bottomed dishes (Cell Vis, Mountain view, CA,USA). The cells were pulsed with 10 or 100 μM indicated compounds in OPTI-MEM for 1 h followed by washing with PBS and incubation in cell media for 0.5 h. Indicated cell samples were treated with halide buffer (75 mM NaI, 1.25 mM KCl, 0.9 mM $CaCl_2$, 0.5 mM $MgCl_2$, 5 mM HEPES, pH = 7.4). The cells were observed using a 40×, 1.30 NA, apochromat oil immersion objective (UPlanFL N, Olympus Corporation of the Americas, Center Valley, PA, USA). BAC channel images were obtained using a filter cube containing 385/70 band pass excitation filter, ET510/80 band pass emission filter and 425 dichroic filter (Chroma, Bellows Falls, VT, USA). Background subtraction was done for all images by measuring the mean intensity over an adjacent cell free area and subtracting the value. Whole cell intensity was calculated by defining a specific ROI for each cell followed by measuring the mean intensity of each ROI using ImageJ (NIH). All Images were taken the same day and were acquired under the same acquisition settings.

**KI assay**

Primary human dermal fibroblast (HDF) cells were plated on 35mm glass bottomed dishes (Cell Vis, Mountain view, CA,USA). The cells were pulsed with 10 or 100 μM indicated compounds in OPTI-MEM for 1 h. This was followed by washing with PBS and incubation in NaI buffer diluted in cell media at indicated concentrations for 0.5h. NaI halide buffer (75 mM NaI, 1.25 mM KCl, 0.9 mM $CaCl_2$, 0.5 mM $MgCl_2$, 5 mM HEPES, pH = 7.4). The cells were observed using a 40×, 1.30 NA, apochromat oil immersion objective (UPlanFL N, Olympus Corporation of the Americas, Center Valley, PA, USA). BAC channel images were obtained using a filter cube containing 385/70 band pass excitation filter, ET510/80 band pass emission filter and 425 dichroic filter (Chroma, Bellows Falls, VT, USA). Background subtraction was done for all images by measuring the mean intensity over an adjacent cell free area and subtracting the value. Whole cell intensity was calculated by defining a specific ROI for each cell followed by measuring the mean intensity of each ROI using ImageJ (NIH). All Images were taken the same day and were acquired under the same acquisition settings.

**Cell toxicity assay**

Approximately $0.4\times10^5$ cells were seeded in 96 well culture plate overnight. The cells were treated with indicated compounds diluted to specified working concentrations in OPTI-MEM for 1 h followed by washing with PBS and then incubation in cell media for 0.5 h. The CellTiter-Blue® Cell Viability Assay was then performed according to the manufacturers' protocol (Promega, Madison, Wi, USA). Following, the plates were read using SpectraMax™ i3/i3x multi-mode plate reader.

**Chloride channel screening assay**

Primary human dermal fibroblast (HDF) cells were plated on 35mm glass bottomed dishes (Cell Vis, Mountain view, CA,USA). The cells were pretreated with 100 μM NPPB or 150 μM FFA overnight. The cells were then pulsed with 10 or 100 μM indicated compounds in OPTI-MEM with NPPB or FFA for 1 h. After overnight incubation, the cells were continuously in the presence of 100 μM NPPB or 150 μM FFA. This was followed by washing and then incubation in NaI halide buffer (75 mM NaI, 1.25 mM KCl, 0.9 mM CaCl2, 0.5 mM MgCl2, 5 mM HEPES, pH = 7.4) diluted in cell media containing NPPB or FFA for 0.5h. The cells were observed using a 40×, 1.30 NA, apochromat oil immersion objective (UPlanFL N, Olympus Corporation of the Americas, Center Valley, PA, USA). BAC channel images were obtained using a filter cube containing 385/70 band pass excitation filter, ET510/80 band pass emission filter and 425 dichroic filter (Chroma, Bellows Falls, VT, USA). Background subtraction was done for all images by measuring the mean intensity over an adjacent cell free area and subtracting the value. Whole cell intensity was calculated by defining a specific ROI for each cell followed by measuring the mean intensity of each ROI using ImageJ (NIH). All Images were taken the same day and were acquired under the same acquisition settings.

**Chloride clamping in HDF**

Primary human dermal fibroblast (HDF) cells were plated on 35mm glass bottomed dishes (Cell Vis, Mountain view, CA,USA). Following, the cells were pulsed with 10 or 100 μM indicated compounds in OPTI-MEM for 1 h. The cells were then fixed with 4% PFA for 8 min at room temperature, washed three times and retained in 1X PBS. The fixed cells were equalized by incubating in the appropriate chloride clamping buffer containing a specific concentration of chloride, 10 μM nigericin, 20 μM monesin, and 10 μM squaramide[3] (chloride ionophore) for 2.5 h at room temperature. Following, cells were observed using a 40×, 1.30 NA, apochromat oil immersion objective (UPlanFL N, Olympus Corporation of the Americas, Center Valley, PA, USA). BAC channel images were obtained using a filter cube containing 385/70 band pass excitation filter, ET510/80 band pass emission filter and 425 dichroic filter (Chroma, Bellows Falls, VT, USA). Background subtraction was done for all images by measuring the mean intensity over an adjacent cell free area and subtracting the value. Whole cell intensity was calculated by defining a specific ROI for each cell followed by measuring the mean intensity of each ROI using ImageJ (NIH). All Images were taken the same day and were acquired under the same acquisition settings. The chloride clamping buffers containing different chloride concentrations were prepared by mixing chloride positive buffer (120 mM KCl, 20 mM NaCl, 1 mM $CaCl_2$, 1 mM $MgCl_2$, 20 mM HEPES, pH, 7.0) and chloride negative buffer (120 mM $KNO_3$, 20 mM $NaNO_3$, 1 mM $Ca(NO_3)_2$, 1 mM $Mg(NO_3)_2$, 20 mM HEPES, pH 7.0) in different ratios.

**CFTR modulation**

Primary human dermal fibroblast (HDF) cells were plated on 35mm glass bottomed dishes (Cell Vis, Mountain view, CA,USA). The cells were pretreated with 2 μM Ivacaftor or 20 μM CFTRinh172 overnight. The cells were pulsed with 100 μM indicated compounds in OPTI-MEM with Ivacaftor or CFTRinh172 for 1 h. This was followed by washing with PBS and then incubation in cell media for 0.5 h. After overnight incubation, the cells were continuously in the presence of 2 μM ivacaftor or 20 μM CFTRinh172.Following, cells were observed using a 40×, 1.30 NA, apochromat oil immersion objective (UPlanFL N, Olympus Corporation of the Americas, Center Valley, PA, USA). BAC channel images were obtained using a filter cube containing 385/70 band pass excitation filter, ET510/80 band pass emission filter and 425 dichroic filter (Chroma, Bellows Falls, VT, USA). Background subtraction was done for all images by measuring the mean intensity over an adjacent cell free area and subtracting the value. Whole cell intensity was calculated by defining a specific ROI for each cell followed by measuring the mean intensity of each ROI using ImageJ (NIH). All Images were taken the same day and were acquired under the same acquisition settings.

**Cellular distribution assay**

Primary human dermal fibroblasts (HDF) or RAW 264.7 macrophages were plated on 35mm glass bottomed dishes (Cell Vis, Mountain view, CA,USA). The cells were pulsed with 100 μM indicated compounds in OPTI-MEM for 1 h. This was followed by washing with PBS and then incubation in cell media for 0.5 h. The cells were subsequently treated with MitoView650, Lipid650, or Lysotracker deep red, each according to their manufacturer's protocol. Following, cells were observed using a 40×, 1.30 NA, apochromat oil immersion objective (UPlanFL N, Olympus Corporation of the Americas, Center Valley, PA, USA). BAC channel images were obtained using a filter cube containing 385/70 band pass excitation filter, ET510/80 band pass emission filter and 425 dichroic filter (Chroma, Bellows Falls, VT, USA). MitoView650, Lipid650, or Lysotracker deep red channel images were obtained using a filter cube containing 620/60 band pass excitation filter, 700/75 band pass emission filter and T660lpxr dichroic filter (Chroma, Bellows Falls, VT, USA). Background subtraction was done for all images by measuring the mean intensity over an adjacent cell free area and subtracting the value.

**Real time: chloride channel screening assay**

Primary human dermal fibroblast (HDF) cells were plated on 35mm glass bottomed dishes (Cell Vis, Mountain view, CA,USA). The cells were pretreated with 150 μM FFA overnight. The cells were pulsed with 100 μM indicated compounds in OPTI-MEM with FFA for 1 h. After overnight incubation, the cells were continuously in the presence of 150 μM FFA if stated. After washing, cells were incubated with 500 μL cell media for 0.5 h at 37 C. Subsequently, cells were placed at room temp and observed using a 40×, 1.30 NA, apochromat oil immersion objective (UPlanFL N, Olympus Corporation of the Americas, Center Valley, PA, USA). BAC channel images were obtained using a filter cube containing 385/70 band pass excitation filter, ET510/80 band pass emission filter and 425 dichroic filter (Chroma, Bellows Falls, VT, USA). After establishing a baseline intensity (approx. 10-20 seconds in each sample) 500 μL of 2x NaI halide buffer was added, and images were taken every 1 s for 5 minutes to view the real time response to Iodide at room temperature. Whole cell intensity values were calculated by defining a specific ROI for each cell and then measuring intensity at different time intervals using ImageJ

(NIH). All Images were taken the same day and were acquired under the same acquisition settings. DMSO was used as a vehicle control.

**Chloride substitution in HeLa cells**

HeLa cells were plated on 35mm glass bottomed dishes (Cell Vis, Mountain view, CA,USA). The cells were pulsed with 100 µM indicated compounds in OPTI-MEM for 1 h. Following, cells were incubated in Chloride or Gluconate buffers for 3 h at 37 C. The cells were then observed using a 60×, 1.42 NA, apochromat oil immersion objective (UPlanFL N, Olympus Corporation of the Americas, Center Valley, PA, USA). BAC channel images were obtained using a filter cube containing 385/70 band pass excitation filter, ET510/80 band pass emission filter and 425 dichroic filter (Chroma, Bellows Falls, VT, USA). Background subtraction was done for all images by measuring the mean intensity over an adjacent cell free area and subtracting the value. Whole cell intensity was calculated by defining a specific ROI for each cell followed by measuring the mean intensity of each ROI using ImageJ (NIH). All Images were taken the same day and were acquired under the same acquisition settings. The chloride buffers contained (145mM NaCl/5 mM KCl, 10 mM HEPES (pH 7.2), and 10mM Glucose). The gluconate buffers contained (145mM NaGluconate/5 mM KGluconate, 10 mM HEPES (pH 7.2), and 10mM Glucose).

**Real time imaging: chloride flux in HeLa cells**

HeLa cells were plated on 35mm glass bottomed dishes (Cell Vis, Mountain view, CA,USA). The cells were pulsed with 100 µM indicated compounds in OPTI-MEM for 1h. Following, cells were incubated in Gluconate buffers for 3h at 37 C. After establishing a baseline intensity (approx.10-15 seconds in each sample) cells were stimulated with a Chloride buffer and fluorescence change was monitored. The cells were then observed using a 60×, 1.42 NA, apochromat oil immersion objective (UPlanFL N, Olympus Corporation of the Americas, Center Valley, PA, USA). BAC channel images were obtained using a filter cube containing 385/70 band pass excitation filter, ET510/80 band pass emission filter and 425 dichroic filter (Chroma, Bellows Falls, VT, USA). Whole cell intensity values were calculated by defining a specific ROI for each cell and then measuring intensity at different time intervals using ImageJ (NIH). All Images were taken the same day and were acquired under the same acquisition settings. The chloride buffers contained (145mM NaCl/5 mM KCl, 10 mM HEPES (pH 7.2), and 10mM Glucose). The gluconate buffers contained (145mM NaGluconate/5 mM KGluconate, 10 mM HEPES (pH 7.2), and 10mM Glucose).

**Real time imaging: PAC flux assay**

HeLa or HeLa PAC-KO cells were plated on 35mm glass bottomed dishes (Cell Vis, Mountain view, CA,USA). The cells were pulsed with 100 µM indicated compounds in OPTI-MEM for 1h and cells were washed with PBS. Following, cells were incubated in chloride buffer (pH 7) for 10 mins prior to imaging. The cells were then observed using a 60×, 1.42 NA, apochromat oil immersion objective (UPlanFL N, Olympus Corporation of the Americas, Center Valley, PA, USA). BAC channel images were obtained using a filter cube containing 385/70 band pass excitation filter, ET510/80 band pass emission filter and 425 dichroic filter (Chroma, Bellows Falls, VT, USA). After establishing a baseline intensity (approx.10-15 seconds in each sample) cells were stimulated with acidic buffer containing 10 mM NaI and fluorescence change was monitored. Whole cell intensity values were calculated by defining a specific ROI for each cell and then measuring intensity at different time intervals using ImageJ (NIH). All Images were taken the same day and were acquired under the same acquisition settings. The chloride buffer contained (mM) :140 NaCl, 5 KCl, 1 $CaCl_2$, 1 $MgCl_2$, 10 HEPES (pH 7). Each pH was adjusted using a citric acid buffer (pH 3.2) to get desired acidic pH and supplemented with 10 mM NaI if stated.

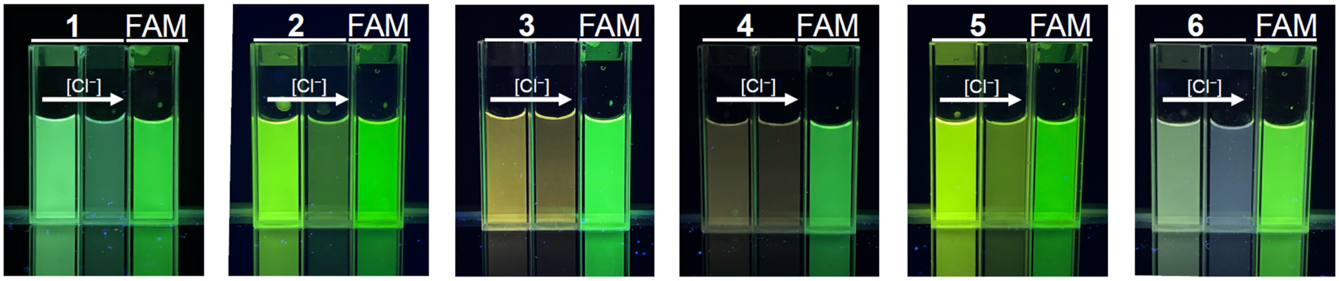


**Figure S1 Chloride sensitive fluorophores 1–6**. Images of 5 μM fluorescein (FAM) and **1–6** in 5 mM UB4 buffer (pH = 7.2) in the presence and absence of 100 mM potassium chloride under UV illumination.

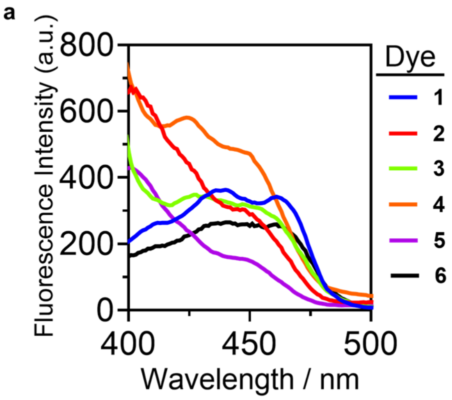


**Figure S2.** Excitation scans of **1–6** in 5 mM UB4 buffer (pH = 7.2).

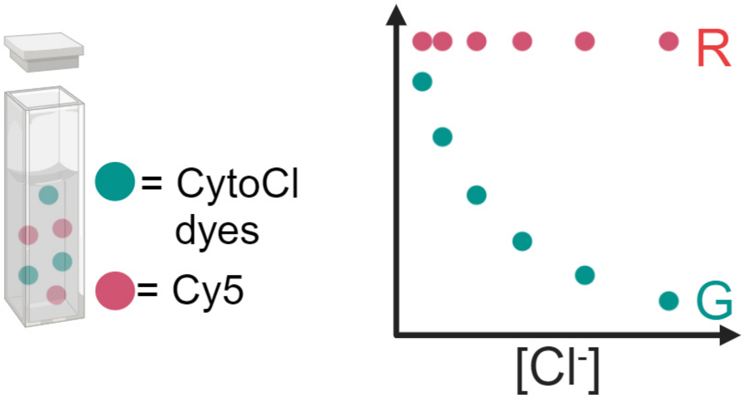


**Figure S3 Schematic diagram illustrating the working principle of *in vitro* characterization**. A stock mixture contains **1–6** and Cyanine5. This mixture was added to solutions with varying pH and [Cl⁻]. Cyanine5 serves as a reference unit because it is insensitive to pH and chloride. Utilizing this system, the concentration ratio between the tested compound (**1–6**) and Cyanine5 remains constant. It creates a fluorescence intensity ratio (G/R) or (R/G) between each compound (**1–6**, G) and Cyanine5 (R). By plotting (R/G) against [Cl⁻] we can generate Stern-Volmer plots and monitor Stern-Volmer constants.

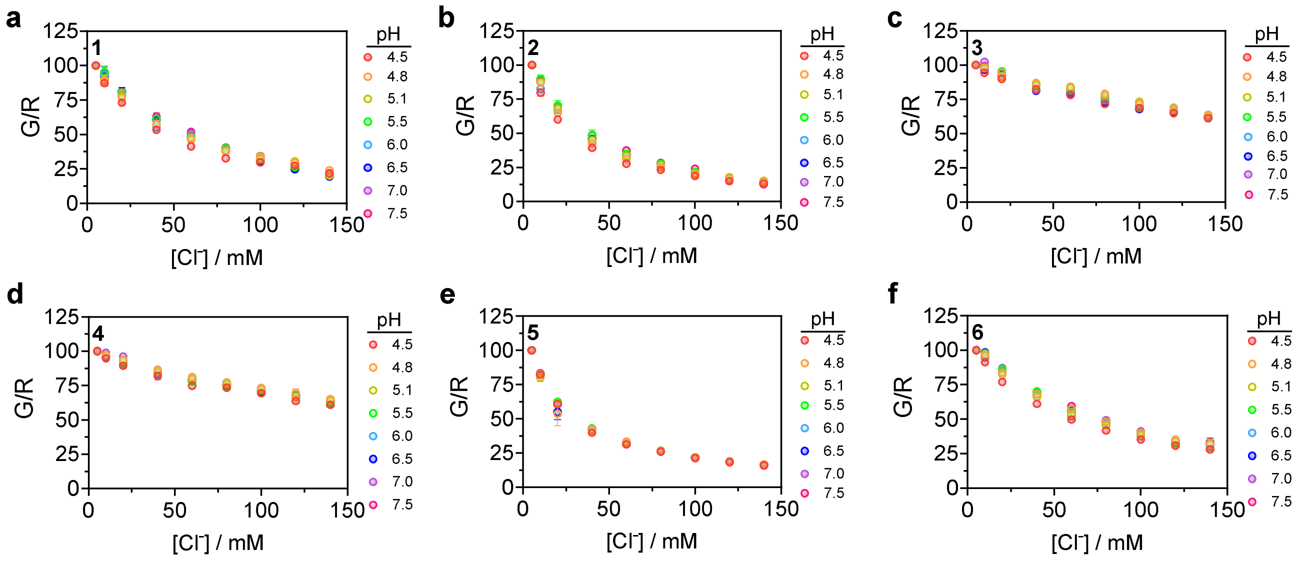


**Figure S4 *In vitro* characterization of 1–6**. Normalized fluorescence intensity ratio (G/R) of **1–6** (G) and Cyanine5 (R) as a function of $Cl^-$ concentration at the indicated pH value. Values were normalized to the G/R at 5 mM $Cl^-$. Experiments were performed in triplicate. Error bars indicate the mean ± standard error of the mean (s.e.m.) of three independent measurements.

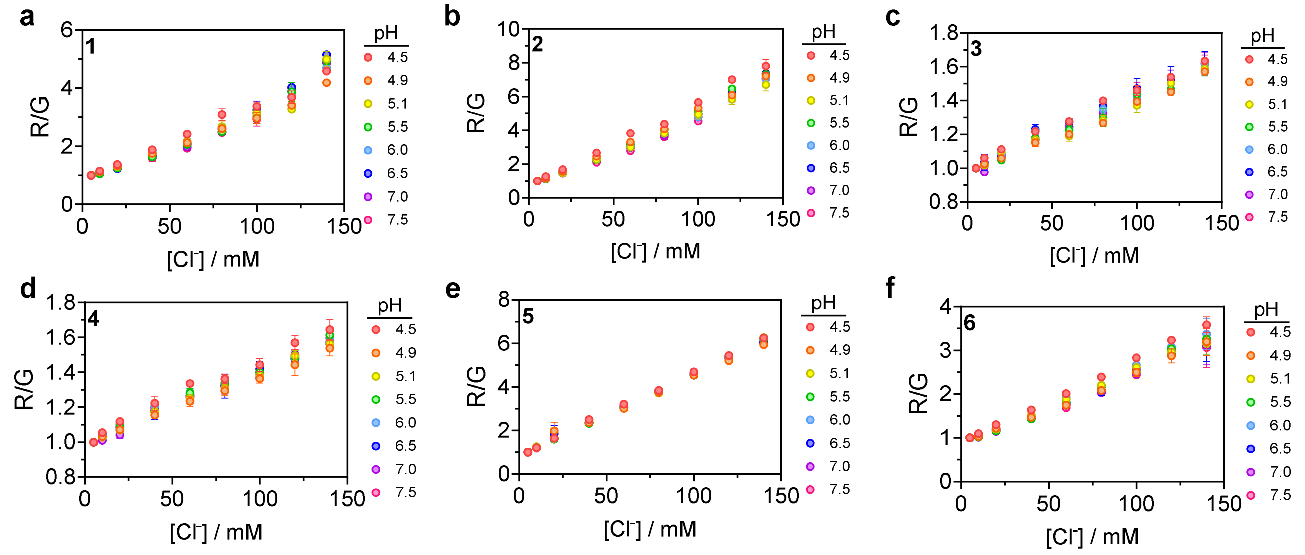


**Figure S5 Stern–Volmer plot of the fluorescence-quenching of 1–6 by chloride**. Normalized fluorescence intensity ratio (R/G) of Cyanine5 (R) and **1–6** (G) as a function of $Cl^-$ concentration at the indicated pH value. Values were normalized to the R/G at 5 mM $Cl^-$. Experiments were performed in triplicate. Error bars indicate the mean ± standard error of the mean (s.e.m.) of three independent measurements.

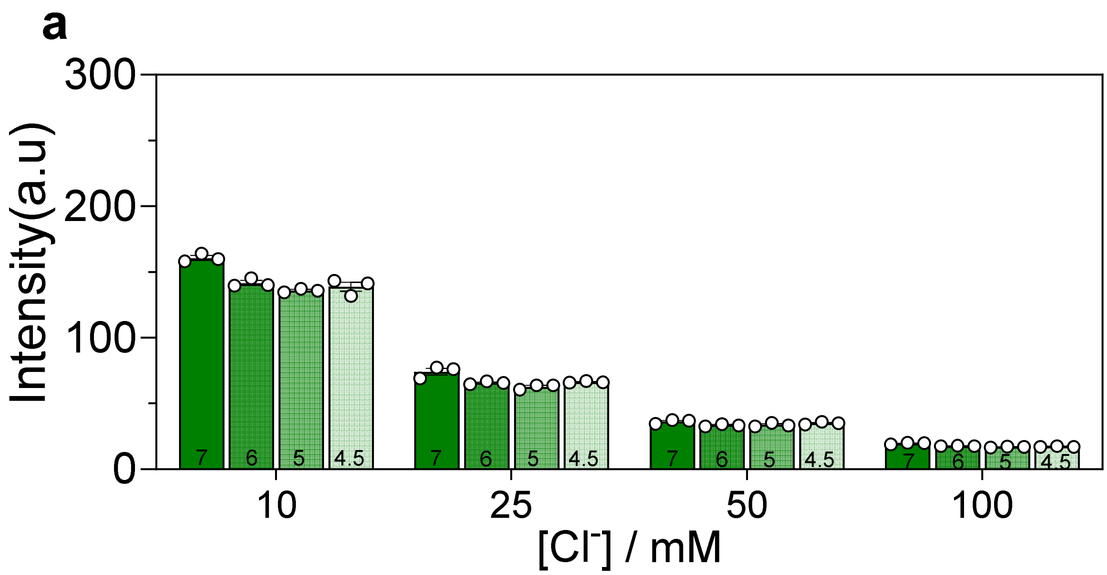


**Figure S6 pH sensitivity profile of 1.** Fluorescence intensity of **1** in 5 mM sodium phosphate buffer at pH 4.5, 5, 6, and 7. Experiments were performed in triplicate. Error bars indicate the mean ± standard error of the mean (s.e.m.) of three independent measurements.

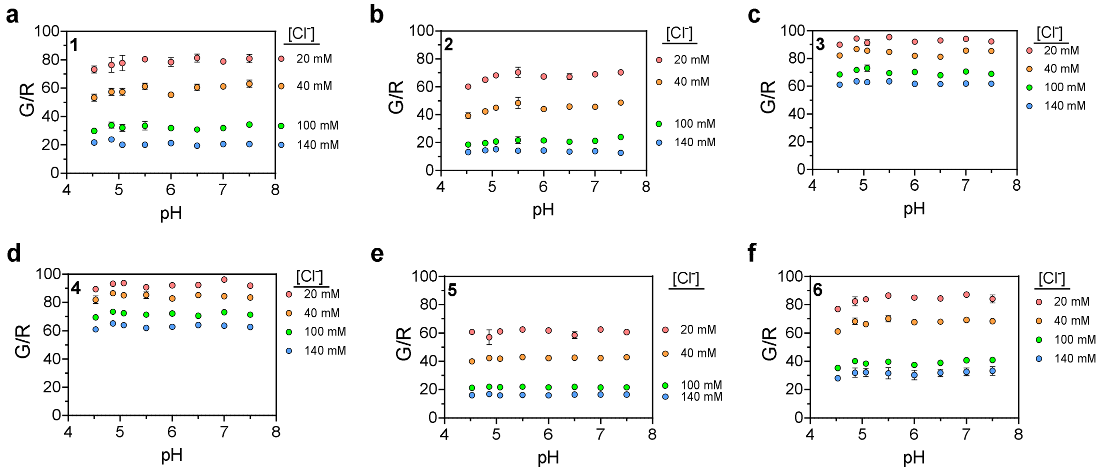


**Figure S7 pH profiles of 1–6.** Normalized fluorescence intensity ratio (G/R) of **1–6** (G) and Cyanine5 (R) in the presence of 20 mM, 40 mM, 100 mM, and 140 mM chloride in UB4 buffer (5 mM HEPES, MES and sodium acetate, 150 mM $KNO_3$, 5 mM $NaNO_3$, 1 mM $Ca(NO_3)_2$ and $Mg(NO_3)_2$) at indicated pH. Values were normalized to the G/R at 5 mM $Cl^-$. Experiments were performed in triplicate. Error bars indicate the mean ± standard error of the mean (s.e.m.) of three independent measurements.

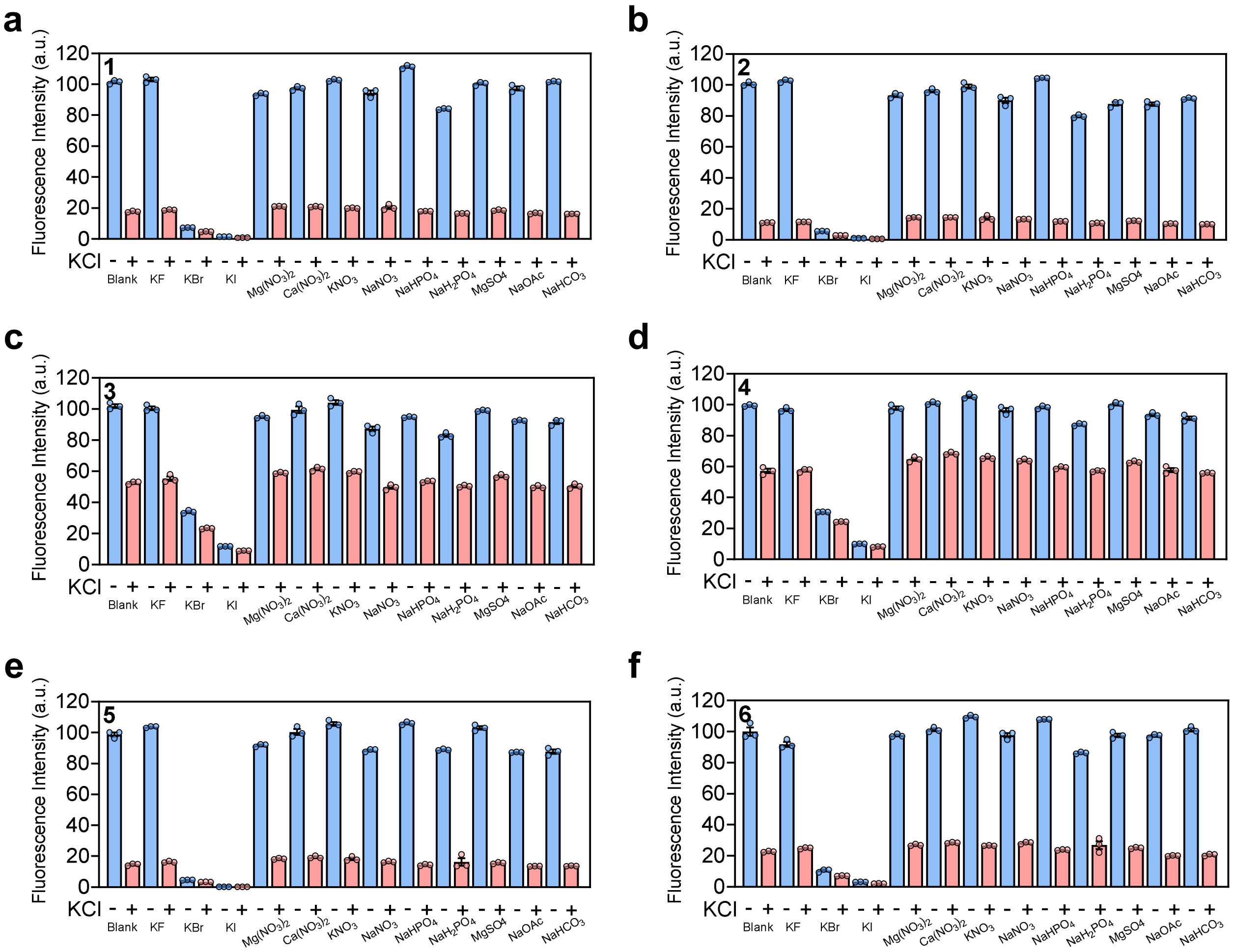


**Figure S8 Selectivity profiles of 1–6.** The fluorescence intensities of **1–6** in the presence of various anions (100 mM KF, KBr, KI, $Mg(NO_3)_2$, $Ca(NO_3)_2$, $KNO_3$, $NaNO_3$, $NaHPO_4$, $NaH_2PO_4$, $MgSO_4$, 5 mM NaOAc, 1 mM $NaHCO_3$) with and without 100 mM KCl in 5 mM UB4 buffer (pH = 7.0). Error bars indicate the mean ± standard error of the mean (s.e.m.) of three independent measurements.

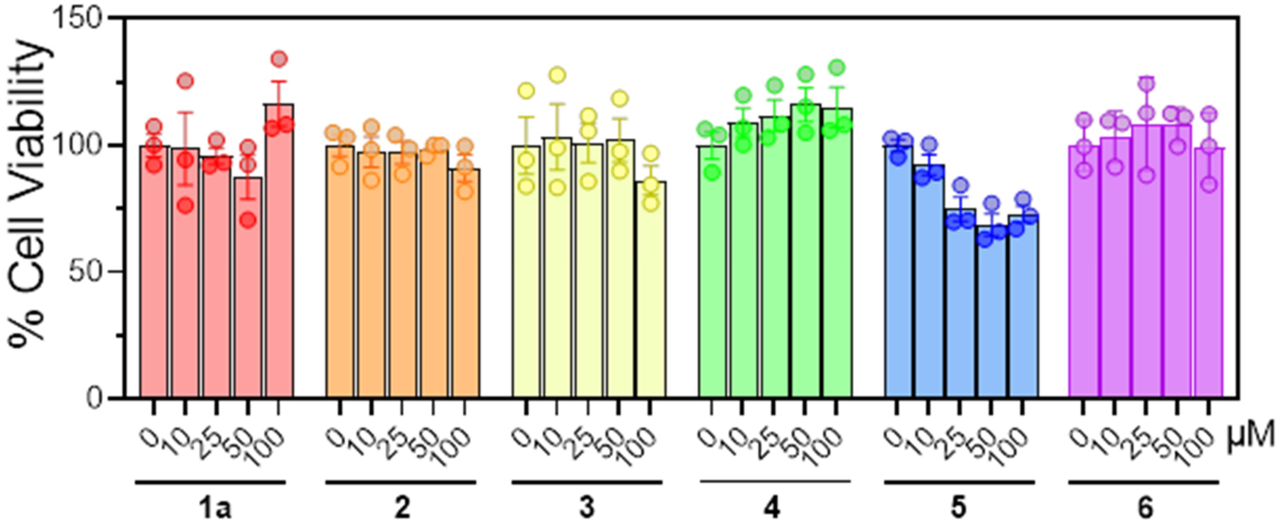


**Figure S9 Cell toxicity of 1–6.** Cell viability of HDF cells upon labeling of **1–6.** Error bars indicate the mean ± standard error of the mean (s.e.m.) of three independent measurements.

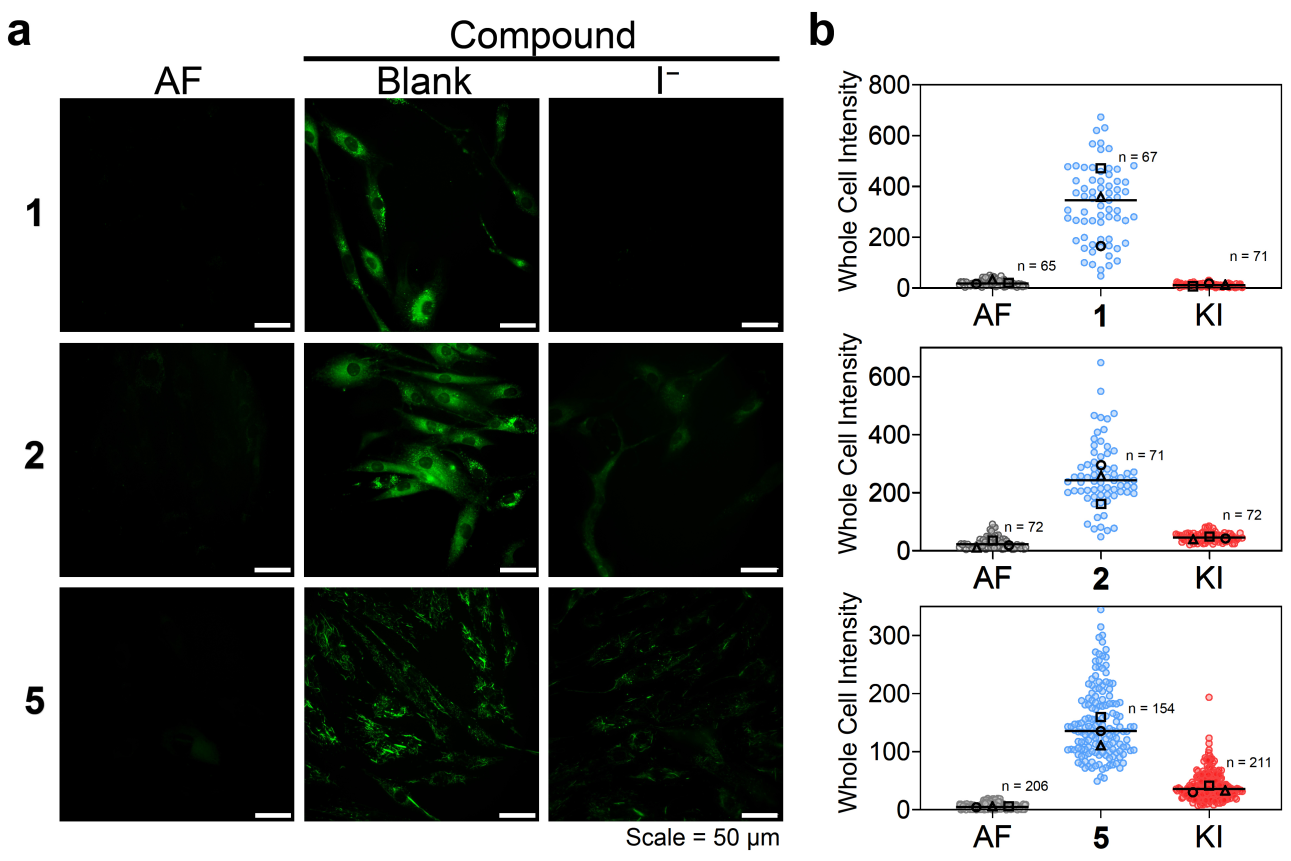


**Figure S10 Cellular uptake of CytoCl dyes. a** Representative fluorescence images of HDF cells labeled with **1–2** and **5**, with and without incubation of 75 mM iodide. AF, autofluorescence. **b**, Quantification of whole cell intensity in indicated conditions. Experiments were performed in triplicate. The median value of each trial is given by a square, circle and triangle symbol (n = number of cells).

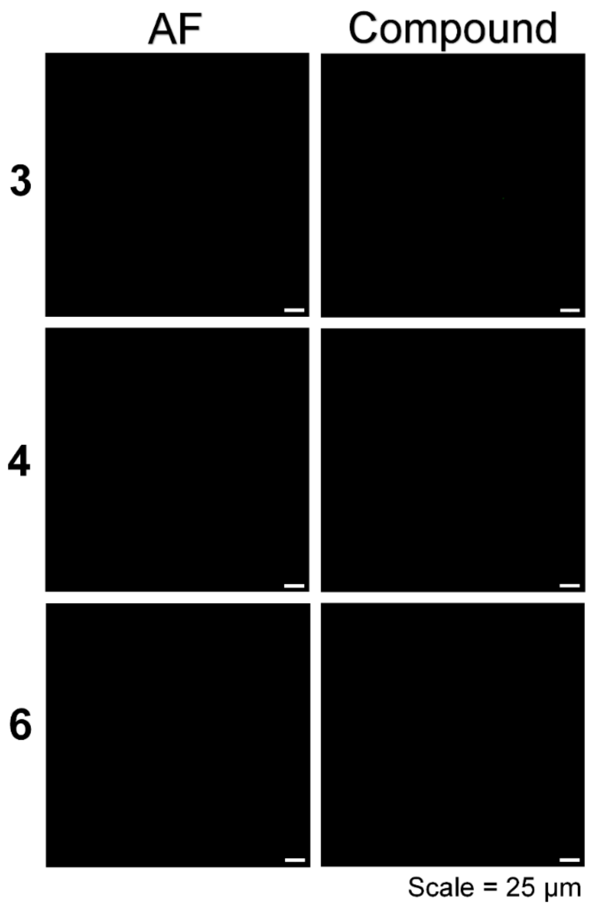


**Figure S11 Cellular uptake of CytoCl dyes.** Representative fluorescence images of HDF cells labeled with **3**–**4** and **6**. AF, autofluorescence. Experiments were performed in triplicate.

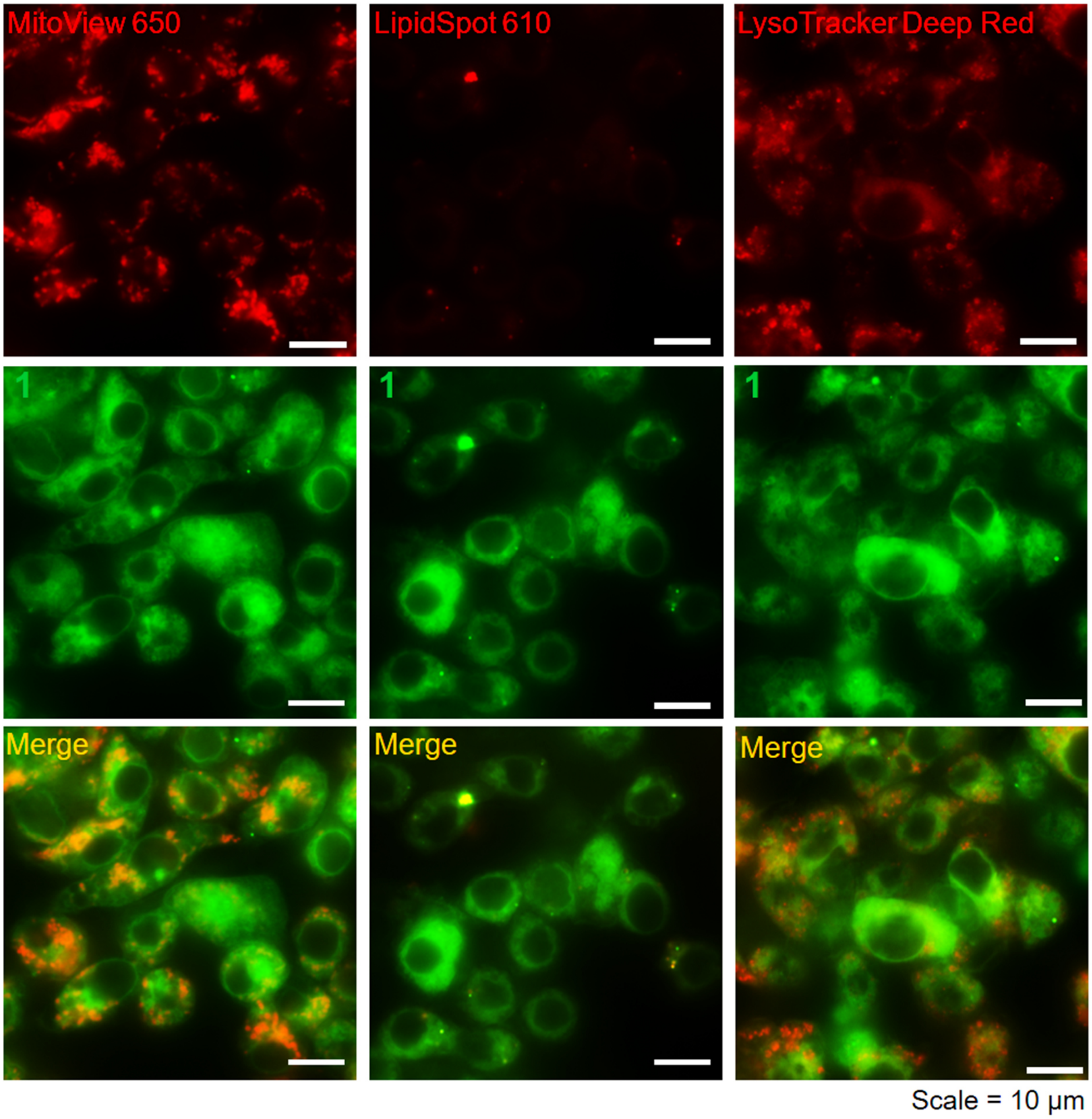


**Figure S12 Cellular distribution of 1 in RAW 264.7 cells.** Representative fluorescence images of RAW 264.7 cells labelled with **1** (green) and organelle markers (red, MitoView 650, LipidSpot610, or LysoTracker deep red). Experiments were performed in triplicate.

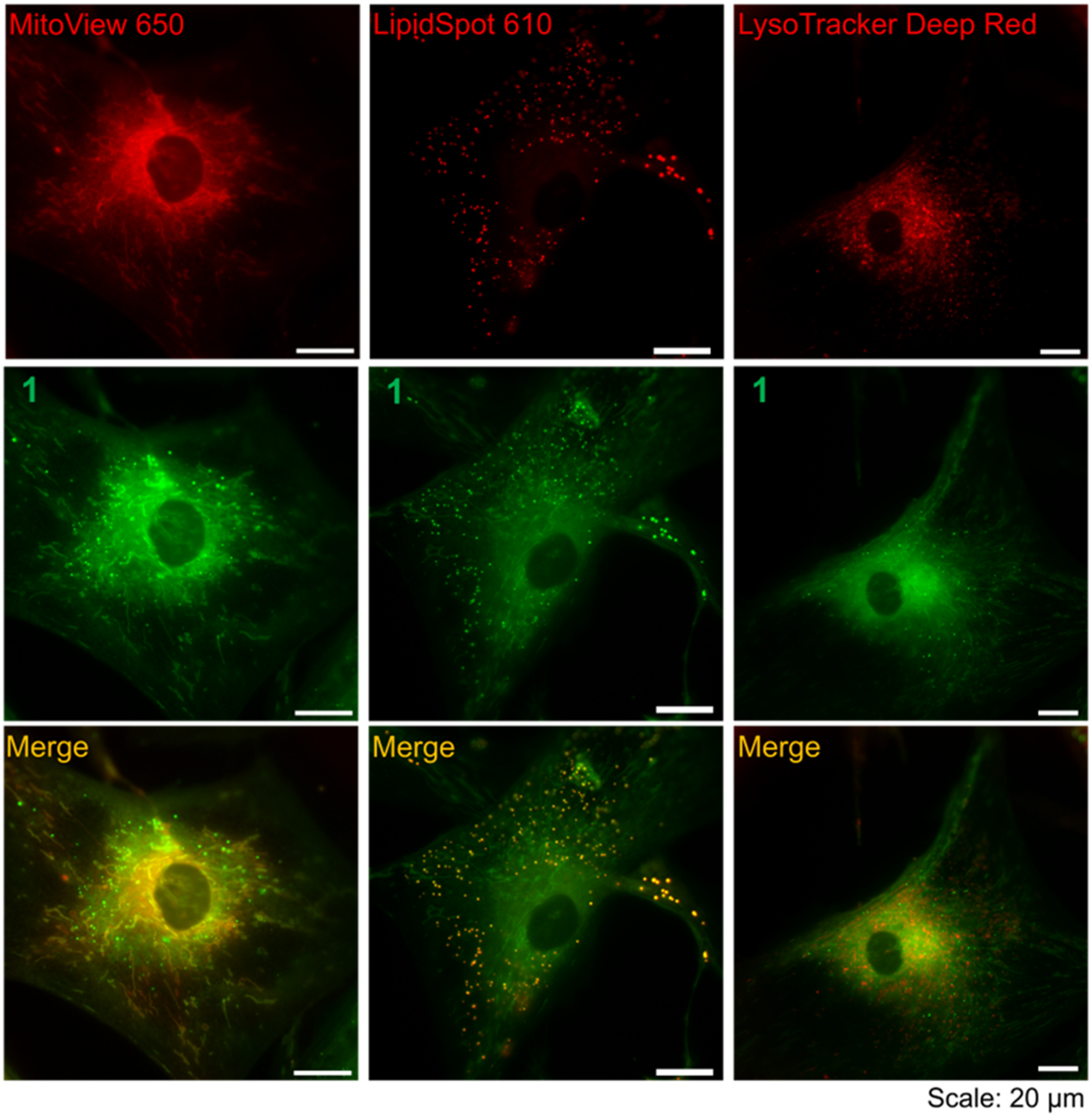


**Figure S13 Cellular distribution of 1 in HDF cells.** Representative fluorescence images of HDF cells labelled with **1** (green) and organelle markers (red, MitoView 650, LipidSpot610, or LysoTracker deep red). Experiments were performed in triplicate.

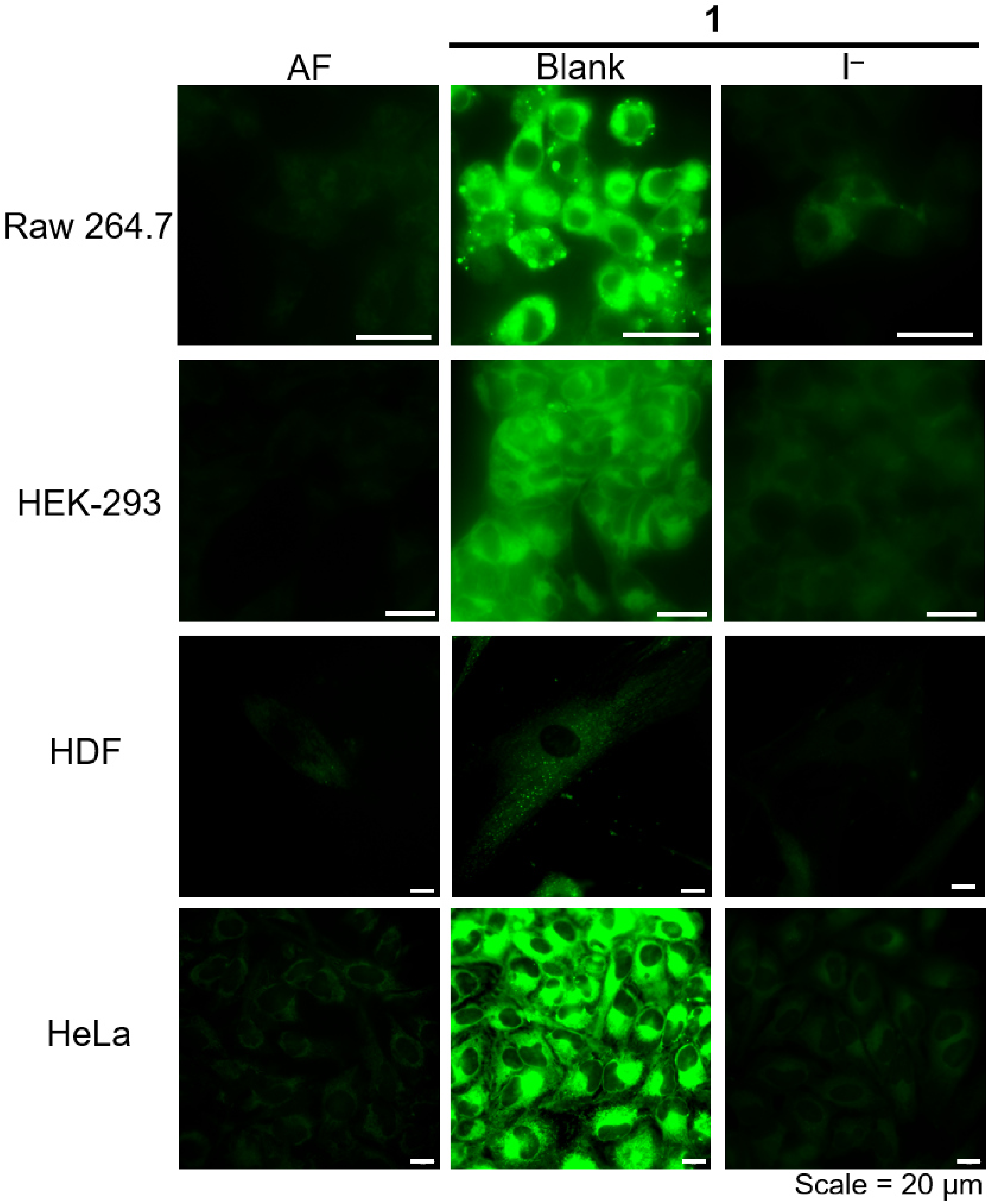


**Figure S14. Cellular uptake of 1 in different cell types.** Representative fluorescence images of **1**-labeled cells, with and without incubation of 75 mM iodide. AF, autofluorescence. Experiments were performed in triplicate.

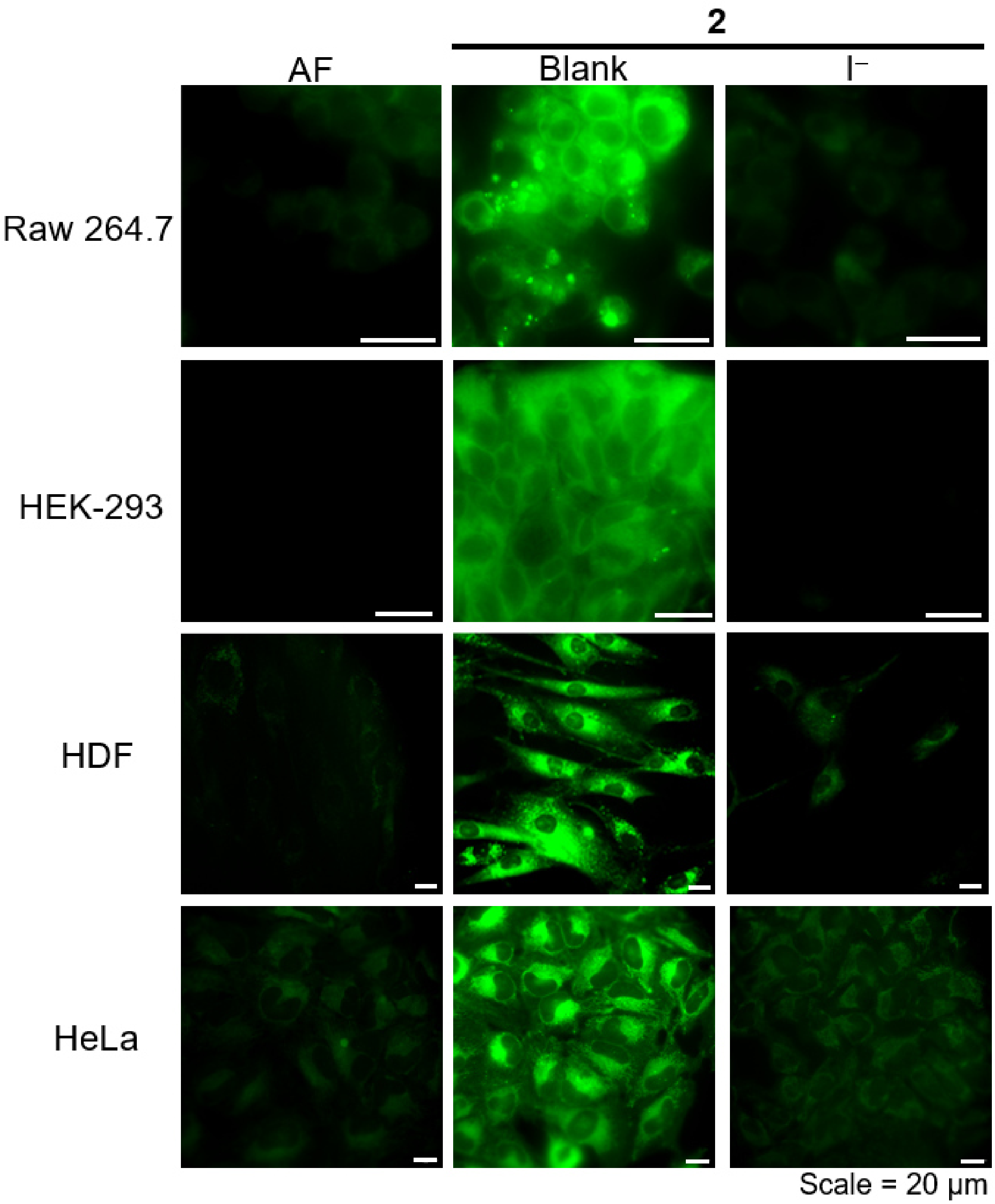


**Figure S15. Cellular uptake of 2 in different cell types.** Representative fluorescence images of **2**-labeled cells, with and without incubation of 75 mM iodide. AF, autofluorescence. Experiments were performed in triplicate.

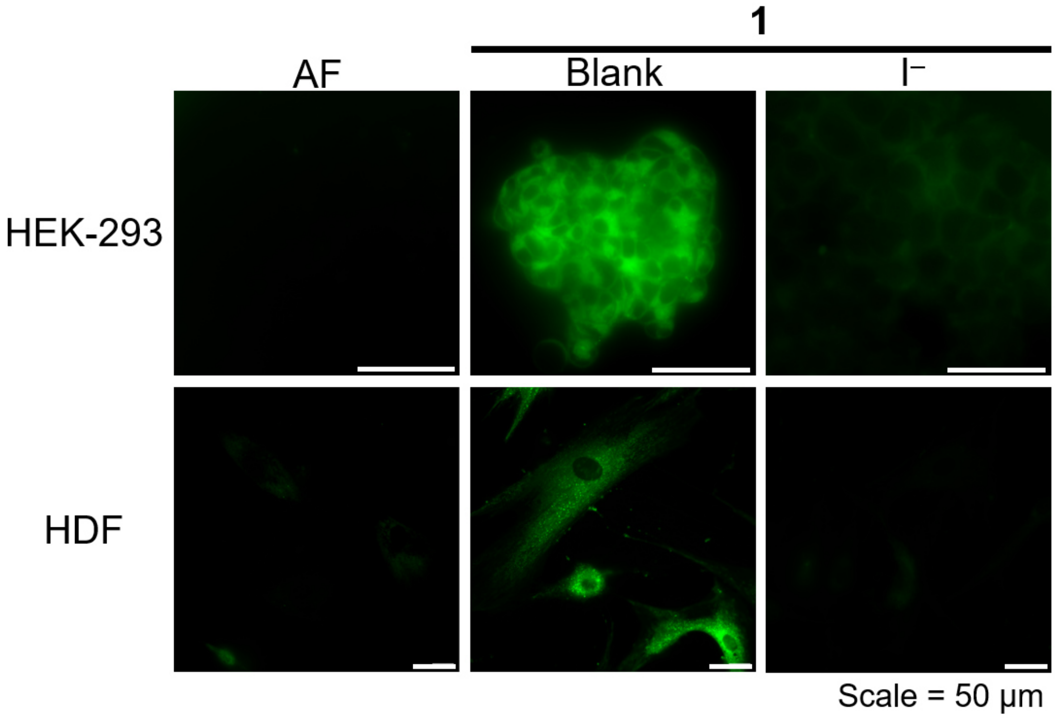


**Figure S16 Imaging 1-labeled cells using GFP channel.** Representative fluorescence images (in GFP channel) of HEK-293 and HDF cells labeled with **1**, with and without incubation of 75 mM iodide. AF, autofluorescence. Experiments were performed in triplicate.

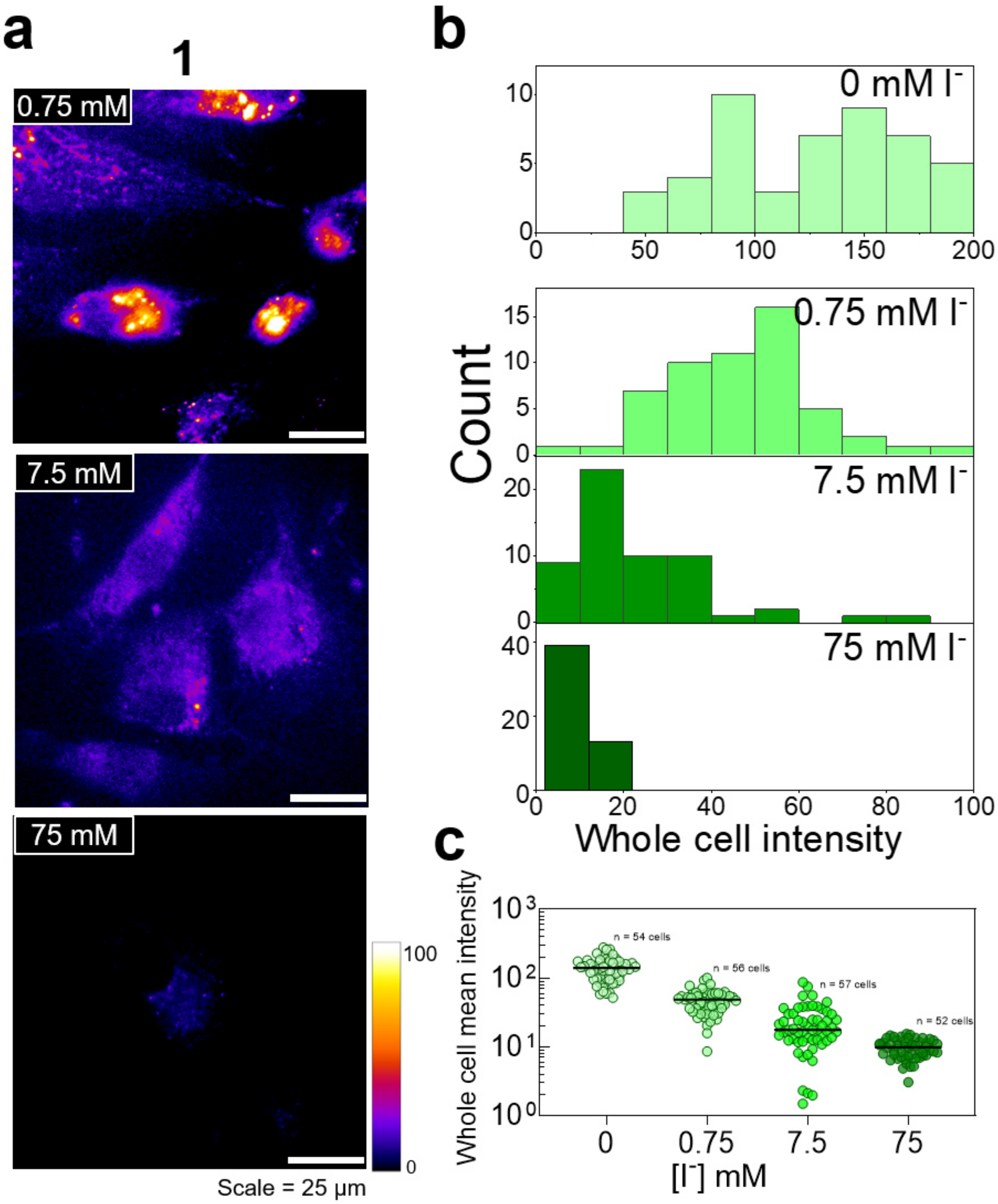


**Figure S17 Intracellular iodide sensing characteristics of 1. a,** Representative fluorescence images of **1**-labeled HDF cells upon incubation with 0 mM, 0.75 mM, 7.5 mM, and 75 mM of $I^-$. **b**,**c**, Quantification of whole cell intensity in indicated conditions. Fluorescence intensity of **1**-labeled cell decreased with increasing [$I^-$]. Experiments were performed in triplicate (n = number of cells).

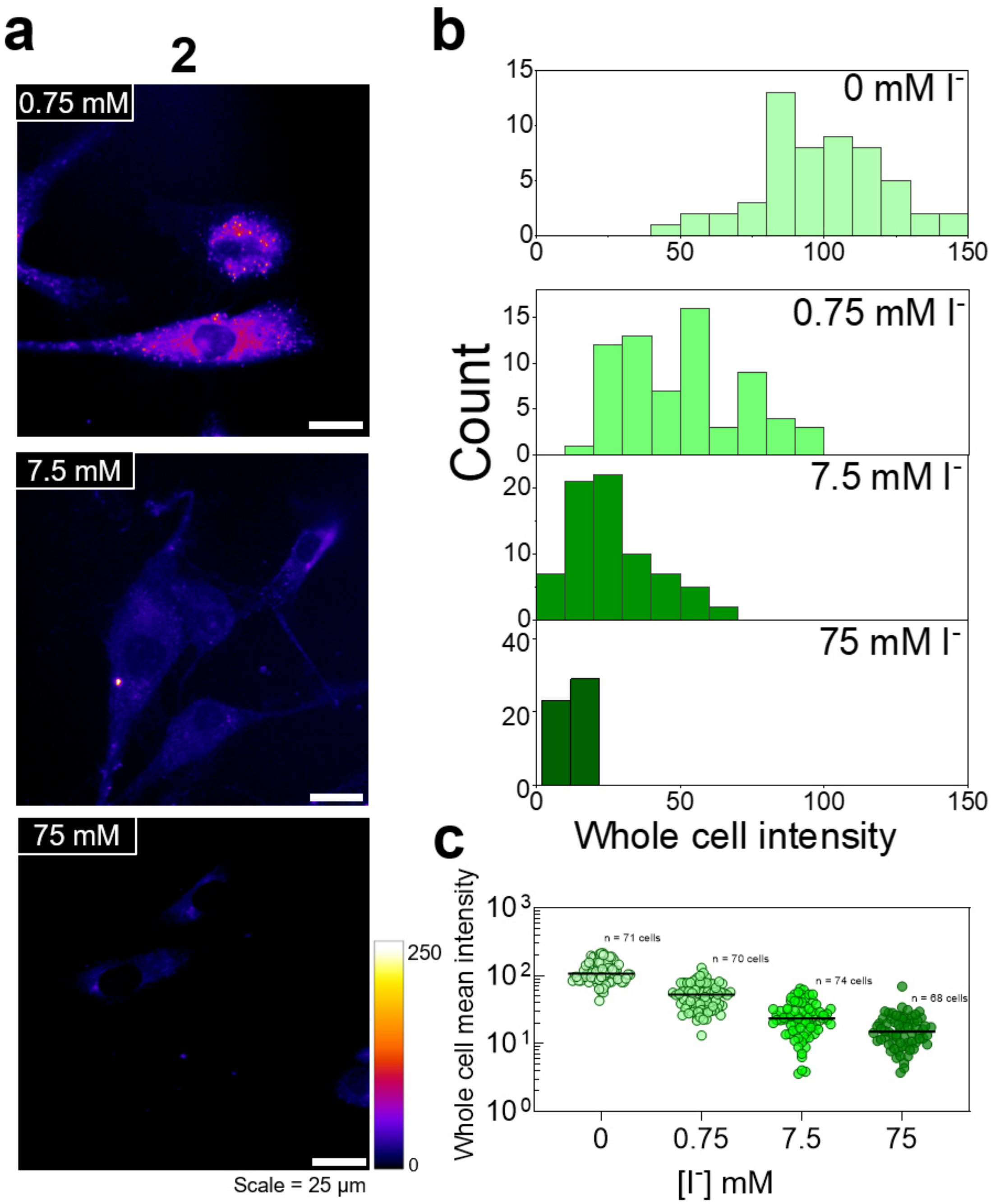


**Figure S18 Intracellular iodide sensing characteristics of 2. a,** Representative fluorescence images of **2**-labeled HDF cells upon incubation with 0 mM, 0.75 mM, 7.5 mM, and 75 mM of $I^-$. **b**,**c**, Quantification of whole cell intensity in indicated conditions. Fluorescence intensity of **2**-labeled cell decreased with increasing [$I^-$]. Experiments were performed in triplicate (n = number of cells).

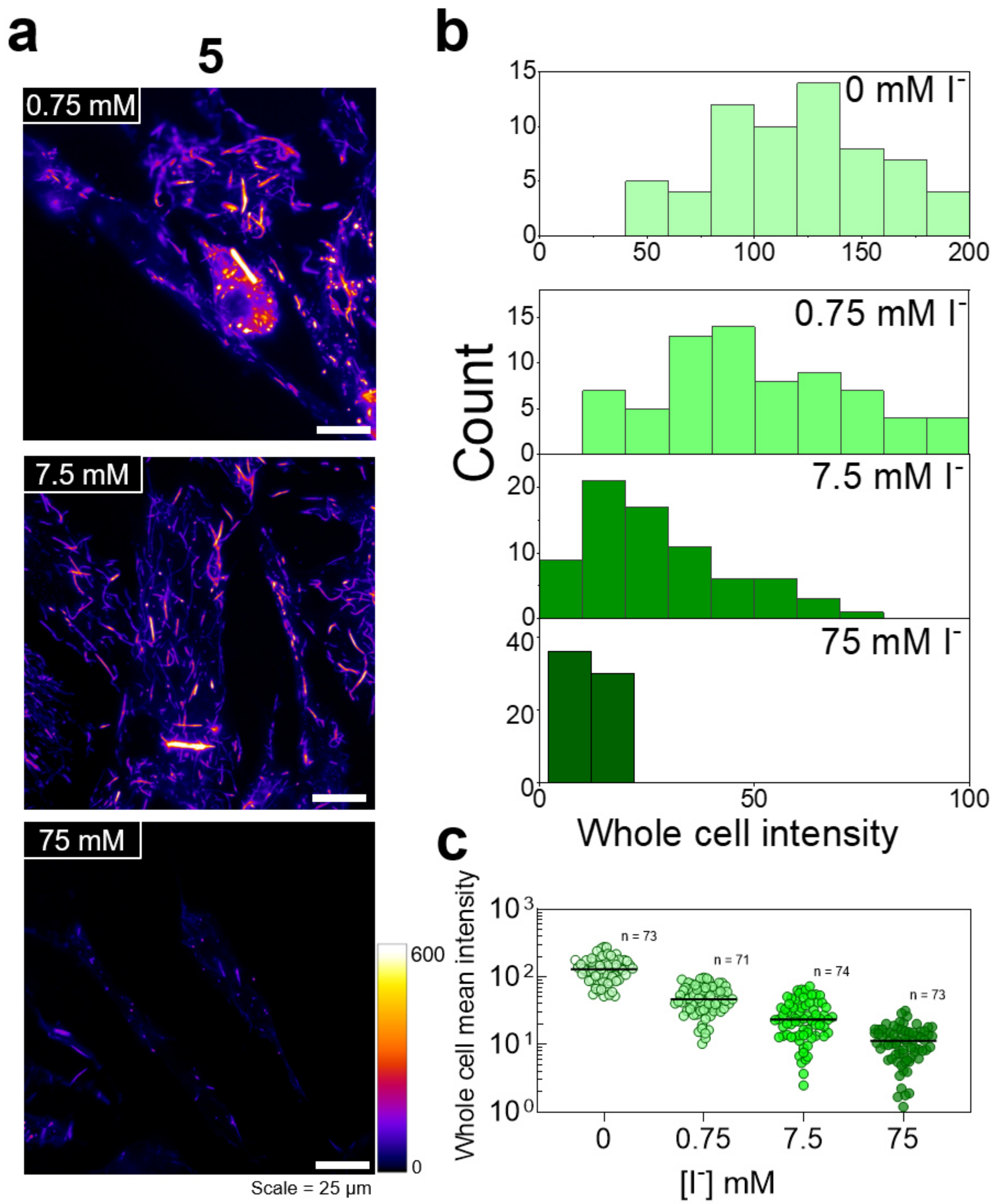


**Figure S19 Intracellular iodide sensing characteristics of 5. a,** Representative fluorescence images of **5**-labeled HDF cells upon incubation with 0 mM, 0.75 mM, 7.5 mM, and 75 mM of $I^-$. **b**,**c**, Quantification of whole cell intensity in indicated conditions. Fluorescence intensity of **5**-labeled cell decreased with increasing [$I^-$]. Experiments were performed in triplicate (n = number of cells).

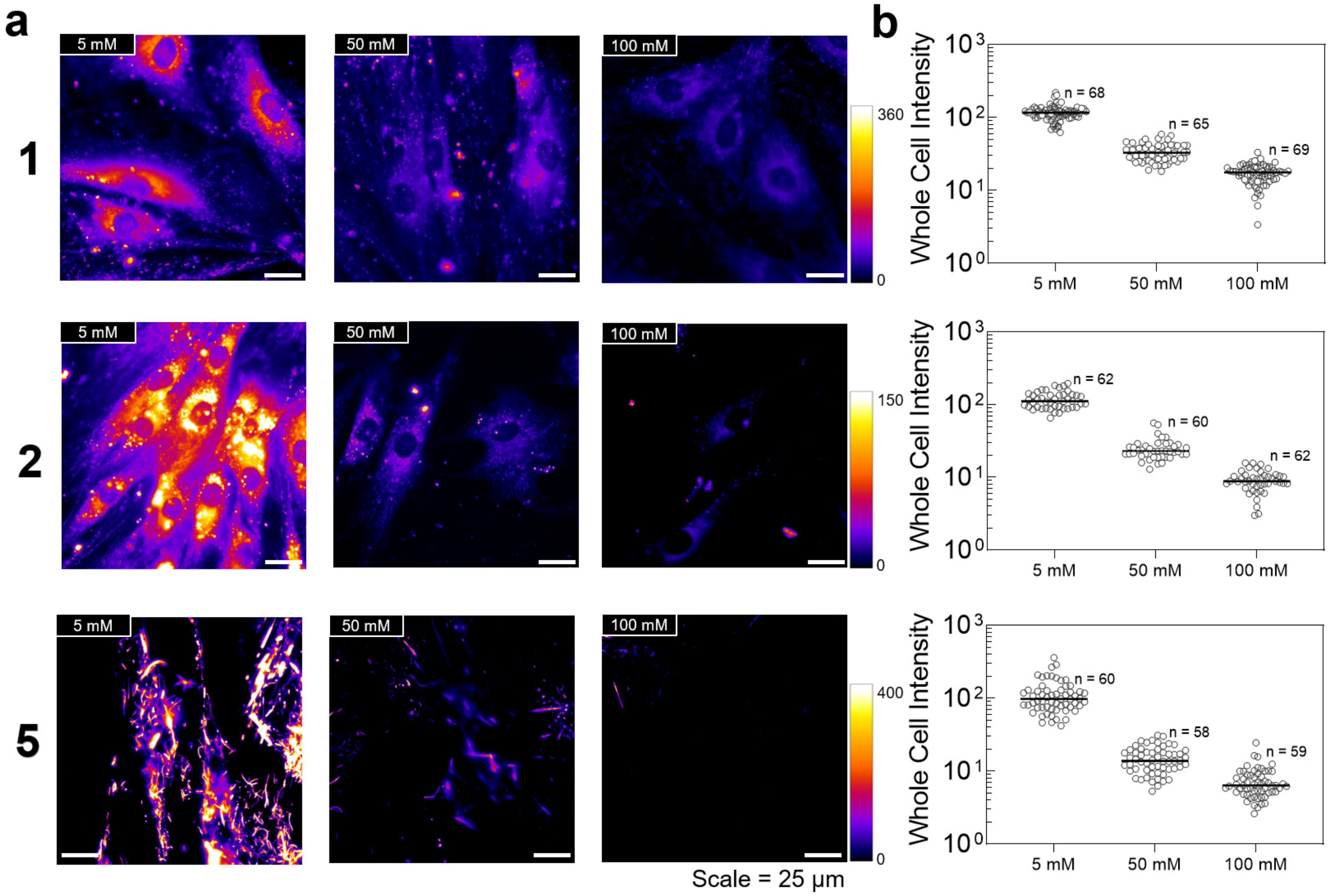


**Figure S20 Intracellular calibration of CytoCl dyes. a**, Representative fluorescence images of primary HDF cells labeled with **1**, **2**, or **5**, and clamped at the indicated [Cl⁻]. **b**, Quantification of whole cell intensity at each [Cl⁻]. Experiments were performed in triplicate (n = number of cells).

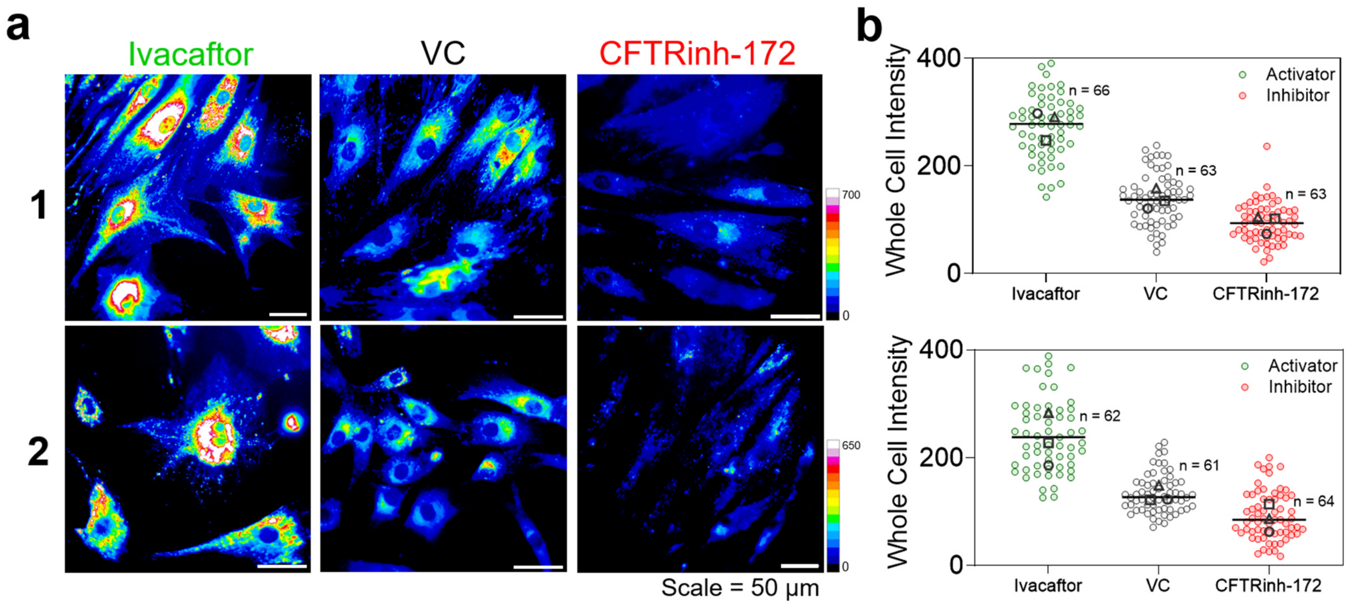


**Figure S21 CytoCl monitors the cytosolic chloride change induced by CFTR modulator. a,** Representative fluorescence images of **1**- or **2**-labeled HDF cells pre-treated with vehicle control (DMSO), 2 μM ivacaftor (CFTR activator), and 20 μM CFTRinh-172 (CFTR inhibitor). **b**, Quantification of whole cell intensity at each condition. Experiments were performed in triplicate. The median value of each trial is given by a square, circle and triangle symbol (n = number of cells).

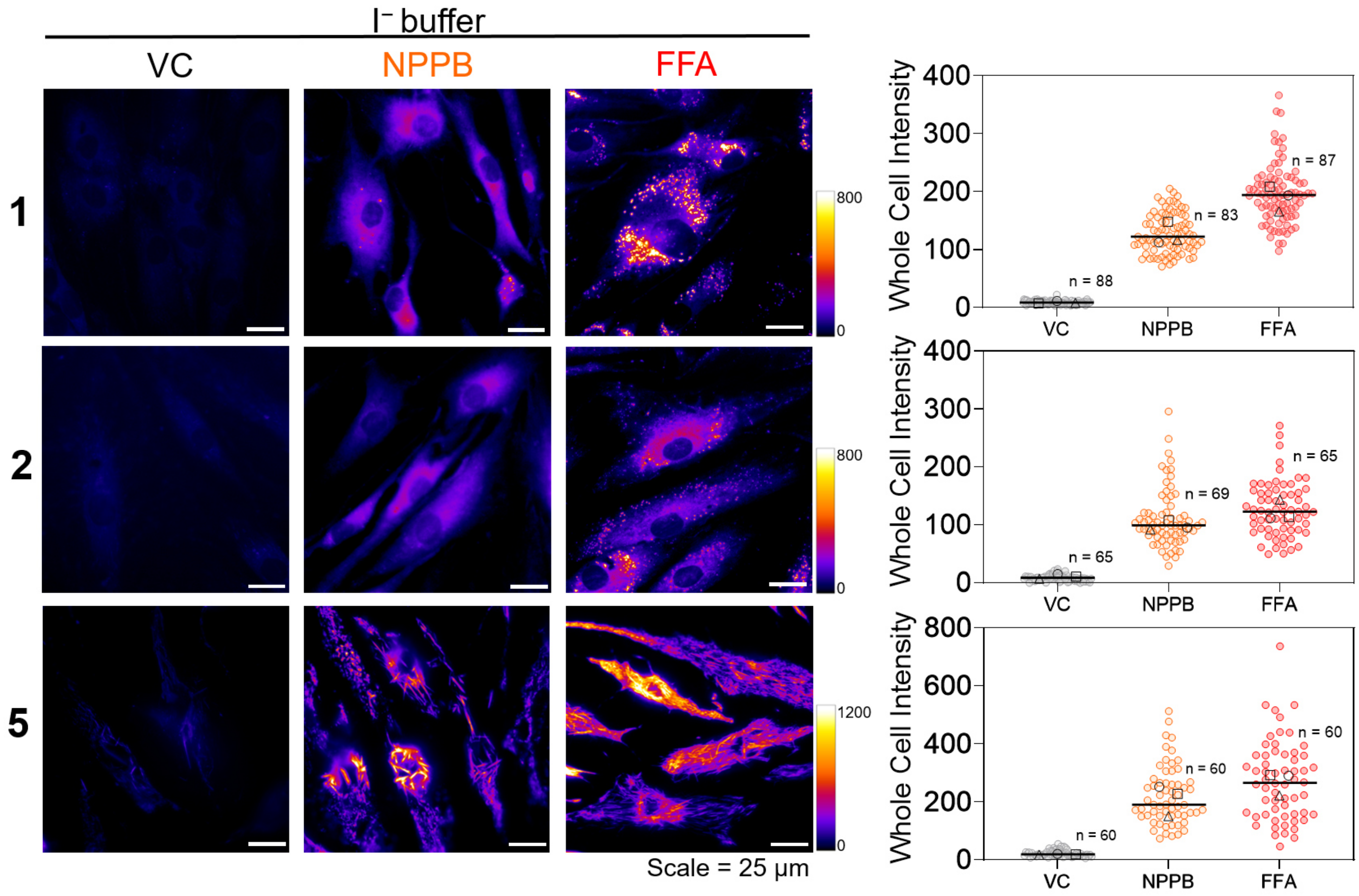


**Figure S22 Visualizing the impact of chloride channel blockers on iodide import.** Primary HDF cells pre-treated with vehicle control (DMSO), 100 µM NPPB, and 150 µM FFA were labeled with **1**, **2**, or **5**, and then subjected to iodide stimulation for 30 min. **a**, Representative fluorescence images of **1**-, **2**- and **5**-labeled HDF cells. **b**, Quantification of whole cell intensity of labeled cells at each condition. Experiments were performed in triplicate. The median value of each trial is given by a square, circle and triangle symbol (n = number of cells).

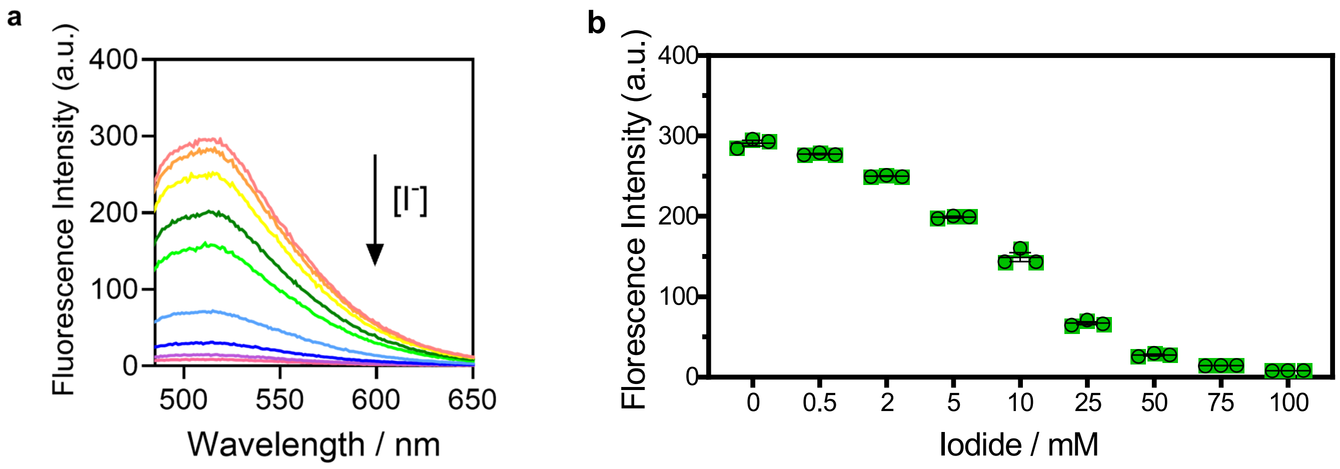


**Figure S23 Dose-dependent response of 1 to iodide ions. a,** Emission spectra and **b,** fluorescence intensity of **1** in UB4 buffer (pH = 7.0) with increasing concentrations of NaI (0-100 mM). Error bars indicate the mean ± standard error of the mean (s.e.m.) of three independent measurements.

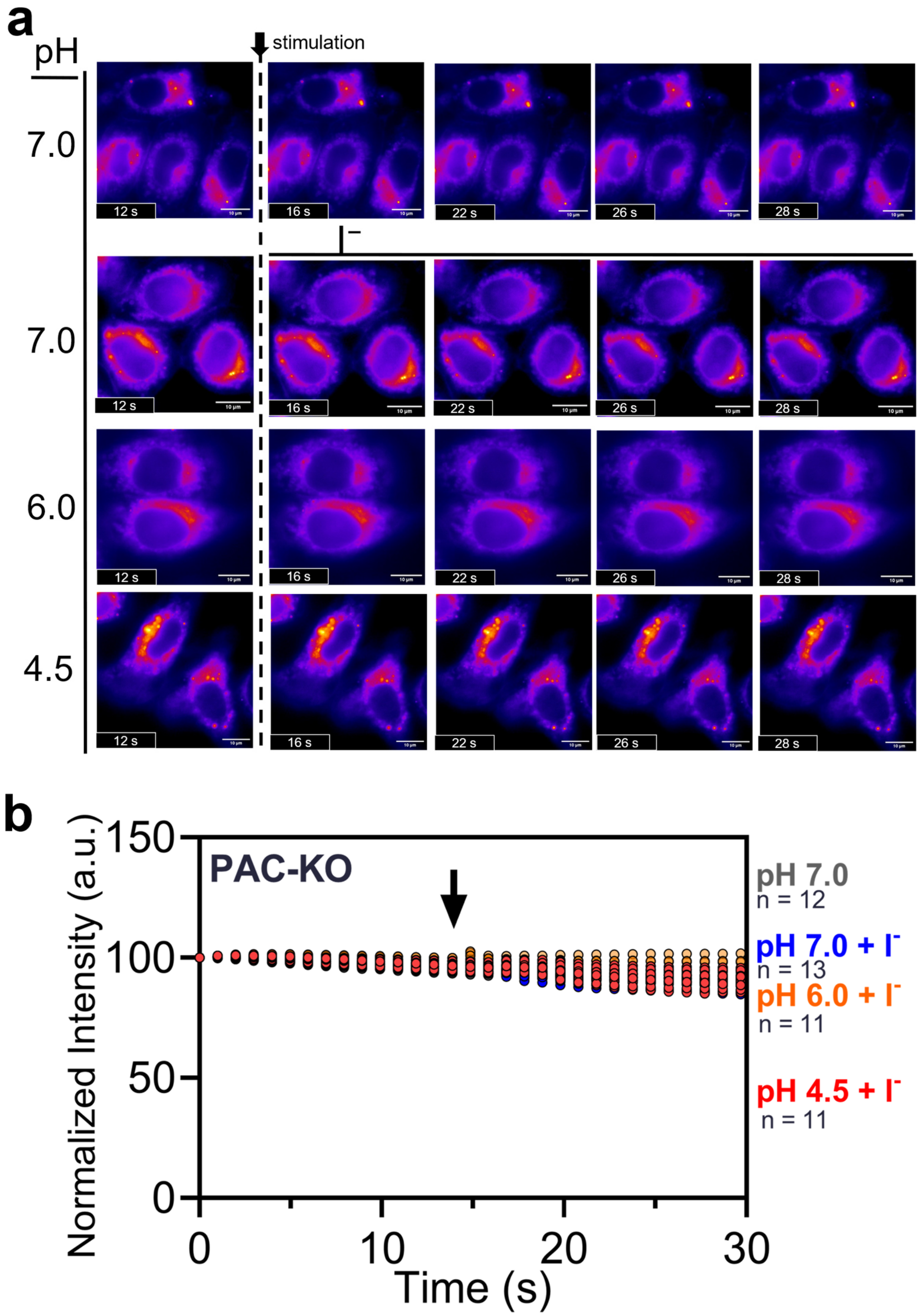


**Figure S24 No acid-induced ion flux was observed in PAC-KO cells.** **1**-labeled PAC-KO HeLa cells were stimulated with $I^-$ containing buffer at various pH. **a** Representative fluorescence images of **1**-labeled PAC-KO HeLa cells at different time points. **b,** Normalized whole cell intensity of **1**-labeled PAC-KO HeLa cells at different time points. Fluorescence intensity at different time points were normalized to the intensity at time = 0 s. The images were recorded at a frame rate of 1 fps. The dark arrow indicates the time of stimulation. Experiments were performed in triplicate (n = number of cells).
